# Structural Basis of Mammalian Respiratory Complex I Inhibition by Medicinal Biguanides

**DOI:** 10.1101/2022.08.09.503333

**Authors:** Hannah R. Bridges, James N. Blaza, Zhan Yin, Injae Chung, Michael N. Pollak, Judy Hirst

## Abstract

The molecular mode of action of metformin, a biguanide used widely in the treatment of diabetes, is incompletely characterized. Here we define the inhibitory drug-target interaction(s) of a model biguanide with mammalian respiratory complex I by combining cryo-electron microscopy and enzyme kinetics. We explain the unique selectivity of biguanide binding to different enzyme states. The primary inhibitory site is in an amphipathic region of the quinone-binding channel and an additional binding site is in a pocket on the intermembrane space side of the enzyme. An independent local chaotropic interaction, not previously described for any drug, displaces a portion of a key helix in the membrane domain. Our data provide a structural basis for biguanide action and enable rational design of novel medicinal biguanides.

**One-Sentence Summary:** Biguanides inhibit complex I by binding in the quinone channel, and exert an independent localized chaotropic effect.

## Main Text

The biguanide metformin is central to the treatment of millions of type-2 diabetes patients worldwide (*1*) and has been studied intensely in recent years for treatment of other conditions, including ischemia-reperfusion injury (*2, 3*), fibrosis (*4*), viral infections (*5*) and cancer (*6*). Optimization of biguanides for novel indications has been hindered by incomplete understanding of their molecular pharmacology and although preclinical evidence for the antineoplastic action of metformin was sufficient to justify dozens of clinical trials (*7*), the results have been disappointing (*8, 9*). Biguanides have been reported to target many cellular proteins including mitochondrial glycerophosphate dehydrogenase (*10*), presenilin enhancer 2 (*11*), F_1_F_o_ ATP synthase (*12*), cytochrome *c* oxidase (*13*), and the chloride intracellular channel 1 (*14*). Notably, several studies have described biguanide inhibition of mitochondrial respiratory complex I (proton-translocating NADH:ubiquinone oxidoreductase) (*12, 15*–*18*) supporting a mode of biguanide action in which decreased production of ATP from oxidative phosphorylation triggers the activation of AMP kinase and inhibition of adenylate cyclase, leading to beneficial downstream effects on gluconeogenic enzymes (in diabetes) and mTOR (in cancer and antiviral treatments) (*1, 6, 18, 19*).

Complex I is a 1 MDa multi-protein assembly that is central to mitochondrial and cellular metabolism. It oxidizes the NADH produced by oxidation of carbohydrates and lipids to maintain the redox state of the mitochondrial NAD^+^ pool, reduces ubiquinone-10 to drive the respiratory chain and oxygen consumption, and pumps protons out of the mitochondrial matrix. This proton pumping contributes to the proton-motive force that drives ATP synthesis through oxidative phosphorylation (*20*). Cryo-electron microscopy (cryo-EM) studies of complex I have revolutionized our understanding of its structure, mechanism and regulation, informing on redox catalysis in the hydrophilic domain, proton translocation across the membrane, and possible mechanisms of coupling between ubiquinone-10 reduction and proton translocation (*21, 22*). Furthermore, cryo-EM has discriminated different resting states of the enzyme (*22*–*25*) based on domain-level reorientations linked to altered conformational states of the quinone-binding channel (Q-channel): the ‘active’ state, with a structurally ordered, turnover-ready Q-channel, and the pronounced ‘deactive’ state, with a locally disordered Q-channel that requires restructuring and reactivation for catalysis. Biguanides bind with an unusual preference to the deactive state of the enzyme (*16*), but their binding site(s) and modes of interaction are unknown: they are expected to bind in a site downstream of the Fe-S clusters (*12*) but do not inhibit in a simple competitive manner. The interaction site is expected to be amphipathic on the basis of the biguanide positive charge and a strong correlation between inhibitory potency/cytotoxicity and hydrophobicity (*12, 26*).

Here we use cryo-EM to reveal the molecular interactions of biguanides with mammalian respiratory complex I, defining how they inhibit catalysis. We identify distinctive binding modes and rationalize biguanide protein-state selectivity to enable future implementation of structure-based drug design in the development of biguanide-based therapies for diverse applications.

### IM1761092 as a model biguanide for structural studies

To investigate the complex I binding site(s) of biguanides, we selected IM1761092 (*27*) (hereafter IM1092), a more hydrophobic (log*P* 2.37) derivative of the metformin-related antidiabetic biguanide phenformin (log*P* 0.34) that contains a 3-chloro-4-iodo-phenyl ring (Fig. 1A) and is a stronger inhibitor of complex I catalysis in bovine heart mitochondrial membranes than phenformin or metformin (IC_50_ more than 10 and 2,000 times lower, respectively, depending on the conditions, Fig. 1B & C). Its behavior is thus consistent with the reported correlation between biguanide inhibitory potency (IC_50_) and hydrophobicity (log*P*) (*28*). Use of a tighter-binding synthetic biguanide with representative biochemical behavior mitigated technical risks from using high mM concentrations of weaker binding biguanides, including excessive adventitious binding, and difficulty identifying biguanide cryo-EM densities.

**Fig. 1.**
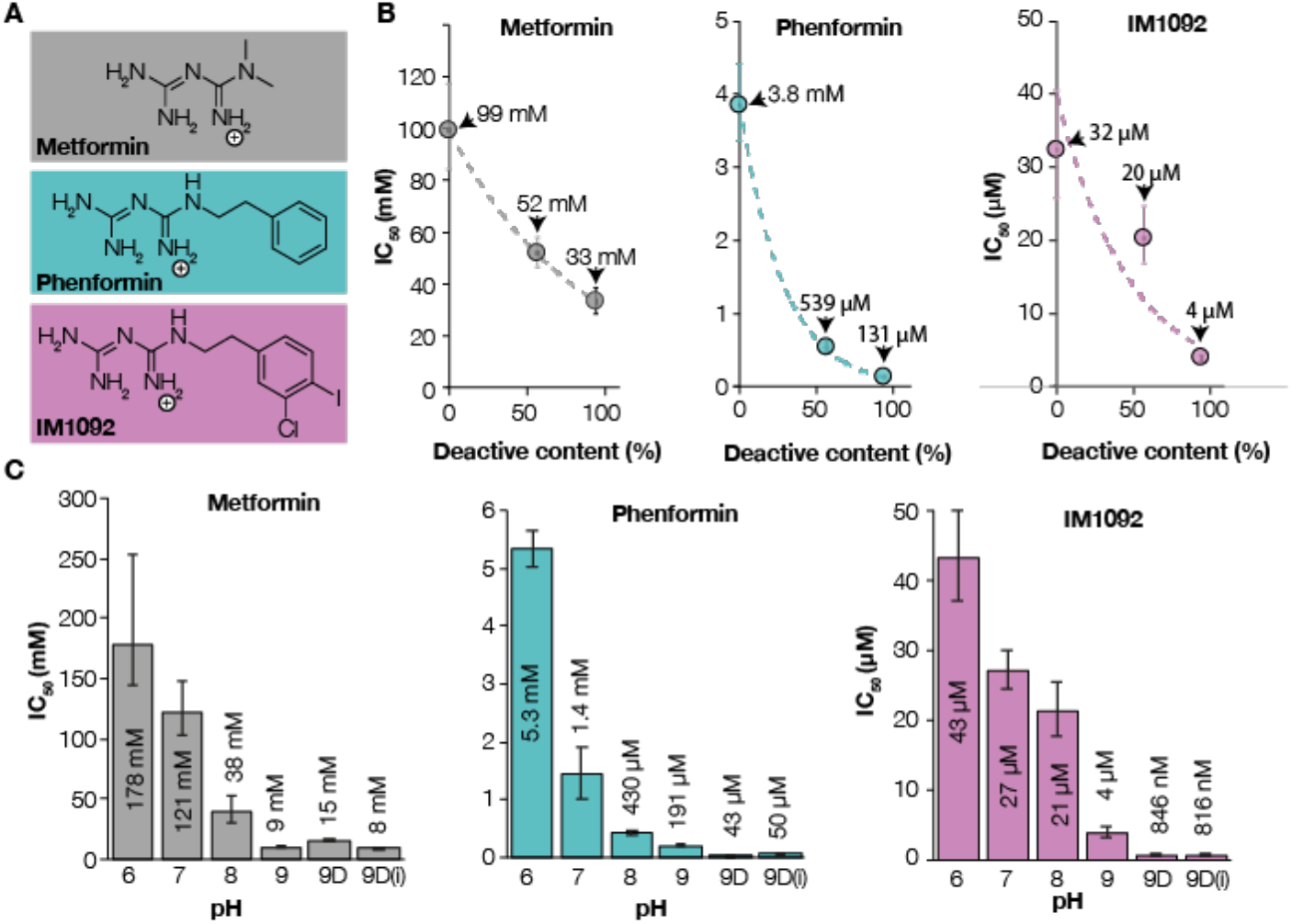
Characterization of biguanide effects on catalysis and regions of structural interest. A) Chemical structures of metformin, phenformin and IM1092 in monoprotonated form. B) Correlation between membrane deactive complex I content and IC_50_ for metformin (gray), phenformin (teal) and IM1092 (orchid). Error bars represent S.E.M for deactive content and 95% confidence intervals for IC_50_. Data are fit to an exponential regression for visualization. C) Effect of pH on IC_50_ in bovine heart membranes for metformin (gray), phenformin (teal) and IM1092 (orchid). 9D: pH 9 deactivated membranes; 9D(i): pH 9 deactivated membranes measured in the presence of antimycin A to inhibit complex III and the alternative oxidase (AOX) to oxidize quinol by a different route, confirming inhibition is on complex I. Error bars represent 95% confidence intervals.

Mammalian mitochondrial membranes ‘as-prepared’ contain a mixture of active and deactive complex I. In the deactive state, an important structural feature of the Q-channel, the loop between TMH1 and 2 in subunit ND3 (ND3 TMH1-2 loop) that carries Cys39 is disordered (*24, 25*), but in the active state, it is ordered and Cys39 is buried (*23, 25, 29, 30*). The two states can be discriminated biochemically by their sensitivity to *N*-ethyl maleimide (NEM), which derivatizes ND3-Cys39 in the deactive state, preventing catalysis, but leaves the active state unaffected. We found biguanide inhibition depends on the amount of the deactive state present in membranes (Fig. 1B). ‘As-prepared’ membranes incubated with both 200 μM IM1092 (10 x IC_50_) and NEM exhibited essentially the same deactive content as the biguanide-free control (55.9 ± 0.9% NEM-sensitive *vs*. 56.7 ± 1.0%, n = 3), indicating that biguanides do not shift the deactive/active population equilibrium. Furthermore, biguanide inhibition is stronger at higher pH (Fig. 1C). As biguanides (pKa ∼11 (*31*)) remain singly protonated at all pHs tested, the pH dependence likely arises from changes in the protein, such as in local charges on residue sidechains or phospholipid headgroups, or from conformational changes.

To inform on the selectivity of biguanides for different states, IM1092 was added to the mixed population of states in the purified resting enzyme, without purposeful deactivation for cryo-EM. No substrates were added because inhibition is weaker when biguanides are added during catalysis, rather than before (*12*). IM1092 has an IC_50_ value of 86 μM for catalysis of detergent-solubilized, purified bovine complex I (Fig. S1A) and cryo-EM grid conditions with 350 µM IM1092 were chosen to maximize binding within the limits of pH and biguanide-induced protein aggregation at high concentrations (Fig. S1). When complex I was incubated in the same conditions as for cryo-EM grid preparation then diluted into inhibitor-free assay buffer, 98 ± 5% (n = 3) of the control catalytic rate was observed, demonstrating full reversibility of inhibition under this condition.

### Overview of Cryo-EM particle populations

17,203 micrographs collected from a single cryo-EM grid yielded 598,287 good particles, which refined to an estimated global resolution of 2.1 Å (Fig. S2), but which exhibited such a high degree of heterogeneity that it was not possible to model parts of the map. Subsequent global classification yielded three major classes resembling the active and deactive states mentioned above and a state called slack (see Supplementary Text), which is of unknown functional relevance but has been described previously in cryo-EM studies of bovine complex I (*22, 24, 25*). Inhibitor-free reference maps of bovine complex I from separate inhibitor-free preparations in detergent (Fig. S3 & S4 and EMD-3731(*24*)) were used for comparison to identify and evaluate novel features found in the maps with IM1092 present. Three regions of interest are: i) densities occupying the Q-channel; ii) unusually poor density in a portion of the C-terminal lateral helix in subunit ND5 and the adjacent subunit NDUFB4; and iii) altered density at the expected position of the NDUFC2 subunit N-terminus. Different classification strategies were tested to disentangle the different states and IM1092 interactions, leading to implementation of a ‘local-first’ classification regime (see Supplementary Text) used in two separate schemes to describe 12 distinct classes (Table 1, Figs. S5 and S6 & S13).

**Table 1.** Key characteristics of classes presented in this work. White, Inhibitor-free; grey, Q-site classification scheme (Fig. S5); green, lateral helix classification scheme (Fig. S6); Unidentified, the presence of density of unknown non-protein origin; Unclear, unclear binding position, but confident identity.

| Model | PDB | Features |  |  |
| --- | --- | --- | --- | --- |
|  |  | Q-site<br>(C-C mask) | ND5/NDUFB4<br>interface | ND2/NDUFB5/NDUFA11<br>(C-C mask) |
| Active inhibitor-free | 7QSD | No density | Intact | NDUFC2-N terminus |
| Active-1092-i | 7R41 | Unidentified | Mixed | IM1092-bound (0.49) |
| Active-1092-ii | 7R42 | Unidentified | Intact | IM1092-bound (0.52) |
| Active-1092-iii | 7R43 | Unidentified | Displaced | Unidentified |
| Active-1092-iv | 7R44 | Unidentified | Disordered | IM1092-bound (0.63) |
| Deactive-1092-i | 7R45 | IM1092-bound (0.66) | Mixed | IM1092-bound (0.60) |
| Deactive-1092-ii | 7R46 | IM1092-bound (0.72) | Mixed | Unidentified |
| Deactive-1092-iii | 7R47 | IM1092-bound (0.68) | Mixed | IM1092-bound (0.68) |
| Deactive-1092-iv | 7R48 | IM1092-bound (0.66) | Intact | IM1092-bound (0.65) |
| Deactive-1092-v | 7R4C | IM1092-bound (0.71) | Displaced | IM1092-bound (0.64) |
| Deactive-1092-vi | 7R4D | IM1092 Unclear | Disordered | IM1092-bound (0.67) |
| Slack-1092-i | 7R4F | IM1092 Unclear | Mixed | IM1092-bound (0.65) |
| Slack-1092-ii | 7R4G | IM1092-bound (0.64) | Mixed | IM1092-bound (0.64) |

### Biguanide-binding site 1: ubiquinone-binding channel

Density was observed in the Q-channel, close to its entrance from the membrane, in all three major classes. Using local-first classification (Fig. S5), the particles were separated into one active class (Active-1092-i, Fig. S8), three deactive classes (Deactive-1092-i, ii and iii, Figs. S9-11), and two slack classes (Slack-1092-i and ii, Figs. S12-13 and Table 1). The major classes were assigned by global comparisons to reference maps and key local features (Table S1 and Supplementary Text). Densities for IM1092 in the Q-channel are clear in the Deactive-1092-i, ii and iii and Slack-1092-ii maps (Fig. 2C, D, E, H) and the cross-correlation fits (C-C_mask_) (*32*) for the fit of the IM1092 molecule into its density were high (0.66-0.72) (Table 1). In Slack-1092-i, while the density is consistent for the chloro-iodo-phenyl moiety, the biguanide moiety of the putative IM1092 molecule was insufficiently resolved to confidently model its orientation (Fig. 2G). The density in the Active-1092-i map occupies a similar position to the densities observed in the other classes (Fig. 2F), but a smaller additional density feature is also observed further into the Q-channel so that, together, they resemble density observed in the Q-channel of the active-apo (inhibitor-free) class of bovine complex I in nanodiscs (EMD-14133) (*22*). With an IM1092 molecule refined in different orientations into the density in the Active-1092-i map near the Q-channel entrance, C-C_mask_ values are low (< 0.5) and the shape of the density was visibly not a good fit; the second density was too small and featureless to ascertain its origin and the identity of these two densities remain unconfirmed. Overall, ∼45-60% of the total population, and ∼56-75 % of the [deactive and slack] population presents clear evidence of IM1092 occupying the Q-channel.

**Fig. 2.**
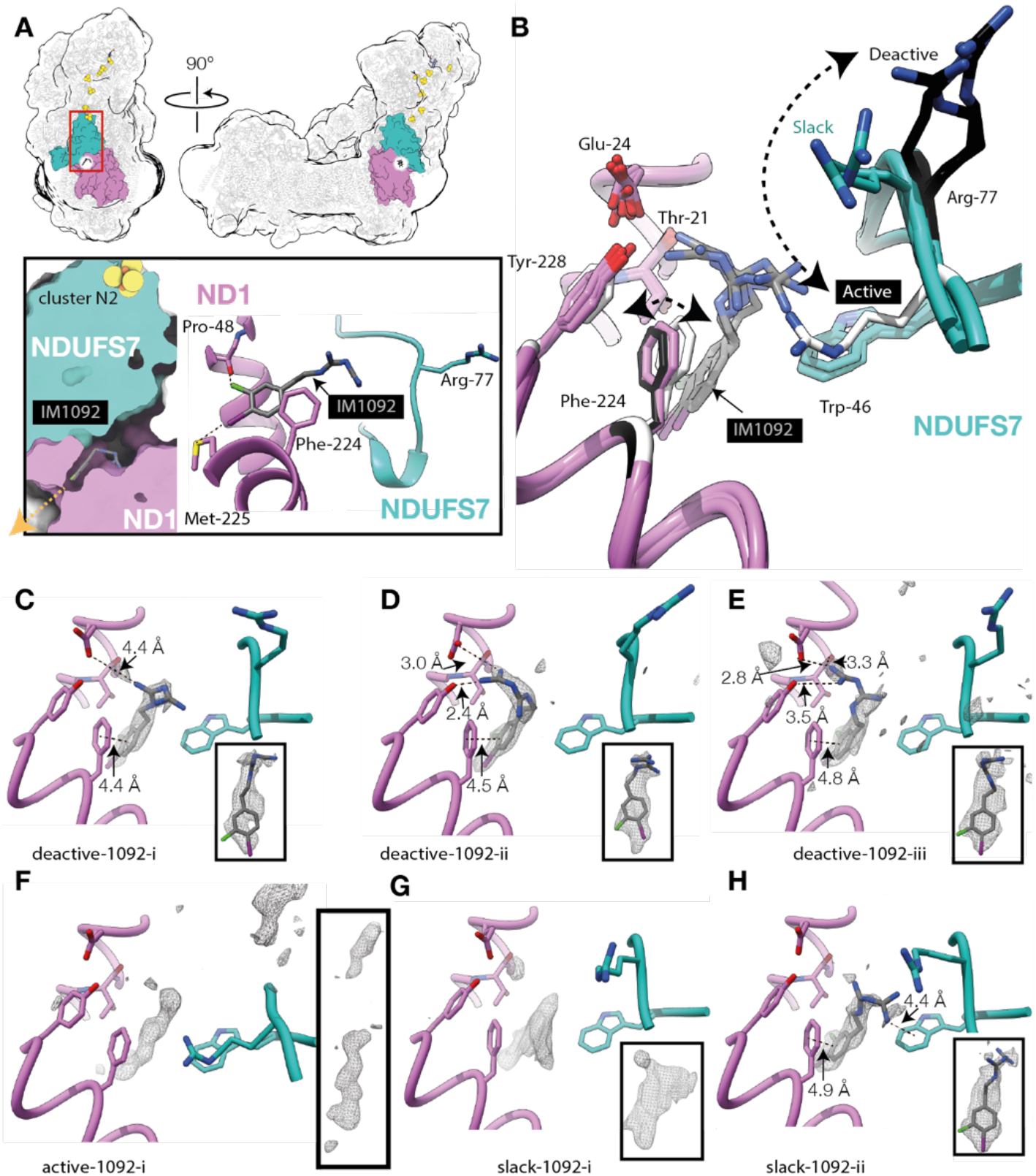
Binding of IM1092 in the Q-channel. A) Overview showing the location of the biguanide binding site and inset showing a closer view and bonding interactions of the chloro-iodo-phenyl group for Deactive-1092-i. Orange arrow shows the route for exit from the Q-channel into the lipid bilayer. B) Overlay of models for the Active-1092-i, Deactive-1092-i, ii and iii, and Slack-1092-i and ii states, aligned to subunit ND1, showing the location and variability of biguanide binding orientations, and relative position of NDUFS7-Arg77. NDUFS7-Arg77 and ND1-Phe224 are white in the active models, mint or orchid in the slack models and black in the deactive models. C-H) Cryo-EM difference map densities (composite vs models) for biguanides bound to C) Deactive-1092-i, D) Deactive-1092-ii, E) Deactive-1092-iii, F) Active-1092-i, G) Slack-1092-i and H) Slack-1092-ii. The insets show the difference map density for the biguanide for each model shown. Biguanides are not modelled in F and G due to uncertainties in the ligand identity or orientation.

In the deactive and slack states, IM1092 binds in an amphipathic site straddling two zones of the Q-channel: the hydrophobic region next to the exit (Fig. 2A) and the charged central region of the channel (*33*). The chloro-iodo-phenyl group of IM1092 points towards the channel exit (Fig. 2A), forming weak halogen bonds from the Cl to the ND1-Pro48 carbonyl and from the I to the sulfur of ND1-Met225, as well as van der Waals interactions with ND1-Phe224, ND1-Phe220, ND1-Leu55 and NDUFS7-Trp46. The biguanide moiety faces into the channel, towards the charged region, and adopts a range of different orientations (Fig. 2 C, D, E, H). In the Deactive-1092-i and Slack-1092-ii states it forms a cation-π interaction with NDUFS7-Trp46, and in the Deactive-1092-i state a weak ionic interaction with the ND1-Glu24 carboxyl also. In the Deactive-1092-ii and -iii states its orientation brings it closer to ND1-Glu24, and it forms a hydrogen bond with the ND1-Tyr228 hydroxyl. In each state with modelled biguanide, the experimental IM1092 map density at higher thresholds is consistent with a mixture of inhibitor binding poses.

Importantly, the conformation of the biguanide-binding region of the Q-channel differs between the three major states (*21*–*23*), regardless of whether an inhibitor is bound or not (Fig. 2B). Notably, NDUFS7-Arg77 swings in an arc from its position in the active state, with its sidechain pointing towards the Q-channel exit, to point approximately towards cluster N2 in the deactive state, adopting an intermediate position in the slack state (Fig. 2B). The NDUFS7-Arg77 guanidinium is ∼7 Å (slack) and 10-14 Å (deactive) away from the bound biguanide (N-N distance), but only ∼4 Å away when IM1092 is refined into the density in the Active-1092-i map in the same orientation, indicating that repulsion between their positive charges may disfavor biguanide binding in the active state.

Available PDB models for mammalian, fungal, bacterial and plant complexes I were compared to assess the conservation of key residues interacting with the biguanide moiety; NDUFS7-Trp46, ND1-Glu24, ND1-Tyr228 and NDUFS7-Arg77 are conserved in all species surveyed. The only high-quality reported human mutation in these residues in the ClinVar database (*34*) is the ND1-E24K mutation associated with LHON/MELAS overlap syndrome (*35*). Human mutations of the residues involved in the biguanide binding site are therefore very rare, consistent with their important role in enzyme function. Due to their being mitochondrial encoded and/or essential for function, these residues cannot easily be artificially mutated in a mammalian system.

### Biguanide interaction site 2: ND5 lateral helix / NDUFB4 interface

Transmembrane subunit ND5 contains an unusual long helix that runs laterally alongside ND4 and ND2, which has been proposed to either stabilize the proton-pumping modules, or act as a transmissive element in proton pumping (*36*). Substantial evidence of disorder at the ND5 lateral helix / subunit NDUFB4 interface was observed in preliminary maps, so the dataset was subject to a separate local-first classification focusing on this region (Figs. S6 & S15). The strategy yielded three major classes: one with a typical well-ordered ND5 lateral helix and NDUFB4, and two with distortion or disordering of ND5 residues 547-564 and nearby NDUFB4 residues 77-92. The classes were further separated into Active-1092-ii, iii and iv (Figs. S16-18), Deactive-1092-iv, v and vi (Figs. S19-21, Table 1), plus three slack classes that all displayed poor density for the downstream ND5 lateral helix and were not further investigated. Overall, ∼65% of the protein population was perturbed in this region, with similar proportions observed for active, deactive and slack (∼52, ∼66 and ∼58%, respectively). Deactive classes with a disrupted helix exhibit a small ‘opening’ of the angles between the membrane and hydrophilic domains, and distal and proximal membrane domains, compared to their better ordered equivalents (Fig. S22, Supplementary Text).

This interesting region of the usually well-ordered ND5 lateral helix does not form a perfect α-helix even in inhibitor-free active or deactive mammalian enzyme structures (*21*–*25*). The Active-1092-ii and Deactive-1092-iv models match the well-ordered inhibitor-free active model in this region. Two π-bulges (Fig. S15) are stabilized by interactions between the lateral helix with nearby waters, and ionic and hydrogen bonding to ND4 (Fig. 3A right).

**Fig. 3.**
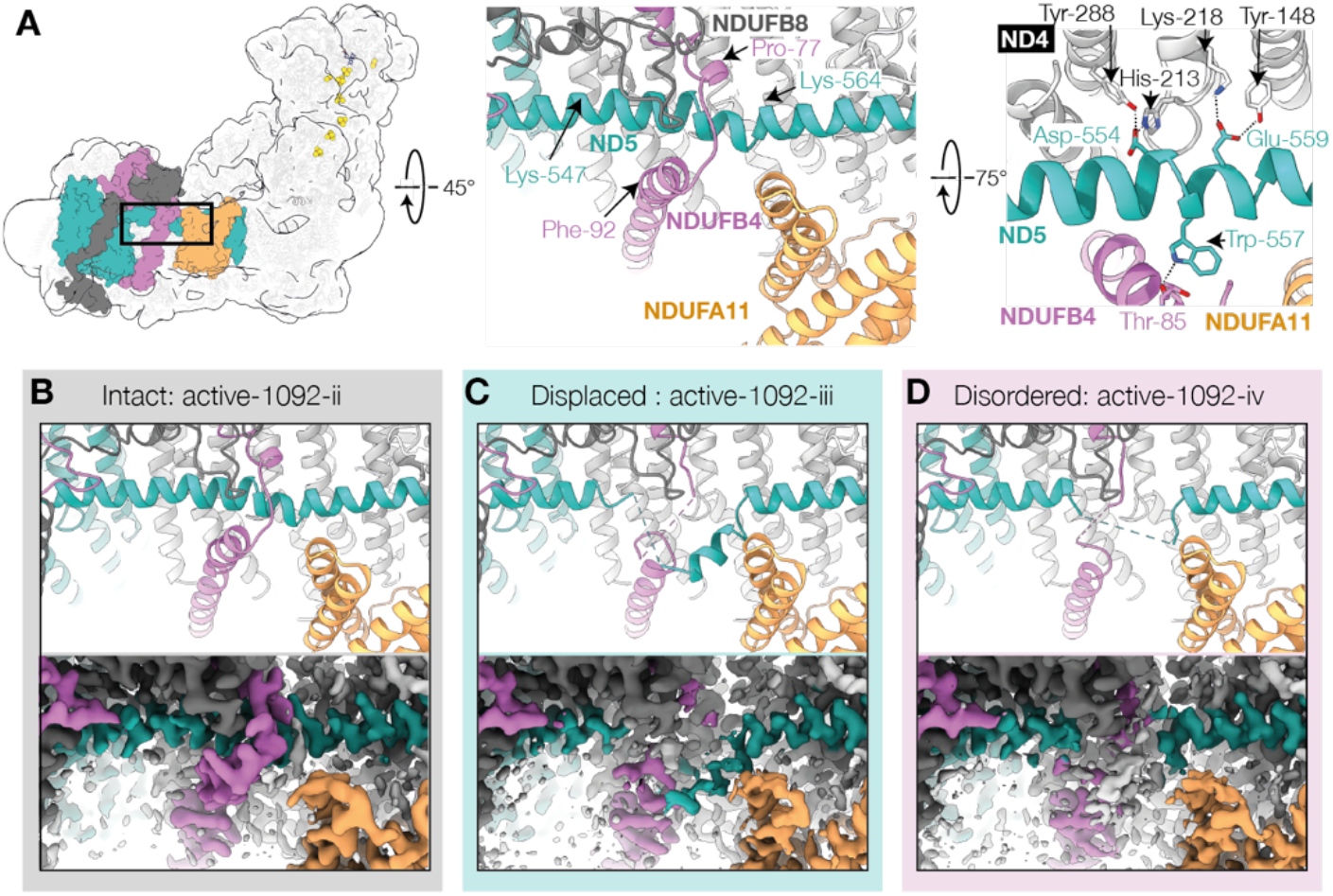
Biguanide-induced distortion and disordering of the ND5 lateral helix and NDUFB4 loop. A) Location of the structure disturbance and two views of the model of Deactive-1092-iv with hydrogen-bonding interactions indicated in black dotted lines. Orchid, NDUFB4; teal, ND5; orange, NDUFA11; dark grey, NDUFB8. B-D) Models and composite cryo-EM maps for B) Active-1092-ii, C) Active -1092-iii, D) Active-1092-iv showing progressive disordering within the series. Details of π-bulge stabilizing interactions and equivalent disordering for the deactive states (Deactive-1092-iv, v, and vi) are shown in Fig. S15.

In Active-1092-iii and Deactive-1092-v (referred to as displaced), a short portion of the ND5 lateral helix is altered: Lys547 to Ser550 are disordered, and an interruption of the helical structure at Leu562 to Pro563 allows a short stretch of helix (residues 550 to 559) to move outwards, away from the complex, and laterally, along the membrane plane (Figs. 3 and S15). A loop in NDUFB4 at residues 78-83, which usually wraps around the lateral helix, becomes disordered from Pro77 to Leu91. In Active-1092-iv and Deactive-1092-vi (referred to as disordered), the whole ND5 region from Lys547 to Lys564 appears disordered, along with the NDUFB4 loop described above (Figs. 3D & S15). Although not observed in the detergent solubilized protein, the bovine enzyme in nanodiscs (*22*) contains two phospholipids near to the distorted region. There are no density features that can be interpreted as a tightly bound biguanide nearby. Considering the positive charge on IM1092, it may interact with ND5-Asp554 and/or Glu559, thereby disrupting hydrogen bonding from the lateral helix to subunit ND4, or interact with nearby stabilizing phospholipids. Generally, π-bulges are energetically unfavorable elements (*37*), requiring stabilization by hydrogen bonding to polar sidechains or water molecules (*38*). The strained nature of this region may make it particularly prone to destabilization by guanidium-like biguanides. No mutations of Asp554 or Glu559 are observed in the ClinVar database (*34*), and acidic residues in these positions are conserved in current mammalian and plant structures, suggesting their important role in stabilizing the membrane domain.

### Biguanide-binding site 3: ND2/NDUFB5/NDUFA11

Density matching IM1092 was observed in a pocket formed by subunits ND2, NDUFB5 and NDUFA11 on the intermembrane space side of the enzyme (Fig. 4 & S23). The pocket is occupied by the N-terminal 7 residues of NDUFC2 in the inhibitor-free enzyme, both here in n-Dodecyl-B-D-maltoside (DDM) and in all states of the bovine enzyme in nanodiscs (*22*). The NDUFC2 N-terminus is displaced by the biguanide, with inhibitor densities observed in most classes here (Fig S23). The biguanide moiety is stabilized by an ionic interaction with ND2-Glu347, as well as by hydrogen bonds with the ND2-Tyr196 backbone carbonyl, ND2-Thr199 hydroxyl, ND2-Asn197 sidechain and NDUFA11-Val140 C-terminal carboxyl, as well as nearby water molecules resolved in some maps (Fig. S23). The IM1092 bound in this site does not interfere with any known catalytically relevant structural elements in the complex so is likely to represent a non-inhibitory interaction.

**Fig. 4.**
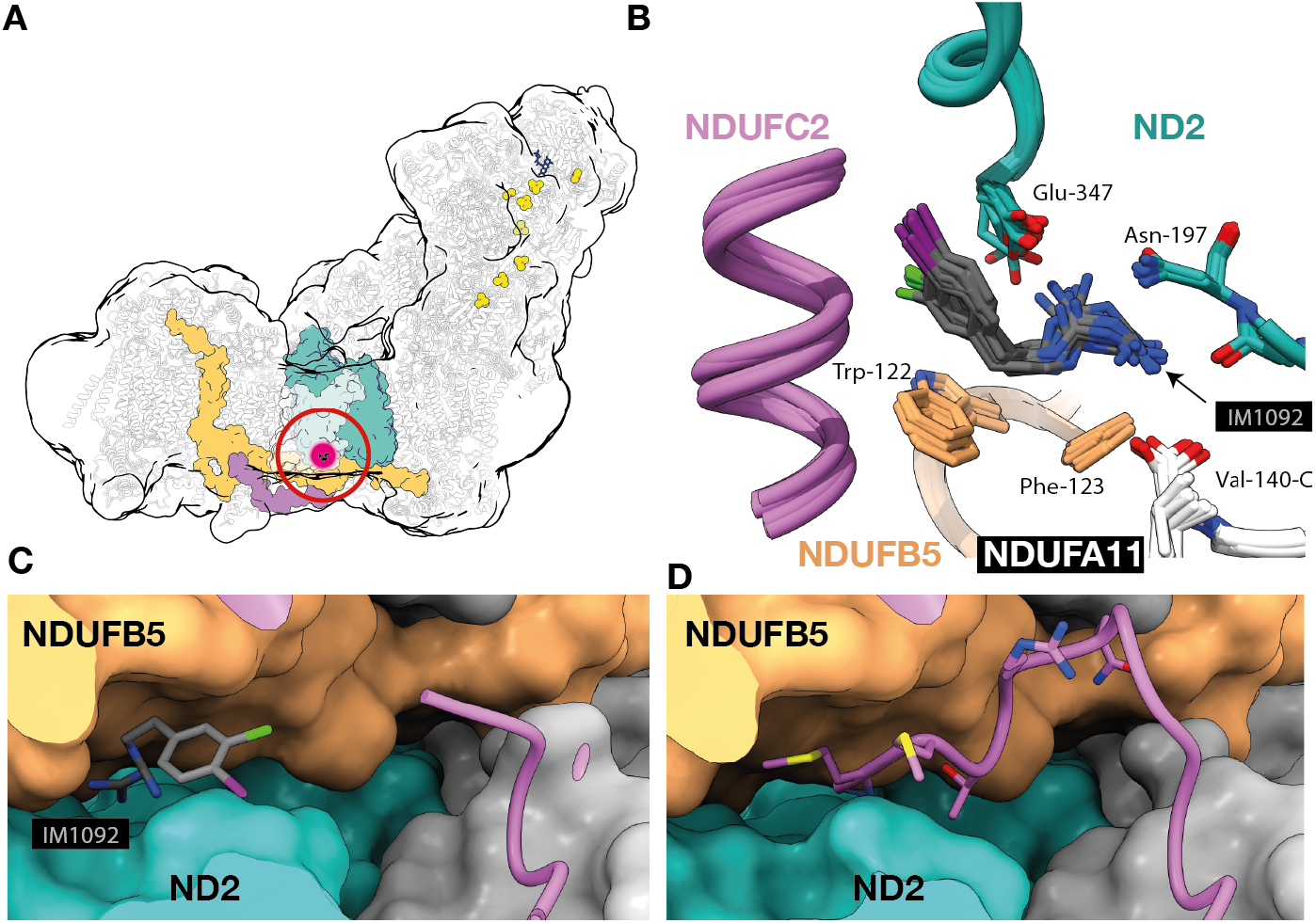
Location of the biguanide binding at the ND2/NDUFB5/NDUFA11 interface. A) Overall location of the binding site. Purple, NDUFC2; teal, ND2; orange, NDUFB5. B) An overlay of all models aligned to ND2. C) Surface representation (Active-1092-ii model) showing subunit NDUFC2 in cartoon with atoms shown for IM1092. D) The same view from the active inhibitor-free model showing the atoms from residues 1-7 of NDUFC2. Local interactions and density for the individual maps and models are shown in Fig. S8.

### Independence of interactions sites

The two local classification schemes used (Figs. S5 & S6) each yielded six models and a summary of their key features is shown in Table 1. All six classes from Q-channel classification (Fig. S5) had poor density for the ND5/NDUFB4 interface, consistent with a mixture of the three lateral helix states being represented there. Overall, ∼65% of the imaged protein population is disrupted in this region (Fig. S6), and disorder is observed in all three major classes, as well as in states with (e.g. Deactive-1092-i) or without clear binding of IM1092 at the Q-site (Active-1092-i). Furthermore, in the classes from the lateral helix classification (Fig. S6), density for IM1092 was observed in the Q-channel of deactive classes with both ordered and displaced lateral helix. Taken together, biguanide-binding in the Q-channel and the state of the lateral helix are not correlated. IM1092 is observed in the ND2/NDUFB5/NDUFA11 pocket regardless of the occupancy of the Q-channel or the status of the lateral helix (Table 1). Therefore, the three interaction sites are independent of one other.

## Discussion

### Access to the quinone-channel

Substantial inhibition of complex I *in vivo* requires the biguanide positive charge, an inherent feature of the two co-joined guanidinium moieties at physiological pH (*31*), to drive biguanide accumulation in the mitochondrial matrix (*39*) by up to 1,000-fold relative to the cytosol, in response to the mitochondrial proton-motive force (PMF). This mitochondrial concentration effect overcomes the relatively weak interactions that biguanides display with intramitochondrial targets *in vitro*, making them as potentially relevant as targets outside of the mitochondrial matrix, such as glycerophosphate dehydrogenase (*10*), which may exhibit greater intrinsic affinities. All three interaction/binding sites described in complex I contain acidic residues close to the membrane/aqueous interface, suggesting that IM1092 accesses them from the membrane, likely with the chloro-iodo-phenyl group acting as an anchor into the hydrophobic membrane core, and the hydrophilic biguanide moiety interacting with the negatively charged phosphate headgroups, as proposed previously for 1-phenyl biguanide (*40*). The most likely route of access for hydrophobic biguanides to the Q-channel is therefore via the matrix-facing phospholipid leaflet. While the Q-channel is reproducibly well-ordered in the active state, deactive (or slack) states have mobile regions of the Q-channel loops that face the mitochondrial matrix (*21*–*24, 41*), and so it is also possible that the Q-channel may become exposed to the matrix in these states, providing an alternative route for hydrophilic biguanides with poor membrane solubility, such as metformin, to enter the Q-channel.

### Major inhibitory site and selectivity for the deactive state

In the Q-channel, IM1092 binds in an amphipathic region with the biguanide moiety stabilized by hydrogen-bonding, cation-π interactions and ionic bonding, while the hydrophobic chloro-iodo-phenyl group is stabilized by weak halogen bonding and van der Waals interactions. We therefore explain the relationship between hydrophobicity and inhibitory potency (*12*) by these additional stabilizing interactions between the hydrophobic portion of the Q-binding site and hydrophobic biguanides. This is not a unique site; other neutral and highly hydrophobic ligands (DDM, cholate, rotenone and IACS-2858) have also been observed binding in overlapping sites in various protein states (*21, 22, 42*) (Fig. S24). Yet biguanides remain unique inhibitors in their state selectivity, shown here and previously (*16*). Of the other inhibitors, only rotenone has been demonstrated to bind to a deactive-like open state, but the binding was not state-specific (*21*) and the protein interactions and *in-vivo* effects of biguanides and rotenone are quite different (*43, 44*).

Despite the locations of the biguanide and ubiquinone binding sites overlapping, metformin does not display classical competitive behavior (*12*). This is likely because neutral, hydrophobic inhibitors such as piericidin A, IACS-2858, and acetogenin (*42, 45, 46*) compete for (active-like) enzyme states capable of forming the Michaelis complex, but biguanide binding occurs most readily in deactive-like states that are not pre-organized due to disordering of the Q-channel loops in NDUFS2, ND3 and ND1 (*24*). The deactive state of complex I is stabilized by high pH (*47*) and here we see a strong correlation between pH and inhibitory potency. Weak biguanide inhibition of active complex I could originate either from binding to the Q-channel, or from an inhibitory nature to the ND5 lateral helix interaction. The conformation of the NDUFS7 β1–β2 loop that contains Arg77 differs dramatically between the active, deactive and slack states. Biguanides bind 7-14 Å from Arg-77 in deactive and slack states, but the arginine guanidinium position is shifted relative to the positions in the DDM and cholate bound bovine nanodisc deactive and slack states (PDB 7QSM & 7QSO) (*22*) (Fig. S24). In all active models, Arg77 is much closer to the putative biguanide binding site (Fig. 2B), presenting a source of steric hindrance and charge repulsion that acts against strong inhibitory binding and explains the unique preference of biguanide inhibitors to inhibit NEM-sensitive (deactive) complex I (manifesting as a lower IC_50_ for deactivated complex I (Fig. 1B) (*16*)). Biguanides are less powerful inhibitors when added during catalysis, rather than before (*12*) and metformin has been proposed to act by slowing down activation (*48*) rather than by classical inhibition. Our data support an inhibitory mode that primarily acts by preventing reactivation of the resting deactive state, and we conclude that the major inhibitory interaction site of biguanides is the Q-channel.

### Local chaotropic drug-protein interactions

A key feature of biguanide interactions with complex I is the displacement and disordering of a portion of the ND5 lateral helix. This type of localized disruption has not been observed previously. Considering the similarity of biguanides to the well-known chaotrope and protein denaturant guanidinium, the biguanide may be attracted to the negatively charged ND5-Asp554 and ND5-Glu559 and specifically destabilize the hydrogen bonding networks between the lateral helix and subunit ND4. In particular, biguanide binding could weaken the interactions of the lateral helix π-bulge segment, allowing secondary structure shifts to form an α-helix and a disordered loop. Following this process, the unraveled segment of the helix would no longer provide lateral support to keep the strained junction between ND2 and ND4 (*49*) tightly together, explaining the slight ‘opening’ of the proximal and distal membrane domain interface (Fig. S22). Protein stability assays (Fig. S1) suggest that complex I is stabilized close to the IM1092 IC_50_ ranges, but destabilized at higher concentrations, and we interpret this to indicate that binding to the Q-channel (and/or ND2/NDUFB5 interface) stabilizes the protein, and chaotropic action occurs at higher concentrations. Our structures here are in a detergent micelle with tightly-bound lipids present, and the chaotropic interaction site we observe is within the phospholipid headgroup plane. Importantly, any inhibitory consequences of structural disturbance here are fully reversible, demonstrated by full recovery of activity after inhibitor dilution. We propose that biguanides such as metformin, phenformin and related drug-leads could exert similar local chaotropic actions on any number of cellular proteins, especially membrane proteins, to inhibit or stimulate their usual functions, presenting an entirely new mode of enzyme-drug interaction, and a potential explanation for the breadth of biguanide targets identified thus far.

### Implications for in-vivo mechanism of action

While other complex I inhibitor classes have been proposed as possible therapeutic compounds (*50*–*53*), biguanides appear to offer a lower toxicity profile than neutral species like rotenone, with reduced risk of Parkinsonism (*44*). This could be influenced by the ready reversibility of biguanide inhibition seen here and previously (*12*), compared to the irreversibility of very hydrophobic, neutral compounds. Negative feedback at the organelle level may be a factor: due to their charge, biguanides accumulate inside the mitochondrial matrix in response to the PMF, and as complex I becomes inhibited, proton pumping by the respiratory chain is attenuated, decreasing the PMF (*54*). Biguanide-induced inhibition of a subpopulation of complex I may therefore weaken its own inhibition in a negative feedback loop due to decreased accumulation of biguanides. *In-vivo*, the active-like state that binds canonical inhibitors is expected to be prevalent in normal tissues (*41*), whereas the opportunities for biguanides to inhibit may be restricted to the weak binding observed here against active-like states, or to the binding of putative catalytic intermediates where NDUFS7-Arg77 has moved away from the inhibitory binding site. The deactive state of complex I forms during oxygen starvation (*48*), such as during ischemia (*2*), and in solid tumor microenvironments (*55*). Discrimination of biguanides for this state means that, unlike canonical inhibitors, hypoxic tissues can be selectively targeted with less risk of also compromising mitochondrial respiration in tissues with normally functioning (active) complex I. Our data here have uncovered the strongest inhibitory binding site of biguanides as the Q-binding channel of complex I, with a further local denaturation of the ND5 lateral helix, providing a basis for future structure-based biguanide drug design for state-specific inhibition of complex I for different therapeutic applications, as well as improved prediction of additional protein target binding sites for biguanide drugs.

## Acknowledgments

We thank Dr Dima Chirgadze (Cambridge), Dr Julike Radecke and Dr Zhengyi Yang (eBIC) for their assistance with grid screening and data collection; Diamond for access and support of the cryo-EM facilities at the UK national electron Bio-Imaging Centre (eBIC), proposals BI22238-10 and EM17057-23, funded by the Wellcome Trust, MRC and BBSRC; members of the Hirst lab, especially Dr Noor Agip, for their participation in bovine mitochondrial preparations; Dr Stephen McLaughlin (LMB) for access to the nano-DSF; Dr Tom Terwilliger (Los Alamos, NM) for guidance with map sharpening and composite map generation; Dr Shujing Ding (MBU) for recording mass spectrometry data; Dr Matt Idanza (Leeds) and Dr Pavel Afanasyev (ETH, Zurich) for their scripts deposited on Github.

## Author contributions

Conceptualization H.R.B., M.N.P., J.H.

Methodology H.R.B., J.N.B.

Validation H.R.B.

Formal analysis H.R.B.

Investigation H.R.B., J.N.B., Z.Y., I.C.

Data Curation H.R.B.

Writing - Original Draft H.R.B.

Writing - Review & Editing H.R.B., J.N.B., Z.Y., I.C., M.N.P., J.H.

Visualization H.R.B.

Supervision J.H.

Project administration H.R.B., J.H.

Funding acquisition J.H.

## Data and materials availability

The structure data accession codes are EMD-14251,EMD-14252, EMD-14253, EMD-15254, EMD-15355 and PDB-7R41 (Active-1092-i), EMD-4256, EMD-14257, EMD-14258, EMD-14259, EMD-14260 and PDB-7R42 (Active-1092-ii), EMD-14261, EMD-14262, EMD-14263, EMD-14264, EMD-14265 and PDB-7R43 (Active-1092-iii) EMD-14266, EMD14267, EMD-14268, EMD-14269, EMD-14270 and PDB-7R44 (Active-1092-iv), EMD-4272, EMD-14273, EMD-14274, EMD14275, EMD14276 and PDB-7R45 (Deactive-1092-i), EMD-14277, EMD-14278, EMD14279, EMD-14280, EMD-14281 and PDB-7R46 (Deactive-1092-ii) EMD-14282, EMD-14283, EMD-14284, EMD-14285, EMD-14286 and PDB-7R47 (Deactive-1092-iii), EMD-14287, EMD-14288, EMD-14289, EMD-14290, EMD-14291 and PDB-7R48 (Deactive-1092-iv), EMDB-14292, EMD-14293, EMD-14294, EMD-14295, EMD-14296 and PDB-7R4C (Deactive-1092-v), EMDB-14297, EMD-14298, EMD-14299, EMD-14300, EMD-14301 and PDB-7R4D (Deactive-1092-vi), EMDB-14302, EMD-14304, EMD-14305, EMD-14306 and PDB-7R4F (Slack-1092-i), EMD-14307, EMD-14308, EMD-14309, EMD-14310, EMD-14311 and PDB-7R4G (Slack-1092-ii), EMBD-14127, EMD-14128, EMD-14129, EMD14130, EMD-14131 and PDB-7QSD (Inhibitor-free active) and EMDB-14126 (Inhibitor-free slack). Raw micrograph images are available at EMPIAR with accession codes EMPIAR-10991 (in presence of IM1761092) and EMPIAR-10984 (inhibitor-free). Otherwise, all data needed to evaluate the conclusions in the paper are present in the paper and/or the Supplementary Materials.

## Supplementary Materials

Materials and Methods

Supplementary Text

Figs. S1 to S24

Tables S1 to S15

References (1-25)

## References and Notes

1. M. Foretz, B. Guigas, L. Bertrand, M. Pollak, B. Viollet, Metformin: From mechanisms of action to therapies. Cell Metab. 20, 953–966 (2014).

2. K. Skemiene, E. Rekuviene, A. Jekabsone, P. Cizas, R. Morkuniene, V. Borutaite, Comparison of effects of metformin, phenformin, and inhibitors of mitochondrial complex i on mitochondrial permeability transition and ischemic brain injury. Biomolecules. 10, 1–17 (2020).

3. X. Wang, L. Yang, L. Kang, J. Li, L. Yang, J. Zhang, J. Liu, M. Zhu, Q. Zhang, Y. Shen, Z. Qi, Metformin attenuates myocardial Ischemia-reperfusion injury via Up-regulation of antioxidant enzymes. PLoS One. 12, e0182777 (2017).

4. N. Sato, N. Takasaka, M. Yoshida, K. Tsubouchi, S. Minagawa, J. Araya, N. Saito, Y. Fujita, Y. Kurita, K. Kobayashi, S. Ito, H. Hara, T. Kadota, H. Yanagisawa, M. Hashimoto, H. Utsumi, H. Wakui, J. Kojima, T. Numata, Y. Kaneko, M. Odaka, T. Morikawa, K. Nakayama, H. Kohrogi, K. Kuwano, Metformin attenuates lung fibrosis development via NOX4 suppression. Respir. Res. 17, 107 (2016).

5. S. Lehrer, Inhaled biguanides and mTOR inhibition for influenza and coronavirus (Review). World Acad. Sci. J. 2, 1–1 (2020).

6. H. Zhao, K. D. Swanson, B. Zheng, Therapeutic Repurposing of Biguanides in Cancer. Trends in Cancer. 7, 714–730 (2021).

7. M. Pollak, Overcoming Drug Development Bottlenecks With Repurposing: Repurposing biguanides to target energy metabolism for cancer treatment. Nat. Med. 20, 591–593 (2014).

8. S. Kordes, M. N. Pollak, A. H. Zwinderman, R. A. Mathôt, M. J. Weterman, A. Beeker, C. J. Punt, D. J. Richel, J. W. Wilmink, Metformin in patients with advanced pancreatic cancer: a double-blind, randomised, placebo-controlled phase 2 trial. Lancet Oncol. 16, 839–847 (2015).

9. P. J. Goodwin, W. Parulekar, K. A. Gelmon, L. E. Shepherd, J. A. Ligibel, D. L. Hershman, P. Rastogi, I. A. Mayer, T. J. Hobday, J. Lemieux, A. M. Thompson, K. I. Pritchard, T. J. Whelan, S. D. Mukherjee, H. I. Chalchal, C. D. Oja, K. S. Tonkin, V. Bernstein, B. E. Chen, V. Stambolic, Effect of metformin versus placebo on weight and metabolic factors in initial patients enrolled onto NCIC CTG MA.32, a multicenter adjuvant randomized controlled trial in early-stage breast cancer (BC). J. Clin. Oncol. 31, 1033–1033 (2013).

10. A. K. Madiraju, D. M. Erion, Y. Rahimi, X. M. Zhang, D. T. Braddock, R. A. Albright, B. J. Prigaro, J. L. Wood, S. Bhanot, M. J. MacDonald, M. J. Jurczak, J. P. Camporez, H. Y. Lee, G. W. Cline, V. T. Samuel, R. G. Kibbey, G. I. Shulman, Metformin suppresses gluconeogenesis by inhibiting mitochondrial glycerophosphate dehydrogenase. Nature. 510, 542–546 (2014).

11. T. Ma, X. Tian, B. Zhang, M. Li, Y. Wang, C. Yang, J. Wu, X. Wei, Q. Qu, Y. Yu, S. Long, J. W. Feng, C. Li, C. Zhang, C. Xie, Y. Wu, Z. Xu, J. Chen, Y. Yu, X. Huang, Y. He, L. Yao, L. Zhang, M. Zhu, W. Wang, Z. C. Wang, M. Zhang, Y. Bao, W. Jia, S. Y. Lin, Z. Ye, H. L. Piao, X. Deng, C. S. Zhang, S. C. Lin, Low-dose metformin targets the lysosomal AMPK pathway through PEN2. Nature. 603, 159–165 (2022).

12. H. R. Bridges, A. J. Y. Jones, M. N. Pollak, J. Hirst, Effects of metformin and other biguanides on oxidative phosphorylation in mitochondria. Biochem. J. 462, 475–487 (2014).

13. T. E. LaMoia, G. M. Butrico, H. A. Kalpage, L. Goedeke, B. T. Hubbard, D. F. Vatner, R. C. Gaspar, X. M. Zhang, G. W. Cline, K. Nakahara, S. Woo, A. Shimada, M. Hüttemann, G. I. Shulman, Metformin, phenformin, and galegine inhibit complex IV activity and reduce glycerol-derived gluconeogenesis. Proc. Natl. Acad. Sci. U. S. A. (2022), doi:10.1073/pnas.2122287119.

14. F. Barbieri, I. Verduci, V. Carlini, G. Zona, A. Pagano, M. Mazzanti, T. Florio, Repurposed Biguanide Drugs in Glioblastoma Exert Antiproliferative Effects via the Inhibition of Intracellular Chloride Channel 1 Activity. Front. Oncol. 9, 135 (2019).

15. M. Y. El-Mir, V. Nogueira, E. Fontaine, N. Avéret, M. Rigoulet, X. Leverve, Dimethylbiguanide inhibits cell respiration via an indirect effect targeted on the respiratory chain complex I. J. Biol. Chem. 275, 223–228 (2000).

16. S. Matsuzaki, K. M. Humphries, Selective inhibition of deactivated mitochondrial complex i by biguanides. Biochemistry. 54, 2011–2021 (2015).

17. M. R. Owen, E. Doran, A. P. Halestrap, Evidence that metformin exerts its anti-diabetic effects through inhibition of complex 1 of the mitochondrial respiratory chain. Biochem. J. 348, 607 (2000).

18. G. Rena, D. G. Hardie, E. R. Pearson, The mechanisms of action of metformin. Diabetologia. 60, 1577–1585 (2017).

19. R. A. Miller, Q. Chu, J. Xie, M. Foretz, B. Viollet, M. J. Birnbaum, Biguanides suppress hepatic glucagon signalling by decreasing production of cyclic AMP. Nature. 494, 256–260 (2013).

20. J. Hirst, Mitochondrial complex I. Annu. Rev. Biochem. 82, 551–575 (2013).

21. D. Kampjut, L. A. Sazanov, The coupling mechanism of mammalian respiratory complex I. Science (80-.). 370, eabc4209 (2020).

22. I. Chung, J. J. Wright, H. R. Bridges, B. S. Ivanov, O. Biner, C. S. Pereira, G. M. Arantes, J. Hirst, Cryo-EM structures define ubiquinone-10 binding to mitochondrial complex I and conformational transitions accompanying Q-site occupancy. Nat. Commun. 13, 2758 (2022).

23. A. N. A. Agip, J. N. Blaza, H. R. Bridges, C. Viscomi, S. Rawson, S. P. Muench, J. Hirst, Cryo-em structures of complex i from mouse heart mitochondria in two biochemically defined states. Nat. Struct. Mol. Biol. 25, 548–556 (2018).

24. J. N. Blaza, K. R. Vinothkumar, J. Hirst, Structure of the Deactive State of Mammalian Respiratory Complex I. Structure. 26, 312-319.e3 (2018).

25. J. Zhu, K. R. Vinothkumar, J. Hirst, Structure of mammalian respiratory complex i. Nature. 536, 354–358 (2016).

26. T. Sakai, Y. Matsuo, K. Okuda, K. Hirota, M. Tsuji, T. Hirayama, H. Nagasawa, Development of antitumor biguanides targeting energy metabolism and stress responses in the tumor microenvironment. Sci. Rep. 11, 4852 (2021).

27. S. W. Kim, H. W. Kim, S. H. Yoo, J. S. Lee, H. J. Heo, H. B. Lee, J. A. Kook, Y. W. Lee, M. J. Kim, W. Cho, Guanidine Compounds and use thereof US2017/0073331 A1. Mar 16 (2017), pp. 1–63.

28. H. R. Bridges, V. A. Sirviö, A. N. A. Agip, J. Hirst, Molecular features of biguanides required for targeting of mitochondrial respiratory complex I and activation of AMP-kinase. BMC Biol. 14, 65 (2016).

29. M. Babot, A. Birch, P. Labarbuta, A. Galkin, Characterisation of the active/de-active transition of mitochondrial complex i. Biochim. Biophys. Acta - Bioenerg. 1837 (2014), pp. 1083–1092.

30. E. V. Gavrikova, A. D. Vinogradov, Active/de-active state transition of the mitochondrial complex I as revealed by specific sulfhydryl group labeling. FEBS Lett. 455, 36–40 (1999).

31. D. Kathuria, A. A. Bankar, P. V. Bharatam, “What’s in a structure?” The story of biguanides. J. Mol. Struct. 1152, 61–78 (2018).

32. P. V. Afonine, B. P. Klaholz, N. W. Moriarty, B. K. Poon, O. V. Sobolev, T. C. Terwilliger, P. D. Adams, A. Urzhumtsev, New tools for the analysis and validation of cryo-EM maps and atomic models. Acta Crystallogr. Sect. D Struct. Biol. 74, 814–840 (2018).

33. J. G. Fedor, A. J. Y. Jones, A. Di Luca, V. R. I. Kaila, J. Hirst, Correlating kinetic and structural data on ubiquinone binding and reduction by respiratory complex I. Proc. Natl. Acad. Sci. U. S. A. 114, 12737–12742 (2017).

34. M. J. Landrum, J. M. Lee, M. Benson, G. R. Brown, C. Chao, S. Chitipiralla, B. Gu, J. Hart, D. Hoffman, W. Jang, K. Karapetyan, K. Katz, C. Liu, Z. Maddipatla, A. Malheiro, K. McDaniel, M. Ovetsky, G. Riley, G. Zhou, J. B. Holmes, B. L. Kattman, D. R. Maglott, ClinVar: Improving access to variant interpretations and supporting evidence. Nucleic Acids Res. 46, D1062–D1067 (2018).

35. E. L. Blakely, R. de Silva, A. King, V. Schwarzer, T. Harrower, G. Dawidek, D. M. Turnbull, R. W. Taylor, LHON/MELAS overlap syndrome associated with a mitochondrial MTND1 gene mutation. Eur. J. Hum. Genet. 13, 623–627 (2005).

36. R. G. Efremov, R. Baradaran, L. A. Sazanov, The architecture of respiratory complex I. Nature. 465, 441–445 (2010).

37. J. Ludwiczak, A. Winski, A. M. da Silva Neto, K. Szczepaniak, V. Alva, S. Dunin-Horkawicz, PiPred – a deep-learning method for prediction of π-helices in protein sequences. Sci. Rep. 9, 6888 (2019).

38. J. P. Cartailler, H. Luecke, Structural and Functional Characterization of π Bulges and Other Short Intrahelical Deformations. Structure. 12, 133–144 (2004).

39. F. Davidoff, Effects of guanidine derivatives on mitochondrial function: III. The mechanism of phenethylbiguanide accumulation and its relationship to in vitro respiratory inhibition. J. Biol. Chem. 246, 4017–4027 (1971).

40. N. Samart, C. N. Beuning, K. J. Haller, C. D. Rithner, D. C. Crans, Interaction of a Biguanide Compound with Membrane Model Interface Systems: Probing the Properties of Antimalaria and Antidiabetic Compounds. Langmuir. 30, 8697–8706 (2014).

41. Z. Yin, N. Burger, D. Kula-Alwar, D. Aksentijević, H. R. Bridges, H. A. Prag, D. N. Grba, C. Viscomi, A. M. James, A. Mottahedin, T. Krieg, M. P. Murphy, J. Hirst, Structural basis for a complex I mutation that blocks pathological ROS production. Nat. Commun. 12, 707 (2021).

42. I. Chung, R. Serreli, J. B. Cross, M. E. Di Francesco, J. R. Marszalek, J. Hirst, Cork-in-bottle mechanism of inhibitor binding to mammalian complex I. Sci. Adv. 7, eabg4000 (2021).

43. M. Konopleva, T. Yap, N. Daver, M. Mahendra, J. Zhang, C. Kamiya-Matsuoka, F. Meric-Bernstam, H. Kantarjian, F. Ravandi, M. Collins, M. Di Francesco, E. Dumbrava, S. Fu, S. Gao, J. Gay, S. Gera, J. Han, D. Hong, E. Jabbour, Z. Ju, D. Karp, A. Lodi, J. Molina, N. Baran, A. Naing, M. Ohanian, S. Pant, N. Pemmaraju, P. Bose, S. Piha-Paul, J. Rodon, C. Salguero, K. Sasaki, A. Singh, V. Subbiah, A. Tsimberidou, Q. Xu, M. Yilmaz, Q. Zhang, C. Bristow, M. Bhattacharjee, S. Tiziani, T. Heffernan, C. Vellano, P. Jones, C. Heijnen, A. Kavelaars, J. Marszalek, Targeting Oxidative Phosphorylation with a Mitochondrial Complex I Inhibitor is limited by Mechanism-based Toxicity. Res. Sq. (2022), doi:10.21203/rs.3.rs-1506700/v1.

44. S. Heinz, A. Freyberger, B. Lawrenz, L. Schladt, G. Schmuck, H. Ellinger-Ziegelbauer, Energy metabolism modulation by biguanides in comparison with rotenone in rat liver and heart. Arch. Toxicol. 93, 2603–2615 (2019).

45. H. R. Bridges, J. G. Fedor, J. N. Blaza, A. Di Luca, A. Jussupow, O. D. Jarman, J. J. Wright, A. N. A. Agip, A. P. Gamiz-Hernandez, M. M. Roessler, V. R. I. Kaila, J. Hirst, Structure of inhibitor-bound mammalian complex I. Nat. Commun. 11, 5261 (2020).

46. D. N. Grba, J. N. Blaza, H. R. Bridges, A. N. A. Agip, Z. Yin, M. Murai, H. Miyoshi, J. Hirst, Cryo-electron microscopy reveals how acetogenins inhibit mitochondrial respiratory complex I. J. Biol. Chem. 298, 101602 (2022).

47. A. B. Kotlyar, V. D. Sled, A. D. Vinogradov, Effect of Ca2+ ions on the slow active/inactive transition of the mitochondrial NADH-ubiquinone reductase. Biochim. Biophys. Acta - Bioenerg. 1098, 144–150 (1992).

48. A. Galkin, S. Moncada, Modulation of the conformational state of mitochondrial complex I as a target for therapeutic intervention. Interface Focus. 7, 20160104 (2017).

49. A. Di Luca, V. R. I. Kaila, Molecular strain in the active/deactive-transition modulates domain coupling in respiratory complex I. Biochim. Biophys. Acta - Bioenerg. 1862, 148382 (2021).

50. N. Madhusudhan, B. Hu, P. Mishra, J. F. Calva-Moreno, K. Patel, R. Boriack, J. M. Ready, D. Nijhawan, Target discovery of selective non-small-cell lung cancer toxins reveals inhibitors of mitochondrial complex i. ACS Chem. Biol. 15, 158–170 (2020).

51. J. R. Molina, Y. Sun, M. Protopopova, S. Gera, M. Bandi, C. Bristow, T. McAfoos, P. Morlacchi, J. Ackroyd, A. N. A. Agip, G. Al-Atrash, J. Asara, J. Bardenhagen, C. C. Carrillo, C. Carroll, E. Chang, S. Ciurea, J. B. Cross, B. Czako, A. Deem, N. Daver, J. F. De Groot, J. W. Dong, N. Feng, G. Gao, J. Gay, M. G. Do, J. Greer, V. Giuliani, J. Han, L. Han, V. K. Henry, J. Hirst, S. Huang, Y. Jiang, Z. Kang, T. Khor, S. Konoplev, Y. H. Lin, G. Liu, A. Lodi, T. Lofton, H. Ma, M. Mahendra, P. Matre, R. Mullinax, M. Peoples, A. Petrocchi, J. Rodriguez-Canale, R. Serreli, T. Shi, M. Smith, Y. Tabe, J. Theroff, S. Tiziani, Q. Xu, Q. Zhang, F. Muller, R. A. Depinho, C. Toniatti, G. F. Draetta, T. P. Heffernan, M. Konopleva, P. Jones, M. E. Di Francesco, J. R. Marszalek, An inhibitor of oxidative phosphorylation exploits cancer vulnerability. Nat. Med. 24, 1036–1046 (2018).

52. A. Naguib, G. Mathew, C. R. Reczek, K. Watrud, A. Ambrico, T. Herzka, I. C. Salas, M. F. Lee, N. El-Amine, W. Zheng, M. E. Di Francesco, J. R. Marszalek, D. J. Pappin, N. S. Chandel, L. C. Trotman, Mitochondrial Complex I Inhibitors Expose a Vulnerability for Selective Killing of Pten-Null Cells. Cell Rep. 23, 58–67 (2018).

53. D. A. Russell, H. R. Bridges, R. Serreli, S. L. Kidd, N. Mateu, T. J. Osberger, H. F. Sore, J. Hirst, D. R. Spring, Hydroxylated Rotenoids Selectively Inhibit the Proliferation of Prostate Cancer Cells. J. Nat. Prod. 83, 1829–1845 (2020).

54. Z. Drahota, E. Palenickova, R. Endlicher, M. Milerova, J. Brejchova, M. Vosahlikova, P. Svoboda, L. Kazdova, M. Kalous, Z. Cervinkova, M. Cahova, Biguanides inhibit complex I, II and IV of rat liver mitochondria and modify their functional properties. Physiol. Res. 63, 1–11 (2014).

55. A. Emami Nejad, S. Najafgholian, A. Rostami, A. Sistani, S. Shojaeifar, M. Esparvarinha, R. Nedaeinia, S. Haghjooy Javanmard, M. Taherian, M. Ahmadlou, R. Salehi, B. Sadeghi, M. Manian, The role of hypoxia in the tumor microenvironment and development of cancer stem cell: a novel approach to developing treatment. Cancer Cell Int. 21, 62 (2021).

