## Supplementary Information for "Structural Basis of Mammalian Respiratory Complex I Inhibition by Medicinal Biguanides"

### Materials and Methods

Preparation of bovine mitochondrial membranes

Fresh cow hearts were purchased from an abattoir (C. Humphreys and Sons, Chelmsford, U.K.). They were chopped into 2-inch cubes on site and immediately placed into cold buffer for transport. For the single heart used for the inhibitor-free complex I dataset the buffer (AT buffer) contained 10 mM Tris-HCl (pH 7.4 at 4 °C), 75 mM sucrose, 225 mM sorbitol, 1 mM EGTA, and 0.1% (w/v) fatty acid-free bovine serum albumin. For the two hearts used for the biguanide-inhibited dataset it contained 10 mM Tris-HCl (pH 7.55 at 20 °C), 250 mM sucrose and 0.2 mM EDTA. Mitochondria for the biguanide-inhibited dataset were prepared as described previously (*1*), whereas for the inhibitor-free dataset the tissue was minced and then blended in the Waring blender (15 s) in AT buffer (the addition of 2 M Tris base was omitted) and the centrifugation steps were modified (1,000 x *g* for 10 min, 4 °C, the supernatant filtered through muslin then re-centrifuged at 20,000 x *g* for 27 mins, 4 °C). The mitochondria were frozen for storage either as pellets or following resuspension to 10 mg-protein $mL^{-1}$ in 20 mM Tris-HCl (pH 7.55 at 20 °C), 1 mM EDTA and 10% (v/v) glycerol. To prepare membranes, mitochondria at 5 mg-protein $mL^{-1}$ in resuspension buffer were sonicated with a Qsonica (3 x 5 s bursts at 65% amplitude) then centrifuged at 74,000 x *g* for 1 hr, 4 °C (*2*) and the membranes resuspended to ~10 mg-protein $mL^{-1}$ in the same buffer. Larger scale preparations for kinetic assays started with the tissue from eight hearts in 10 mM Tris-HCl (pH 7.55 at 20 °C), 250 mM sucrose and 0.2 mM EDTA, and membranes were prepared using a Waring blender (*1*, *3*).

Preparation of bovine complex I

Complex I for cryo-EM analyses was prepared by a method adapted from that of Jones et al. (*4*). Starting from ~50 mg membrane protein, membranes were diluted in resuspension buffer, solubilized in 1% *n*-dodecyl β-D-maltoside (DDM, Glycon), centrifuged at 80,000 x *g* for 20 min and filtered (0.2 μM). The complexes were then separated by ion-exchange chromatography (Q-Sepharose, GE Healthcare, as described previously (*4*)) and complex I-containing fractions pooled and concentrated using a 50 kDa MWCO spin column (Merck Millipore). The samples were clarified using a Spin-X 0.22 μm centrifugal filter (Corning) then applied to a Superose 6 Increase 5/150 (inhibitor-free dataset) or 10/300 column (biguanide-inhibited dataset) (GE Healthcare) and eluted in 20 mM Tris-HCl (pH 7.14 at 20 °C), 150 mM NaCl and 0.05% DDM. The peak fractions for the biguanide-inhibited dataset were concentrated using a 50 kDa MWCO spin column (Merck Millipore) and the protein concentration quantified using a nanodrop UV-vis spectrophotometer ($\varepsilon_{280}$ = 0.2 $mg^{-1}$ mL); IM1092 was then added from a 5 mM stock solution in the same buffer to a final concentration of 350 μM for a ~15 min incubation at 4 °C before grid freezing. Both final protein concentrations were ~3 mg $mL^{-1}$. Complex I for enzyme kinetics was prepared using the same method but starting with ~300 mg membrane protein and a first centrifugation of 8,500 x *g* for 12 min; the purified protein was concentrated to ~20 mg $mL^{-1}$ and frozen in the presence of 30% glycerol for storage at -80 °C.

<u>Kinetic measurements</u>

$NADH:O_2$ oxidoreduction by membranes was measured at 10 µg-protein $mL^{-1}$ using 200 µM NADH and 1.5 µM horse heart cytochrome *c* (Sigma Aldrich) in aerated buffer containing 10 mM Tris-$SO_4$ (pH 7.5 at 32 °C) and 250 mM sucrose or 50 mM Bis-tris propane (pH values as indicated). For assays with deactivated membranes at pH 9, 400 µM NADH was added because a catalytic lag of ~20 mins was observed prior to achieving linear catalysis. Where appropriate, 1 µM antimycin A and 15 µg $mL^{-1}$ AOX (prepared as described previously (*5*)) were added to assays. NADH:decylubiquinone (NADH:DQ) oxidoreduction by complex I was measured at 0.5 µg-protein $mL^{-1}$ using 200 µM NADH and 200 µM DQ in buffer containing 0.075% soy bean asolectin (Avanti Polar Lipids), 0.075% CHAPS (Merck Chemicals) and 20 mM Tris-HCl (pH 7.5 at 20 °C). All assays were performed at 32 °C and initiated by addition of NADH. Rates were measured by linear regression of the maximal slope and quantified by the absorbance of NADH at 340-380 nm ($\varepsilon$ = 4.81 $mM^{-1}\cdot cm^{-1}$). IM1092 was added from DMSO stock solutions, with appropriate DMSO controls. $IC_{50}$ values observed for the NADH:DQ assay are higher than for the $NADH:O_2$ assay because the larger hydrophobic phase volume present in the NADH:DQ assay effectively dilutes the IM1092. To test the reversibility of IM1092 binding, aliquots of complex I (2.5 mg $mL^{-1}$) with or without 350 µM IM1092 were incubated at 4 °C for 15 min, then diluted into the NADH:DQ assay as described above. Preparation of deactive membranes was achieved by heating membranes at 10 mg $mL^{-1}$ at 37 °C for 20 mins in the absence of NADH. Preparation of active membranes was achieved by incubating 'as prepared' membranes at 10 mg $mL^{-1}$ at 4 °C with 2 mM *N*-ethylmaleimide (NEM) for 1 hr in the absence of NADH; residual activity is only attributed to the active population of the complex. The sensitivity of membrane activity to NEM was determined to evaluate the active/deactive ratio by incubating the membranes (2 mg-protein $mL^{-1}$) with 2 mM NEM at 4 °C for 20 mins then measuring the $NADH:O_2$ activity relative to a DMSO control. The NEM-insensitivities were 86.4 ± 7.4 and 52.0 ± 0.4% (n = 3) for the membranes used for the inhibitor-free and biguanide-inhibited datasets, respectively. $IC_{50}$ curves were fitted using log(inhibitor) vs. normalized response with variable slope in Prism 8.

<u>Nano-DSF and protein aggregation and assays</u>

Purified complex I was diluted at 4 °C to 100 µg $mL^{-1}$ or 2.5 mg $mL^{-1}$ in Tris-HCl pH corrected to either 7.14 or 7.96 at room temperature and IM1092 was added immediately prior to loading into capillaries for analysis in a Prometheus NT.48 (NanoTemper). The temperature was increased from 15 to 95 °C (3 °C $min^{-1}$). Protein stability was measured using the 350/330 nm ratio and protein aggregation was measured in back-reflection mode as scattering. In all cases, due to complexity in the traces, the non-derivatized data was used to discern an overall melting temperature ($T_m$) as the temperature at which the protein was overall 50% aggregated or denatured, by fitting to a Boltzmann sigmoid in Prism 8.

<u>Confirmation of protein sequences</u>

The complex I sample used for the biguanide-inhibited dataset was treated with iodoacetamide, precipitated with ethanol, then digested with chymotrypsin or trypsin in 50 mM $NH_4HCO_3$ and analyzed by LC-MS as described previously (*6*). Briefly, peptides were fractionated by liquid chromatography using a gradient of 5-40% acetonitrile in 0.1% (v/v) formic acid and analyzed on a Q-Exactive Plus Orbitrap mass spectrometer (Thermo Scientific). Data were acquired from 400 to 1600 m/z for precursor ions and the ten most abundant precursor ions fragmented by HCD in nitrogen. Peptide fragmentation data were searched against the mammalian sequences from SwissProt (January 2020) using MASCOT and allowing for missed cleavages (5 for chymotrypsin, 4 for trypsin) with error tolerant searches allowing for variable oxidation of methionine and cysteine carbamidomethylation. For subunit NDUFS2, a tryptic peptide with mass 1074.6662 Da exhibited a fragmentation pattern consistent with a Q>R substitution at position 129 compared to Uniprot sequence P17694 (Mascot score of 32, 95% confidence threshold of 15). In addition, a matching preparation from the same batch of mitochondria was analyzed by ESI for intact protein masses (*6*). A mass of 36,708.6 Da correlating to subunit NDUFA10 with a N>K substitution compared to Uniprot sequence P34942 (calculated substituted mass 36,706.88 Da) was observed, and a matching substitution at position 255 confirmed by error-tolerant searches of chymotryptic peptides (a mass 1583.8028 Da with a consistent fragmentation pattern as observed previously (*7*), Mascot score of 50, 95% confidence threshold of 26). These data explain cryo-EM density discrepancies observed at these two positions, so all models here contain Arg and Lys, respectively.

Mitochondrial DNA was extracted from 100 µL (1.8 mg-protein $mL^{-1}$) of the mitochondria used for the biguanide-inhibited dataset using a Qiagen DNeasy blood and tissue kit according to the manufacturer's instructions. The regions of the mitochondrial DNA that encode the seven ND subunits were amplified from 10 ng of template DNA by PCR (Q5 high fidelity polymerase, 30 cycles, 67.5 °C annealing temperature) using the primers `GGTGCAACCGCTATCAAAGG` and `AAGCAGCTTCAATTCTGCCG` for ND1-2 and `CGACGGAGTTTACGGCTCAA` and `AGCTCCGTTTGCGTGTATGT` for ND3-6. The products were treated with a QIAGEN PCR cleanup kit and subjected to Sanger sequencing with the following primers:

| Forward | Reverse |
|---|---|
| 5′(TGTTTCTATCTATTGATGAGGCT)3′ | 5′(GGCTGTGTGCTGTGGAGTTA)3′ |
| 5′(TTGGGTTAACTCCACAGCACA)3′ | 5′(TGACGGAGGCAGATTGAGC)3′ |
| 5′(GCTCAATCTGCCTCCGTCAA)3′ | 5′(GCAGTTCTTGCATACTTTTTCGG)3′ |
| 5′(AGGATAGTAGTTTATCCGTTGGTCT)3′ | 5′(AGGAAGATACCTGCTACCACTA)3′ |
| 5′(AACTCCCGTCTCAGCACTAC)3′ | 5′(TTTCTAGGGCTAAGATGAAGCCT)3′ |
| 5′(TCCTAGGCTTCATCTTAGCCC)3′ | 5′(GGCTGGAAGGTCGATGAATG)3′ |
| 5′(ACAACACAAAACCCTGCCCT)3′ | 5′(AAAGTTTCGTGGGGGTGGAG)3′ |
| 5′(AGGACCATTTGCCCTCTTCT)3′ | 5′(TTTGCGTACTCTGCTATGAAGAA)3′ |
| 5′(TGACTCAATCAACAGCCTCA)3′ | 5′(AGTATTGAGGCTATTGGGCT)3′ |

Sequencing data gave good coverage in both forward and reverse directions and uncovered eight nucleotide differences from the reference genome NC_006853.1 but only one led to an amino acid substitution (V411I in subunit ND5). Both the variant and reference sequence nucleotide were observed suggesting either heteroplasmy or a difference between the two cow hearts in the sample. Therefore, the reference sequence was used for structure modeling.

<u>Cryo-EM grid preparation, data acquisition and processing</u>

PEG-thiol derivatized UltrAuFoil® gold grids (R 0.6/1, Quantifoil Micro Tools GmbH) were prepared and frozen with complex I as described previously (*8*) at 4 °C, using an FEI Vitrobot IV

(Thermo Fisher) with blotting for 9.5 s at blotforce setting -10. Screening images were taken using a Talos Arctica at 200 kV at 73k x magnification, -2.5 µm defocus with a 100 µm objective aperture and 50 µm C2 aperture, and a Falcon 3 detector in linear mode. The final datasets were collected from a single grid each using a Gatan K3 detector and a BioContinuum GIF energy filter (biguanide-inhibited dataset) or a K2 detector with a Quantum GIF energy filter (inhibitor-free dataset) mounted on an FEI 300 keV Titan Krios with a 100 µm objective aperture, 70 µm C2 aperture and EPU software version 2.6 (biguanide-inhibited dataset) or 2.4 (inhibitor-free dataset) at the UK National Electron Bio-Imaging Centre (eBIC). The energy filter was operated in zero-energy-loss mode with slit width 20 eV. Biguanide-inhibited data were collected in super-resolution mode at 1.072 Å $pixel^{-1}$ (nominal magnification 81,000×) with defocus range -1.0 to -2.4 µm in 0.2 µm increments and the autofocus routine run every 10 µm. The dose rate was 23.7 electrons $Å^{-2}$ $s^{-1}$ with 1.7 s exposure captured in 40 frames (total dose ~40 electrons $Å^{-2}$). Data were acquired as one shot per hole in aberration-free image shift (AFIS) mode with 5 s delay after stage shift and 3 s delay after image shift. Inhibitor-free data were collected in counting mode at 1.056 Å $pixel^{-1}$ (nominal magnification 47,600×) with defocus range -1.5 to -3.1 µm in 0.2 µm increments and the autofocus routine run every 10 µm. The dose rate was 4.2 electrons $Å^{-2}$ $s^{-1}$ with 12 s exposures captured in 25 frames (total dose ~50 electrons $Å^{-2}$). Data were acquired as one shot per hole with 10 s delay after stage shift.

The inhibitor-free dataset was processed using RELION-3.0 (*9*) (Fig. S3). First, beam-induced movement for 2,370 micrographs was corrected using RELION's implementation of MotionCor2, both with and without dose weighting. CTF estimations were taken from non-dose weighted micrographs using GCTF-1.18 (*10*). Micrographs that contained non-vitreous ice were manually excluded. Model-based autopicking yielded a total of 56,908 particles, which were extracted from dose-weighted micrographs and CTF corrected with an amplitude contrast of 0.1. 2D classification yielded 46,507 particles, which were re-extracted and subjected to 3D classification with searches from 7.5° until 0.9°, yielding 32,668 good particles. Particles were 3D refined and subjected to Bayesian polishing with the first 12 frames, yielding a map at 3.3 Å, then classified with searches from 7.5 to 1.8° into two classes. Cycles of 3D refinement and CTF refinement (for beamtilt and per particle defocus, astigmatism and B-factor correction) produced two maps, one that resembled the previously described active state (*11*, *12*) at 3.1 Å (18,231 particles), and one with poor density for subunit NDUFA11 and a more 'open' conformation that resembled the previously described slack state (*12*) at 3.5 Å (14,437 particles). All resolutions are defined using the FSC = 0.143 criterion.

The biguanide-inhibited dataset was processed using RELION-3.1 (*9*, *13*) (Figs. S2, S5 and S6). First, beam-induced movement for 17,203 micrographs was corrected for using RELION's implementation of MotionCor2, both with and without dose weighting. CTF estimations were made from non-dose weighted micrographs using CTFFIND4 (*14*). Micrographs that contained thick or non-vitreous ice were excluded by fitting the CTF up to 4 Å (to include any diffuse water ring) and removing particles with CTF figure of merit <0.04, as well as estimated resolution >6 Å, and CTF astigmatism outliers (<20 and >500), leaving 14,931 good micrographs. Model-based autopicking using a previous bovine complex I model picked a total of 907,090 particles that were extracted with down sampling to 2.41 Å/pix from dose-weighted micrographs and CTF corrected with an amplitude contrast of 0.1. 2D classification yielded 864,847 particles, which were re-extracted at 1.072 Å/pix. The output from 3D refinement was used as the input for crude 3D classification with an angular sampling interval of 1.8° resulting in 671,489 good protein particles that refined to 2.3 Å. 3D refinement was run using --pad 1 and early-stage refinements also utilized --maxsig 2000 to limit the orientations considered and speed up the process. Particles were subject to rounds of CTF and 3D refinement to correct for anisotropic magnification, beamtilt, trefoil and higher order aberrations, as well as per-particle defocus, astigmatism, B-factor-weighting and spherical aberration. As the images were obtained using AFIS, two scripts (optics_add.py and optics_split.py, https://github.com/afanasyevp/afis) were used to separate the data into 45 optics groups to allow correction for any residual relative beamtilt differences. The data were globally classified into three classes containing 148,441 (22%), 292,296 (44%) and 230,746 (34%) particles respectively, all of which displayed a preferred orientation unusual for DDM solubilized complex I, and presumably induced by biguanide addition. Therefore, to improve particle orientation distributions, particles were filtered by rlnMaxValueProbDistribution using a script (part_angdist_eq.py, https://github.com/attamatti/orientation_equlization) with 1,000 bins and a standard deviation of 1.2. Particle motion was further corrected using Bayesian polishing during which the data were rescaled to 0.731 Å/pix. After removal of duplicates, a total of 598,297 particles were CTF-refined and refined together with solvent flattening to a resolution of 2.11 Å based on the FSC = 0.143 criterion. This map was highly fragmented in the distal membrane region because the particles comprised several distinct states (see below). Global classification back into three classes using an angular sampling of 0.47° yielded classes of 120,325 (20%), 287,942 (48%) and 190,030 (32%) particles, correlating to the previously described active, deactive and slack states (*8*, *11*, *12*).

Targeted subclassification

*i) Quinone-binding channel (Fig. S5)*

The preliminary deactive map (see above) was manually edited in Chimera (*15*) to remove density outside of the proximal membrane domain region. To generate this and subsequent masks for local classifications, RELION was used to apply a lowpass filter of 10 Å with extension of 10 pixels and a soft edge of 10 pixels, unless otherwise stated. Focused classification on the Q-channel region was performed without alignment using a T (regularization parameter) value of 100 using a mask generated from a molmap density of a crudely fitted IM1092 PDB molecule, generated in Chimera. Two major classes were identified with densities in the Q-channel, one with a density resembling IM1092 (70% of particles) and one with a long density feature (16%), plus two minor classes (8% and 5%) at lower resolution and without any obvious ligand densities occupying the Q-channel. The class containing the long density, likely endogenously-bound ubiquinone-10, was not investigated further. The particles in the major 70% class were reverted to include the density of the entire complex, followed by consensus refinements, and then global classification with angular searches at 0.47° into three classes corresponding to the active, deactive and slack states (48,367, 226,483 and 143,470 particles, respectively). The active class was not resolved further and is named the Active-1092-i state. ND6-TMH4 was poorly ordered in the slack class and a subclassification on the expected location of this helix allowed two minor classes to be discarded, although the density remained disordered. In the deactive and slack classes, the densities for subunits NDUFS2 (residues 53-60), ND3 (31-46), ND1 (62-64 and 207-216) and NDUFA9 (254-278,186-198 and 323-334) were disordered, as observed previously for deactive complex I (*8*, *11*, *12*). Therefore, these regions were subjected to further subclassification without alignment and with a T value of 100, using a mask that was generated from a molmap of these loops from the preliminary map of the active state. The approach yielded three deactive classes (Deactive-1092-i to iii) and two major slack classes, (Slack-1092-i and ii) (see Fig. S5).

*ii) ND5 lateral helix (Fig. S6)*

Particle subtraction and focused refinement of the distal membrane domain were performed in the same manner as for the proximal membrane domain (see above). This was followed by rounds of focused classification on the ND5 lateral helix without alignment, using a T value of 100. The mask for this was generated from a molmap of PDB 5O31, for the corresponding region of subunits ND5 and B15 found to be partially disordered here, generated without binary extension and with a soft edge of 5 pixels. Three classes were readily identified: one with an intact and well-ordered lateral helix and NDUFB4 loop (termed ordered), one in which the ND5 lateral helix is displaced and the NDUFB4 loop partially disordered (termed displaced), and one with disorder in both the ND5 lateral helix and NDUFB4 loop (termed disordered). The particles in each class were reverted to include the density of the entire complex, followed by consensus refinements, and then global classification with angular searches only at 0.47° to separate the active, deactive and slack populations. The classification thus yielded nine maps in total (see Fig. S6): Active-1092-ii, iii and iv and Deactive-1092-iv, v and vi (ordered, displaced and disordered, respectively), plus three slack classes, one with well-ordered NDUFB4, which were not investigated further because they all contain a poorly defined ND5 lateral helix from residue ~660 onwards.

Preparation of consensus and composite maps

To aid modeling, composite maps were generated to improve map quality in the distal membrane domain. The consensus maps were manually segmented into three sections, crudely comprising the hydrophilic, proximal membrane and distal membrane domains, followed by focused refinement. All masks used for solvent flattening, subtraction and post processing were generated using the molmap command in Chimera (*15*) from a near-complete model, low-pass filtered to 15 Å with a soft edge of 10 pixels. The final globally sharpened consensus and focused maps were corrected for the Material Transfer Function of the detector and sharpened with estimated B-factors in RELION. For the biguanide data, the maximum resolution for Bfactor consideration (--autob_highres) was set to the highest resolution in the RELION localres calculation, to avoid over sharpening. Local half map-based sharpening was applied to consensus maps using *phenix.auto_sharpen* with a window of 15x15x15 and overlap of 5, setting the resolution to the highest resolution observed in the RELION localres output. A near-complete model was fitted into the locally sharpened consensus map by using all-atom refine in Coot 0.9-pre (*16*) and this was further rigid body fitted into each of the focus refined maps using Chimera (*15*). The globally sharpened focused and consensus refinement maps for each class were then further combined together using *phenix.combine_focused_maps* to generate composite maps of much better clarity than the globally sharpened, or locally sharpened consensus maps in the distal membrane domain (Fig S7). Composite maps were carefully compared to the consensus refined map in Coot 0.9-pre (*16*) and Chimera (*15*) to ensure there were no gross map distortions, especially at map extremities, or artefacts at inter-map boundaries. Local resolution maps and Fourier shell coefficient (FSC) curves from half maps for all maps presented are shown in Figs. S4, 8-13 & 16-21.

Model building

An initial bovine active-state model was created by mutating an earlier mouse active-state model (PDB 6ZR2) in Coot 0.9-pre using *mutate residue range.* The initial model was then rigid body fitted into the consensus locally sharpened Active-10920-ii map using *phenix.real_space_refine*, followed by all-atom refine in Coot 0.9-pre and manual adjustment of the model. The resulting model then served to align the three focus refined maps for the Active-1092-ii state to generate a composite using *phenix.combine_focused_maps*. The model was then refined by cycles of manual refinement in Coot 0.9-pre (*16*) and automated refinement using Phenix-1.18.2 (*17*), guided by both the composite and consensus locally sharpened maps. Phospholipids and DDM molecules were added manually, and water molecules and ions were added using Coot *Find Waters* and manual addition, and checked manually for coordination geometry and density fit. This model was then rigid body fitted into each of the other maps, and refined in the same manner, checking and adding waters, phospholipids and detergent as necessary. Automated real-space refinements were performed using Phenix-1.18.2 against the composite maps using 5 macro cycles of minimization_global and local_grid_search, followed by Atomic Displacement Parameter refinement. Secondary structure restraints were not used, and Ramachandran restraints were set to Oldfield for Favored, and Emsley8k for Allowed and Outlier, to prevent genuine outliers being forcefully twisted. Statistics for all models are presented in Tables S2-14. In every case, the EMRinger score was superior for the composite map compared to the globally sharpened or locally sharpened consensus map. For the inhibitor-free active model, the model for Active-1092-ii was treated in the same way but in this case the locally sharpened map was used for refinement and secondary structure restraints were applied.

<u>Unbiased searches for biguanide densities</u>
As well as manual identification of biguanide-like densities, the ligand search function from Coot 0.9-pre was used to search for additional unassigned densities. The most promising 20 sites were identified using 50 ligand conformers, with the fractions for scoring and correlation set to 0.7, and assessed manually. In addition, difference maps were generated using the *phenix.real_space_diff_map* command using a near-final model without IM1092 coordinates, and by subtracting maps generated from the models by molmap in Chimera (*15*) from the globally sharpened consensus maps using *relion_image_handler* (*9*) (see Figs. 2 and S23). The difference maps were inspected for unassigned densities. A putative density adjacent to ND5-Tyr35 in the Active-1092-iii state that resembles IM1092 was not modelled because no feasible bonding interactions could be identified for it. Fitting of IM1092 into density was assessed by a combination of subjective explanation of the density and cross-correlation ($C\text{-}C_{mask}$) of the IM1092 molecule by using Phenix-1.18.2. Where $C\text{-}C_{mask}$ was ~0.5 or poorer and the density was subjectively not well explained, IM1092 was not included in the final model.

<u>Statistical methods</u>
For kinetic and scattering assays, all data were recorded with at least three technical replicates and number of replicates and measures of error (95% confidence interval or S.E.M) are indicated, and errors were propagated where appropriate.

### Supplementary Text

<u>Background: Characteristics of the active, deactive and slack classes</u>
Global classification of cryo-EM datasets for bovine complex I typically resolve three major classes (*11*, *12*), and the same behavior is observed here (Fig. S2). The classes are differentiated by both large-scale global features and by the status of individual structural elements. The ***active*** classes exhibit the most acute (closed) angle between the hydrophilic and membrane domains (Fig.

S22A), well-ordered structures for all elements of the Q-binding site (including the ND3 TMH1-2, ND1 TMH5-6 and NDUFS2 β1-β2 loops), a specific orientation of NDUFS7-Arg77 and its adjacent loop, and α-helical ND6-TMH3. This class has been described as the 'active' state of mouse, bovine and porcine complex I (*2*, *11*, *12*, *18*) and as the 'closed' state of ovine complex I (*19*). The ***deactive*** classes, identified previously for the mouse, bovine and porcine enzymes (*2*, *8*, *12*, *18*, *20*), exhibit a less acute (more open) angle (Fig. S22A), poorly ordered loops in ND3 (residues 31-46), ND1 (62-64 and 207-216) and NDUFS2 (46-64), a different orientation of NDUFS7-Arg77 and adjacent β-strand, plus a π- bulge in ND6-TMH3. The ***slack*** classes also exhibit the same disordered loops, β-strand in NDUFS7 and π-bulge in ND6-TMH3, but large portions of subunit NDUFA11 and the C-terminal lateral helix of ND5 (residues 562-606) are also not resolved (*11*, *12*). Classes of ovine complex I referred to as 'open' states resemble both the deactive and slack states of the bovine enzyme (*19*).

Local-first classification

To classify different sub-states from the final set of selected particles (Fig. S2), we tested different orders of image processing steps. First, the particles were classified globally into the active, deactive and slack states, and then subjected to focused classification with a soft mask around the areas of interest. This is the simplest strategy and was followed previously to classify a dataset of the bovine enzyme in nanodiscs (*12*). We found the resultant classes here exhibited lack of clarity in the density shapes within the classified regions. Therefore, an alternative 'local-first' strategy was tested in which the order of the classification steps was reversed. The maps exhibited enhanced separation of differently shaped densities in the classified regions, which we ascribe to an improved separation of the local densities due to better signal-to-noise during classification over small density volumes. The local-first strategy we applied (Figs. S5 & S6) started from a consensus refinement of all the (heterogenous) particles followed by: 1) particle subtraction for the domain of interest; 2) focused refinement on the domain of interest to improve alignment; 3) focused classification with a small, soft mask over the area of interest; 4) particle reversion to include density for the entire protein; 5) global classification.

Relative motions of classes in the biguanide-inhibited dataset

To improve qualitative descriptions of apparent angle between the hydrophilic and membrane domains as 'open' or 'closed' we used the positions of three Cα atoms (NDUFB3-Ala75, NDUFA1-Val14 and NDUFS1-Glu165) to define the angle 'HD-MD flex' (Fig. S22A). Similarly, we used 'HD-MD twist', defined by the dihedral angle between four Cα atoms (NDUFB3-Ala75, NDUFA1-Val14, NDUFS1-Leu651 and NDUFV1-Ser38) (Fig. S22B), to describe how the hydrophilic domain rotates on the membrane domain, and 'HD-MD tilt', the dihedral angle between NDUFB8-Phe111, ND5-Tyr42, NDUFA13-Leu44 and NDUFV1-Asp292 (Fig. S22C) to describe how it tilts relative to it. These three parameters are most affected by structural elements at the domain interface, which also form the Q-channel, and, accordingly, local classifications of the Q-channel region (Fig. S5) separated different subclasses with both different degrees of Q-channel loop order and subtle differences in these global angles (see Deactive-1092-i, ii and iii and Slack-1092-i and ii in Fig. S21). Between the proximal and distal sections of the membrane domain we describe the two angles 'PMD-DMD flex' (orthogonal to the membrane plane) and 'PMD-DMD bend' (in the membrane plane) using the Cα atoms for NDUFB3-Ala75, ND2-Gln168 and NDUFA1-Val14 (Fig. S22D) and NDUFB3-Ala75, ND2-Met211, and NDUFA1-Val14 (Fig. S22E), respectively. Classification on the ND5 lateral helix (Fig. S6) reveals the classes with the most ordered lateral helix (Active-1092-ii and Deactive-1092-iv) have the most closed flex and bend angles compared their helix-disrupted counterparts Active-1092-iii and iv, and Deactive-1092-v and vi, respectively (Fig. S22). Overall, the angles between the domains clearly demonstrate the similar global conformations for the substates in each major class, including the clear demarcation of active substates with the smallest HD-MD flex angles.

Q-site loop conformations from analysis of the biguanide-inhibited dataset

In all the active subclasses separated here, as well as in the inhibitor-free active state, the variable structural elements in subunits ND3, ND1, NDUFS2 and NDUFS7 that form the Q-channel are well ordered and essentially identical. In the deactive state these elements (ND3 TMH1-2 loop, ND1 TMH5-6 loop, and NDUFS2 β1–β2 loop) are usually considered disordered, but in the biguanide-bound substates Deactive-1092-i, ii and iii, local-first classification resolved them partially into different conformations. ND3 residues 31-45 and ND1 209-214 remain unresolved, but NDUFS2 46-64 are resolved in Deactive-1092-ii, and ND3 25-31 and ND1 215-219 are resolved in all three states with different degrees of ordering. The biguanide-bound Q-channel in Deactive-1092-ii thus differs from that in Deactive-1092-i and iii, as well as from the DDM-bound Q-channel in deactive bovine complex I in nanodiscs (PDB 7QSM) (*12*). We note that in the biguanide dataset, no DDM is observed in the Q-channel, and so must be out-competed by the biguanide ligand, and the different ligand identity likely creates order in the local structures. By contrast, in the two biguanide-bound slack substates the NDUFS2 β1–β2 loop (46-64) is in the same 'inward collapsed' state as it is in with a cholate molecule bound in the slack Q-channel in nanodiscs (*12*). In this case, the related structures with different ligands bound (with the backbone carbonyl of NDUFS2-Gln54 hydrogen bonding to the backbone amide of NDUFS7-Arg77, blocking off the top of the Q-channel) indicate the conformation is specific to the slack state, not a biguanide-induced effect. Fig. S24 compares the three different ordered conformations of the NDUFS2 loop observed. ND1 residues 203-217 are resolved in only one of the two biguanide-bound slack states, whereas ND3 20-50 remain poorly resolved in both. All structures published so far are therefore consistent with exposure of ND3-Cys39 to derivatizing agents such as NEM in both the deactive and slack states. Finally, we note that a focus-revert-classify approach applied to the as-prepared (native) ovine enzyme (*19*) also did not completely resolve the Q-channel elements, and different positions for the loops in NDUFS2 or ND1 were not identified. The maps for deactive substates (Deactive-1092-iv, v and vi) that were not locally classified on lateral helix region display a similar degree of order at the Q-channel loops, to that observed in the native ovine open classes, suggesting a combination of classification strategies may be required in the future to better separate deactive /open substates.

The slack state in the biguanide-inhibited dataset

Our biguanide-bound slack classes exhibit a π-bulge in ND4 TMH6. The same feature was described previously for the slack class in bovine nanodiscs (PDB 7QSO) (*12*), and is also present in some of the open states of ovine complex I (PDB 6ZKV, 6ZKN and 6ZKM for the deactive-open4, and rotenone-open2 and -open3 classes) (*19*). A $Q_{10}$ molecule is bound nearby in the nanodiscs structure, and a rotenone molecule is present in the same site in the rotenone-bound open2 and open3 states (*12*, *19*). While the slack classes described here contain only a weak and unassigned density in this position, it is thus possible that ligand binding stabilizes the π-bulge. It was not previously noted that the ND4 TMH6 π-bulge rotates Tyr152 out of its position pointing towards the ND5 lateral helix (where it is found in the active and deactive states), into hydrogen bonding distance of the ND4 Glu123-Lys206 ion pair in the slack state (Fig. S15). These two residues are part of the central hydrophilic axis running along the center of the membrane domain and they are crucial for energy transduction. PropKa (*21*) calculations suggest that the pKa of Lys206 shifts from around 8.5 to 7.5 between the Active-1092-ii/Deactive-1092-iv and Slack-1092-ii states. Analogous residues in subunit ND5 (Tyr174 and Glu145-Lys223) are present in the same positions when the subunit models are overlayed. ND4-Tyr152, via a water molecule, forms part of a hydrogen-bonding network that supports a π-bulge structure in the ND5 lateral helix (as does ND4-Tyr148, in its interaction with the ND5-Glu559 sidechain) perhaps explaining why the lateral helix is not well-ordered in the slack state. It is possible that the π-bulge forms in ND4 TMH6 and Tyr152 rotates between different interacting partners during catalysis.

Composite maps for the biguanide-bound slack states lack clear density for ND6-TMH4, but when a Gaussian filter (width 1) was applied in Chimera (*15*), a density was revealed that suggests the helix has moved laterally towards the distal hydrophobic domain in a position intermediate between that of the active/deactive bovine states, and that of the *Yarrowia lipolytica* and *Thermus thermophilus* enzymes (*22*, *23*). The bovine enzyme, like its ovine counterpart, contains a β-hairpin between ND6-TMH4 and 5, whereas the mouse enzyme contains only a loop (*2*). In the biguanide-bound slack classes, movement of TMH4 and swiveling around of the β-hairpin have occurred (Fig. S14). The same changes are not observed in inhibitor-free bovine slack structures (here in DDM, or in nanodiscs (*12*)) in which the helix density appears only in its deactive-state position. Different rearrangements were observed in the heat-treated ovine open4 state, with the β-hairpin residues converted to an α-helical segment and ND6-TMH4 tilting backwards towards ND1, and in the ovine rotenone-open2 and open3 states (*19*), where the β-hairpin and the first half of ND6-TMH4 appear in the same position as observed here in the active and deactive states. The specific arrangement observed here may thus result from biguanide interactions (in the Q-channel or with nearby phospholipids). As we were unable to produce a final map that contained both this density information as well as high-resolution information, this feature was not built into the deposited PDB models. Instead, a crude approximation is presented in Fig. S14C. It is clear that ND6-TMH4 and the structures between TMH4 and 5 vary in different conditions and different species, but the relevance, if any, of the different structures for catalysis is unclear.

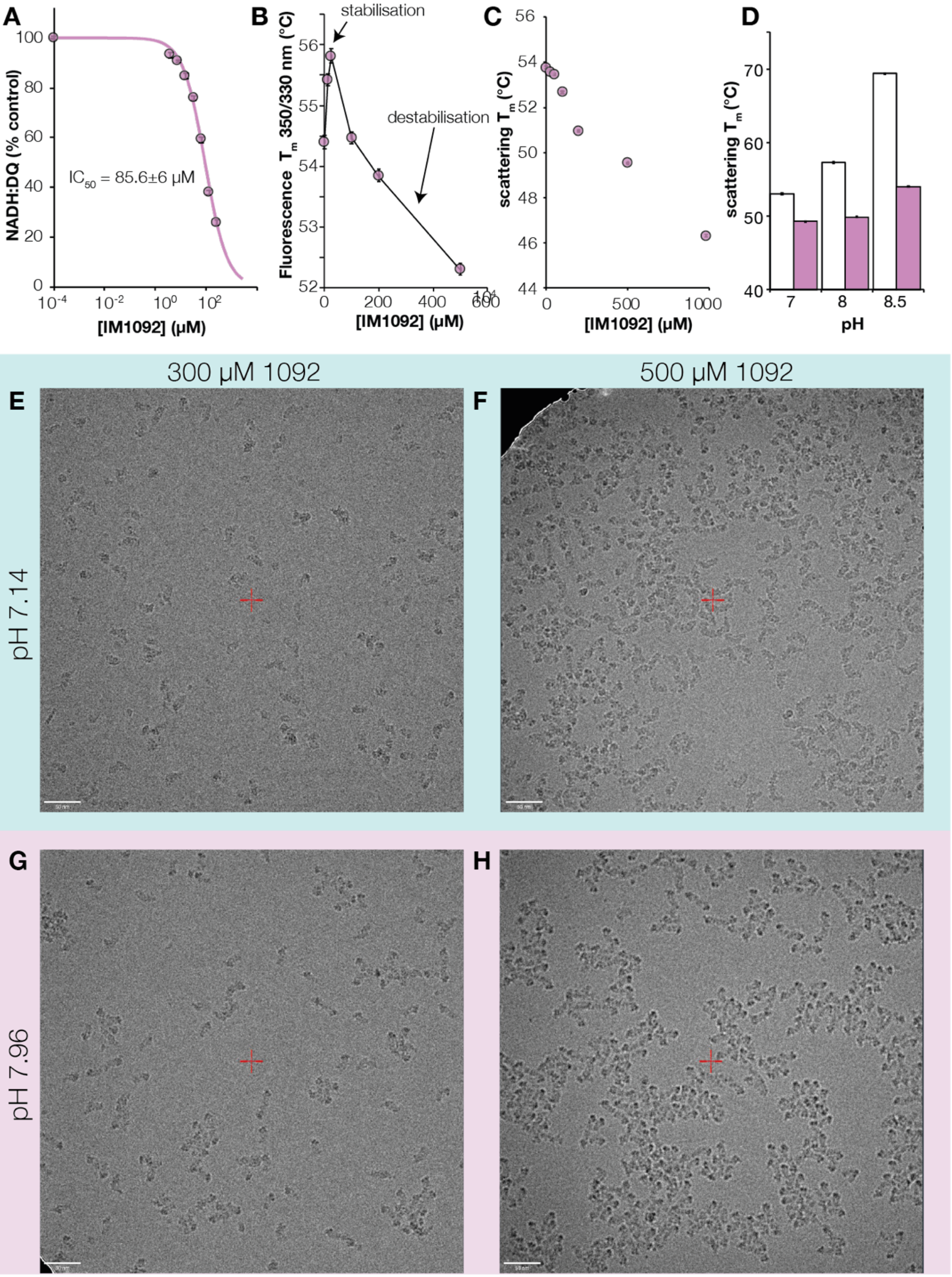


**Fig. S1. Effects of IM1092 on the activity and aggregation of detergent-solubilized complex I.** A) The effect of IM1092 concentration on the NADH:DQ oxidoreduction activity of purified complex I. B) The effect of IM1092 concentration on protein thermostability measured as 350/330 nm fluorescence at 120 μg $mL^{-1}$ protein. Points are joined for illustration, not fitted. C) The effect of IM1092 concentration on the protein melting temperature for protein aggregation. D) The effects of IM1092 on aggregation at different pH values. Control (white) and 500 μM IM1092 (orchid) at 120 µg $mL^{-1}$ complex I. E-H) Cryo-EM images of purified bovine complex I at 2.5 mg $mL^{-1}$ in buffer containing 20 mM Tris-HCl, 150 mM NaCl and 0.05% DDM (pH 7.14 or 7.96 at 20 °C). Data show means and standard error of three independent measurements. Images are from a Talos Arctica operating at 200 kV at 73k x magnification, around -2.5 µm defocus with a 100 µm objective aperture and 50 µm C2 aperture. White scale bars show 50 nm. Aggregation is more severe at both higher pH and higher concentration of IM1092.

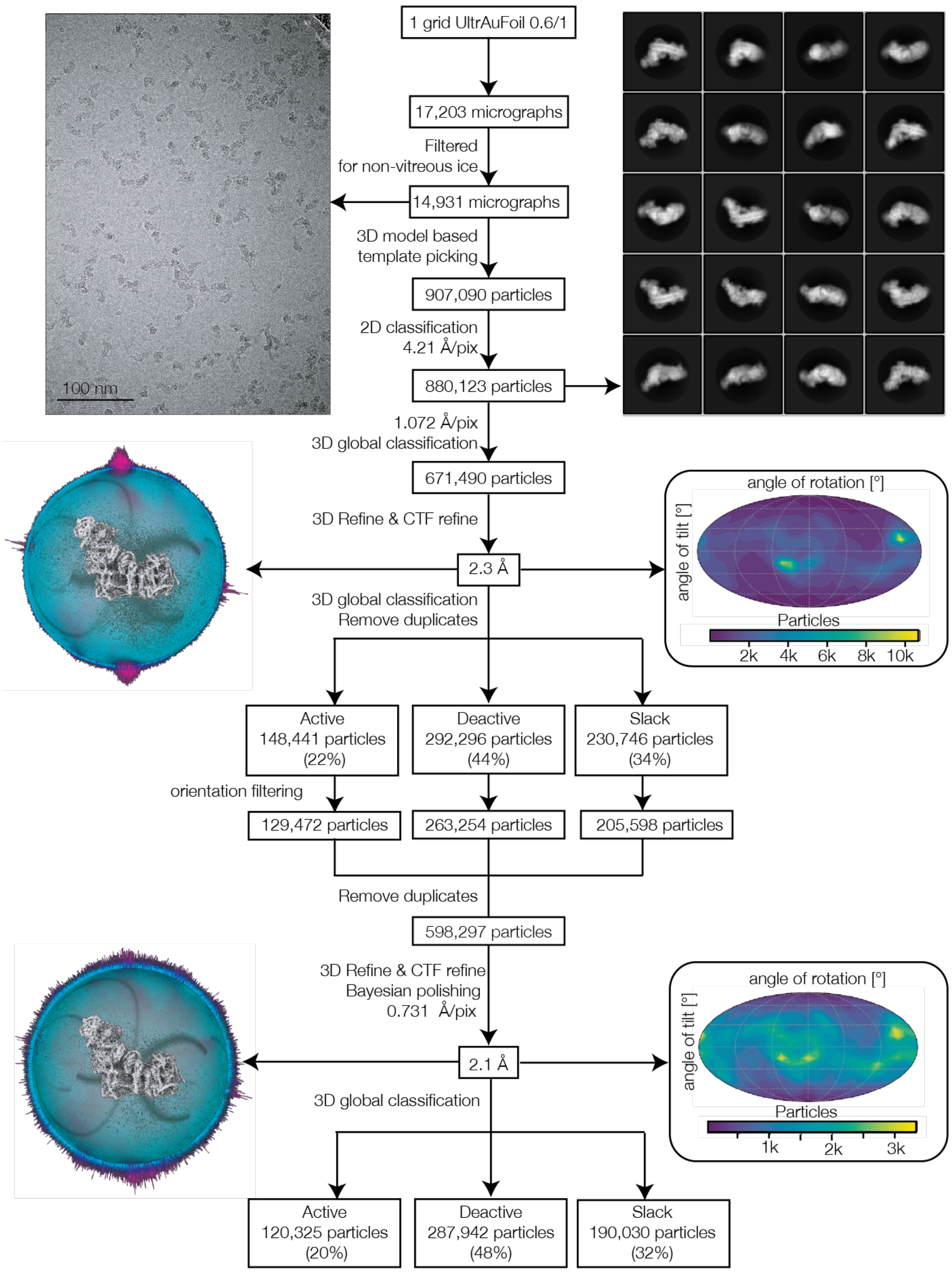


**Fig. S2. Data processing scheme for the biguanide-inhibited dataset.** A typical micrograph and selected 2D classes are shown, along with 3D and 2D projections of orientations before and after particle filtering.

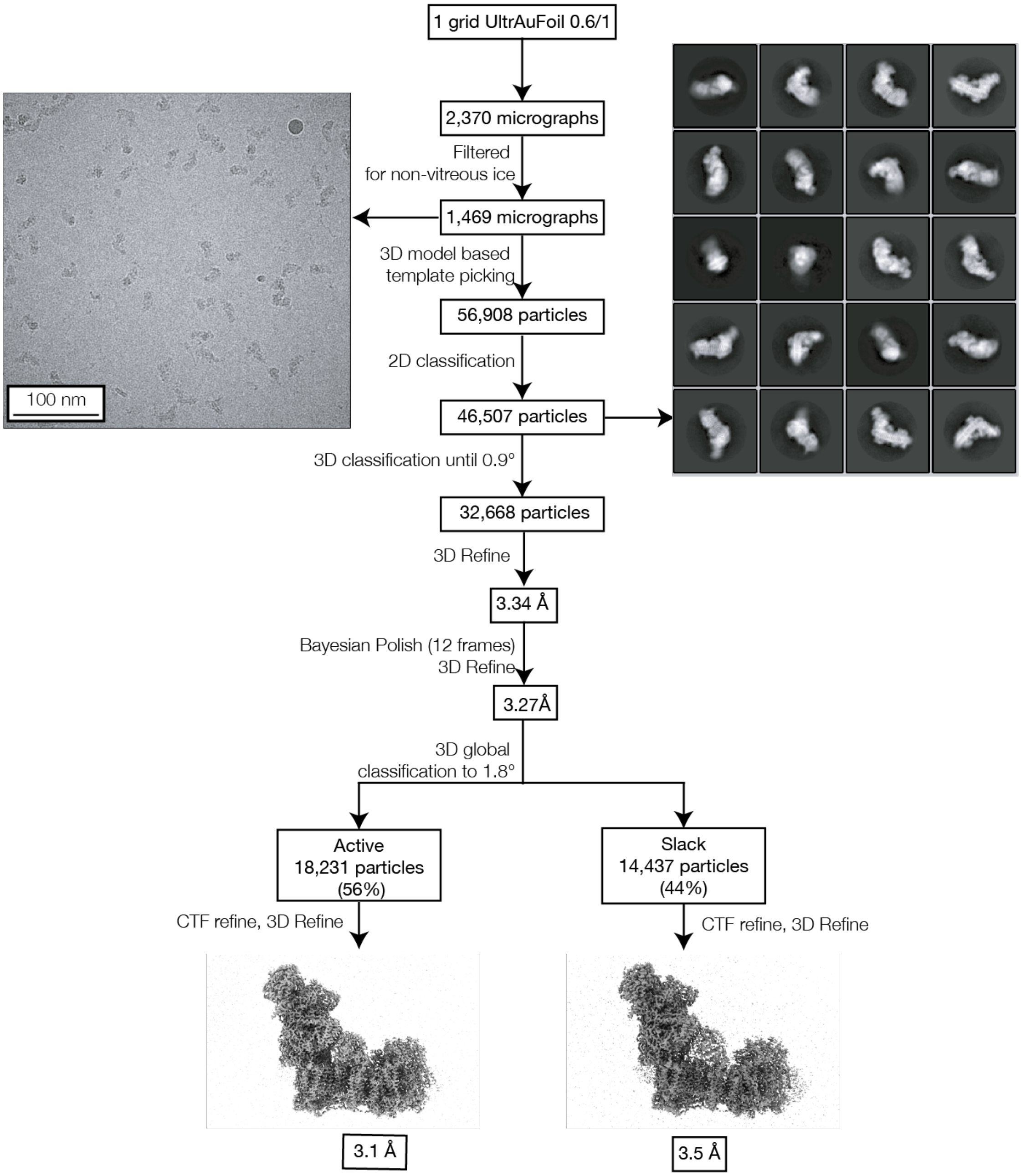


**Fig. S3. General classification scheme for inhibitor-free bovine complex I.** An example micrograph and selected 2D classes are shown.

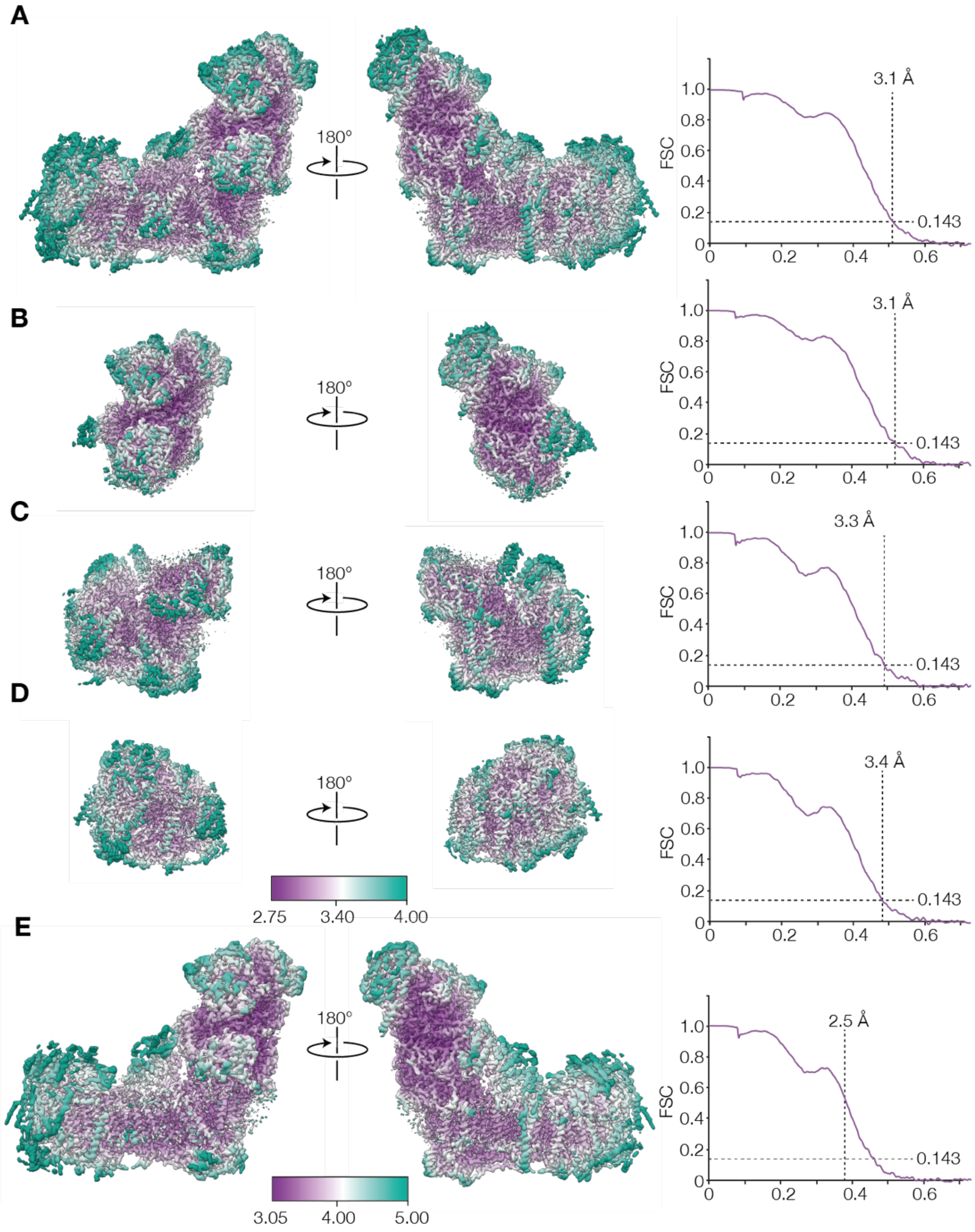


**Fig. S4. Local resolution maps for inhibitor-free active and slack with FSC curves for half maps.** A) Active consensus, B) Active hydrophilic domain, C) Active proximal hydrophobic domain, D) Active distal hydrophobic domain, and E) Slack consensus.

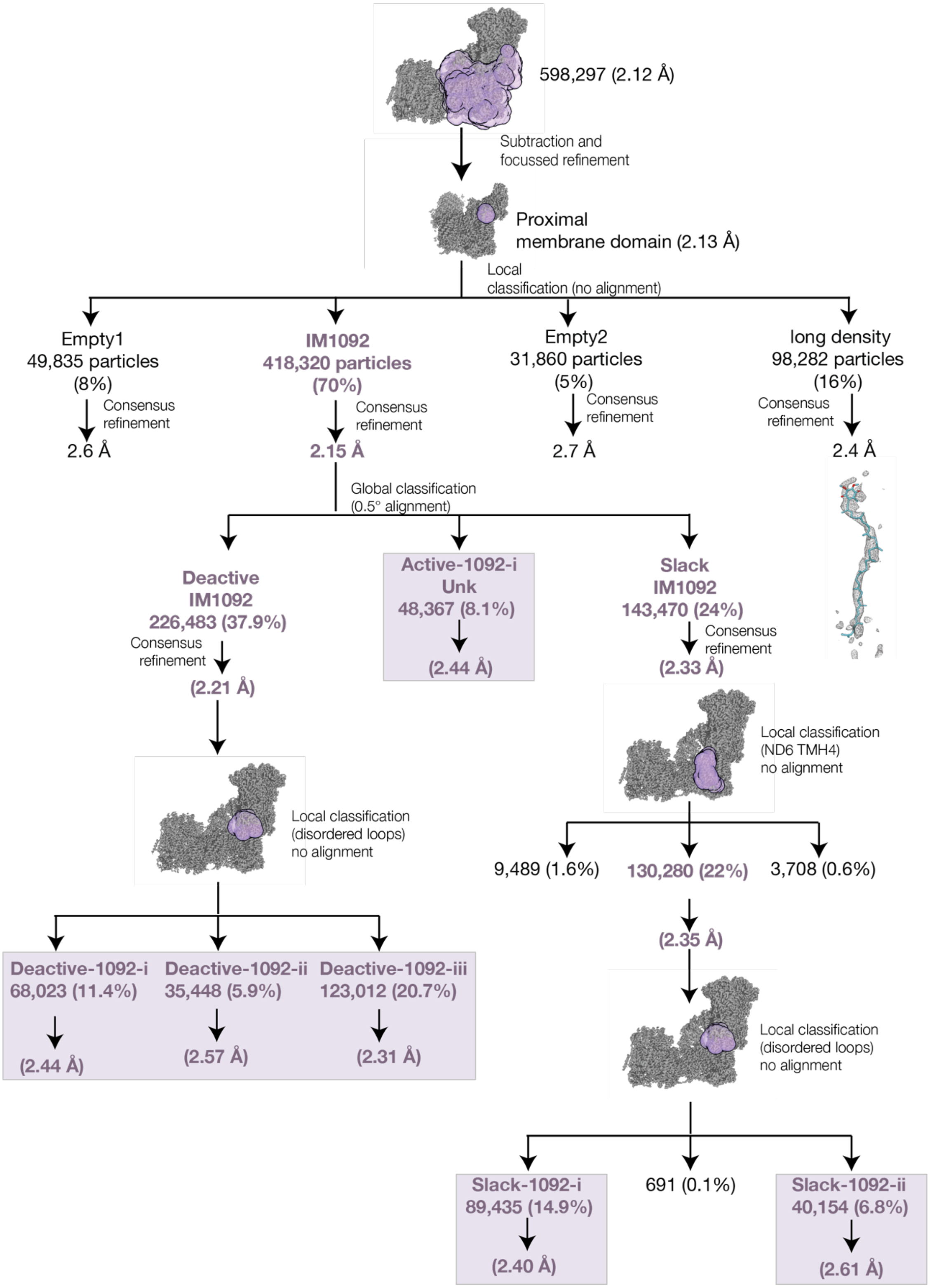


**Fig. S5. Particle classification scheme for the ubiquinone-binding channel region.** Images show the positions of masks around the experimental density used for classification. The maps discussed in this study are highlighted by lilac boxes. 'Empty' refers to a lack of connected density in the channel: we note that it is not certain whether the channels are genuinely empty, or if local resolution or classification were insufficient to elucidate ligand heterogeneity. The 'long density' observed overlays well with the $Q_{10}$ molecule modelled in PDB-7QSK (*12*) and therefore likely represents endogenous $Q_{10}$ retained from the native membrane. PDB-7QSK was fitted in this map, and density is shown in mesh for the difference map between this class and a molmap generated density of PDB-7QSK, within 10 Å of the $Q_{10}$ molecule. The density observed in Active-1092-i described as Unk (unknown) resembles that present in the active-apo class of bovine complex I in nanodiscs (EMD-14133) (*12*) and may represent a heterogeneous mixture of ligands or ligand-binding poses.

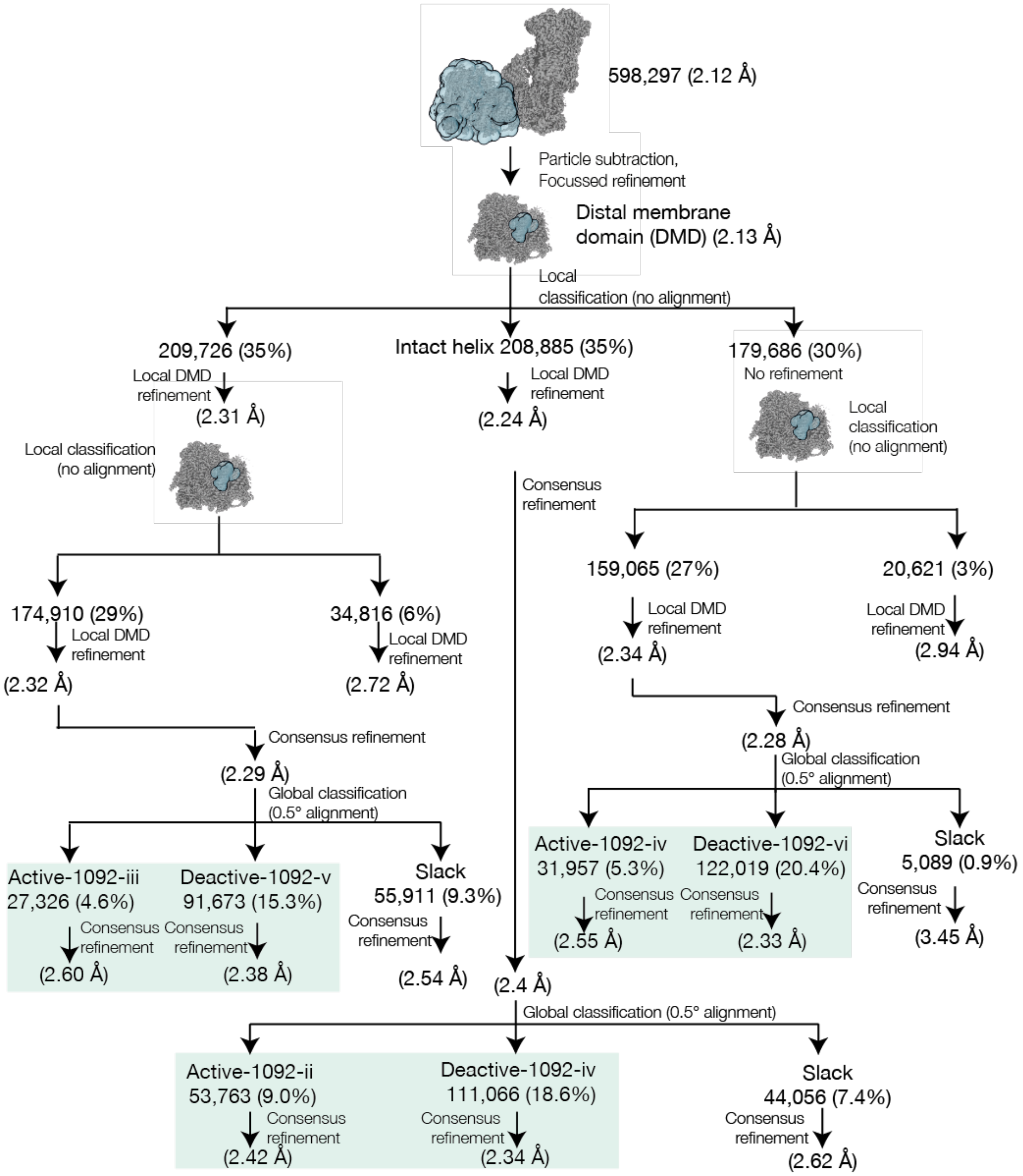


**Fig. S6. Particle classification scheme for the distal membrane domain.** Images show the position of masks around the experimental density used for classification. Maps used in this study are highlighted by green boxes.

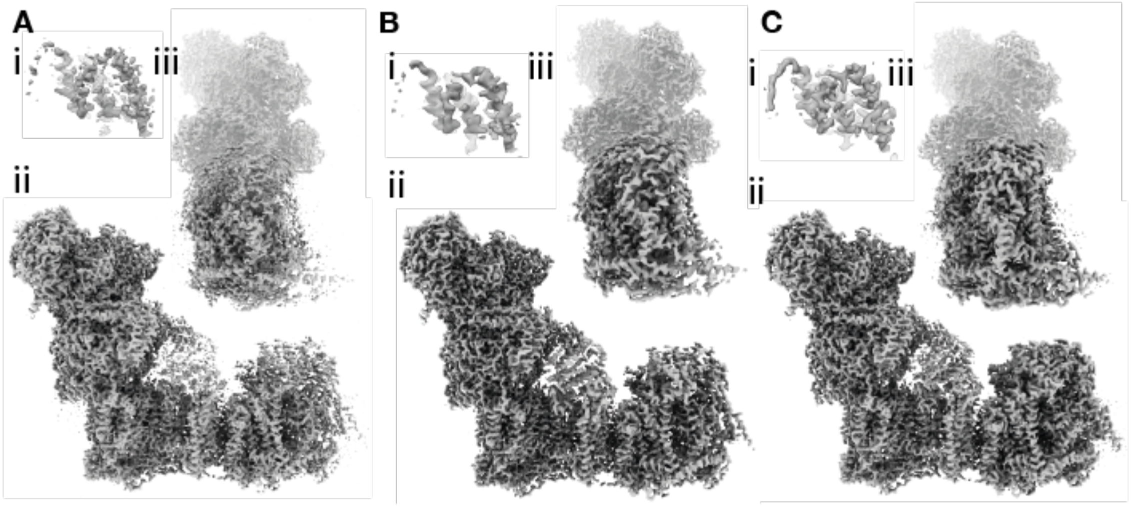


**Fig. S7. Example of the improvement in map quality obtained using a composite map (Slack-1092-ii).** Density is shown for A) globally sharpened consensus, B) locally sharpened consensus, and C) globally sharpened composite maps. Views show ii) the entire complex, iii) rotated by 90 degrees, as well as i) an inset showing subunit NDUFB7 where the N-myristoyl modification becomes clearly visible in panel C indicated by an arrow.

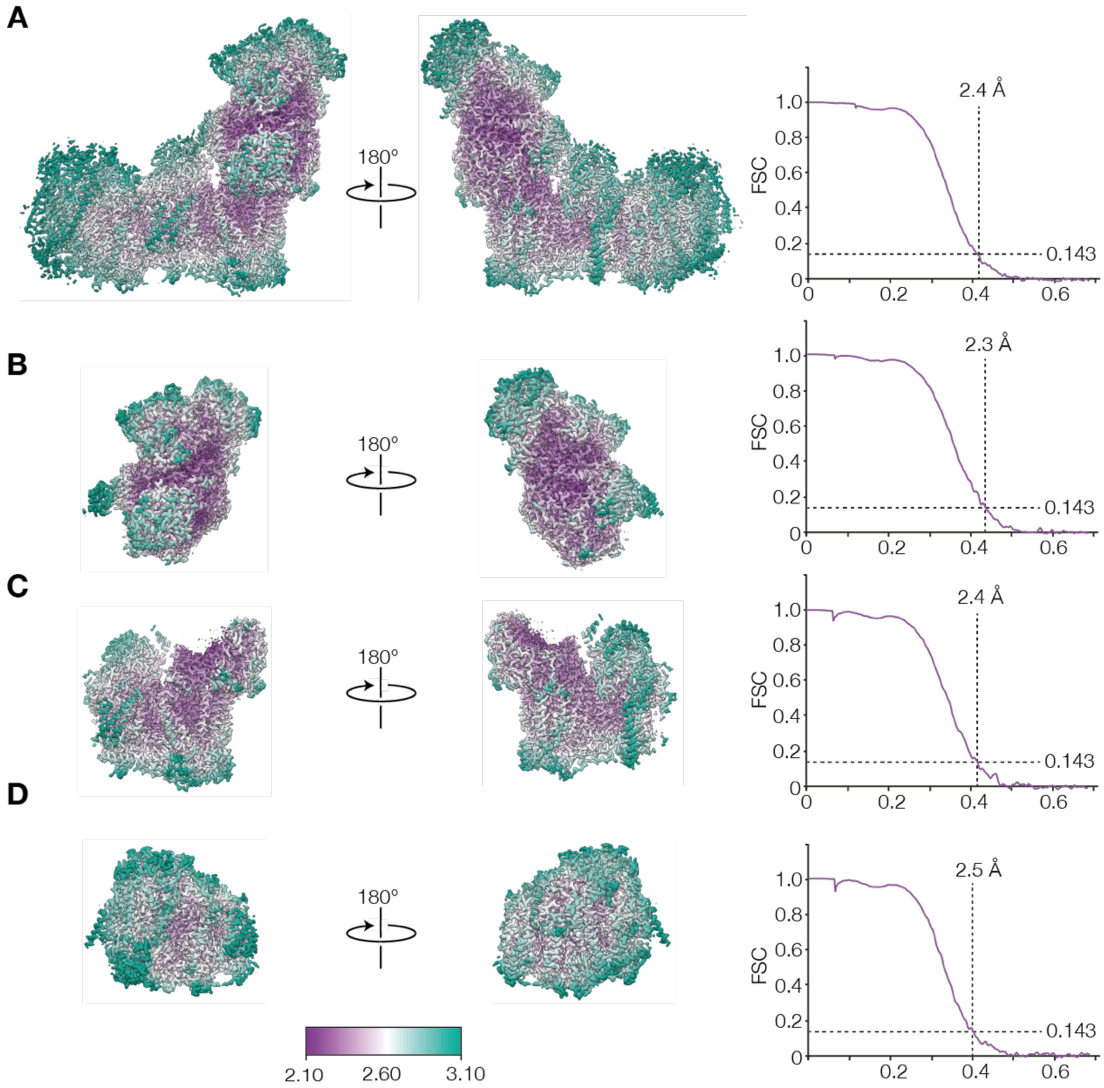


**Fig. S8. Local resolution maps for Active-1092-i and FSC curves for half maps.** A) Consensus, B) hydrophilic domain, C) proximal hydrophobic domain, D) distal hydrophobic domain.

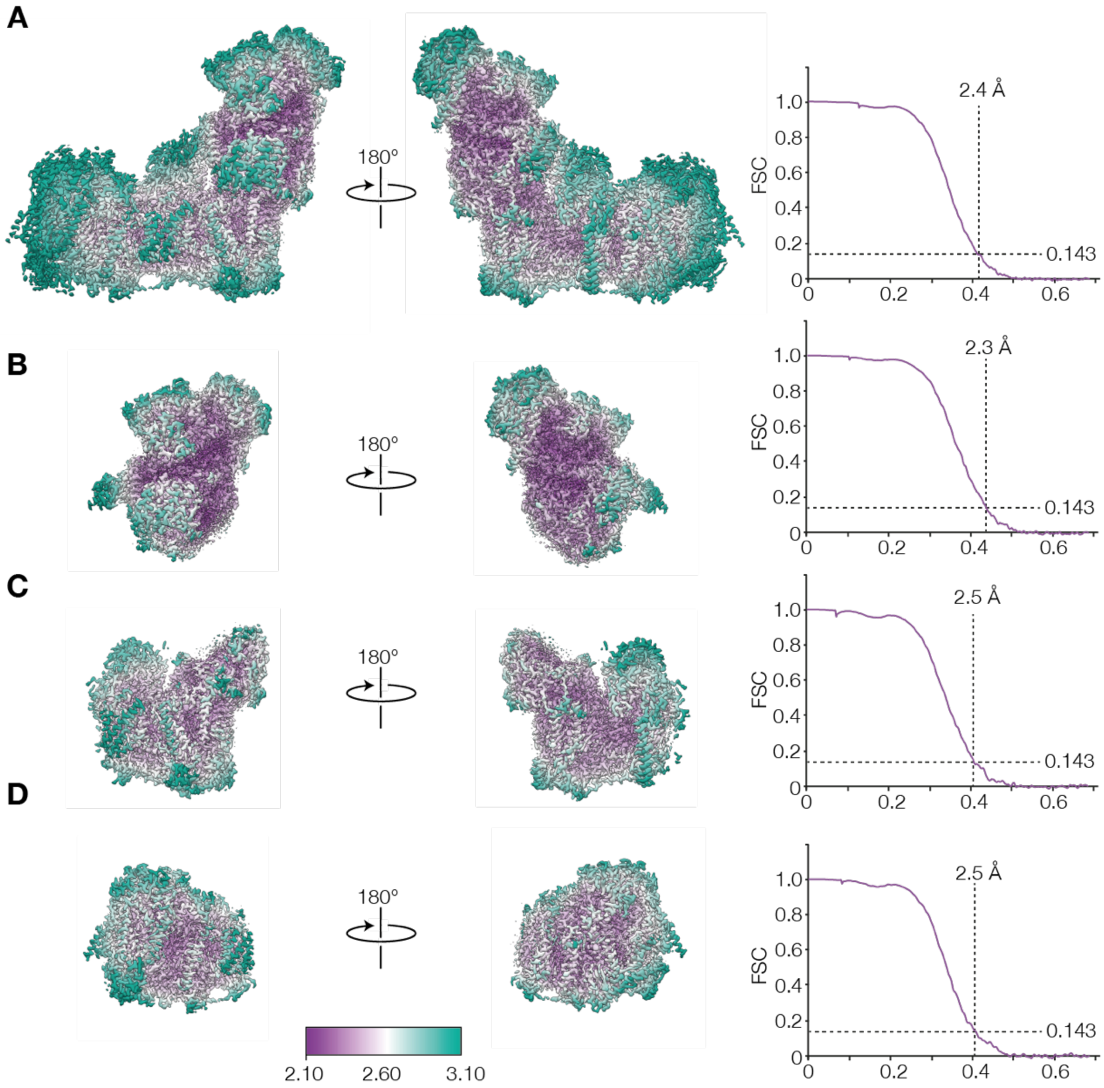


**Fig. S9. Local resolution maps for Deactive-1092-i and FSC curves for half maps.** A) Consensus, B) hydrophilic domain, C) proximal hydrophilic domain, D) distal hydrophobic domain.

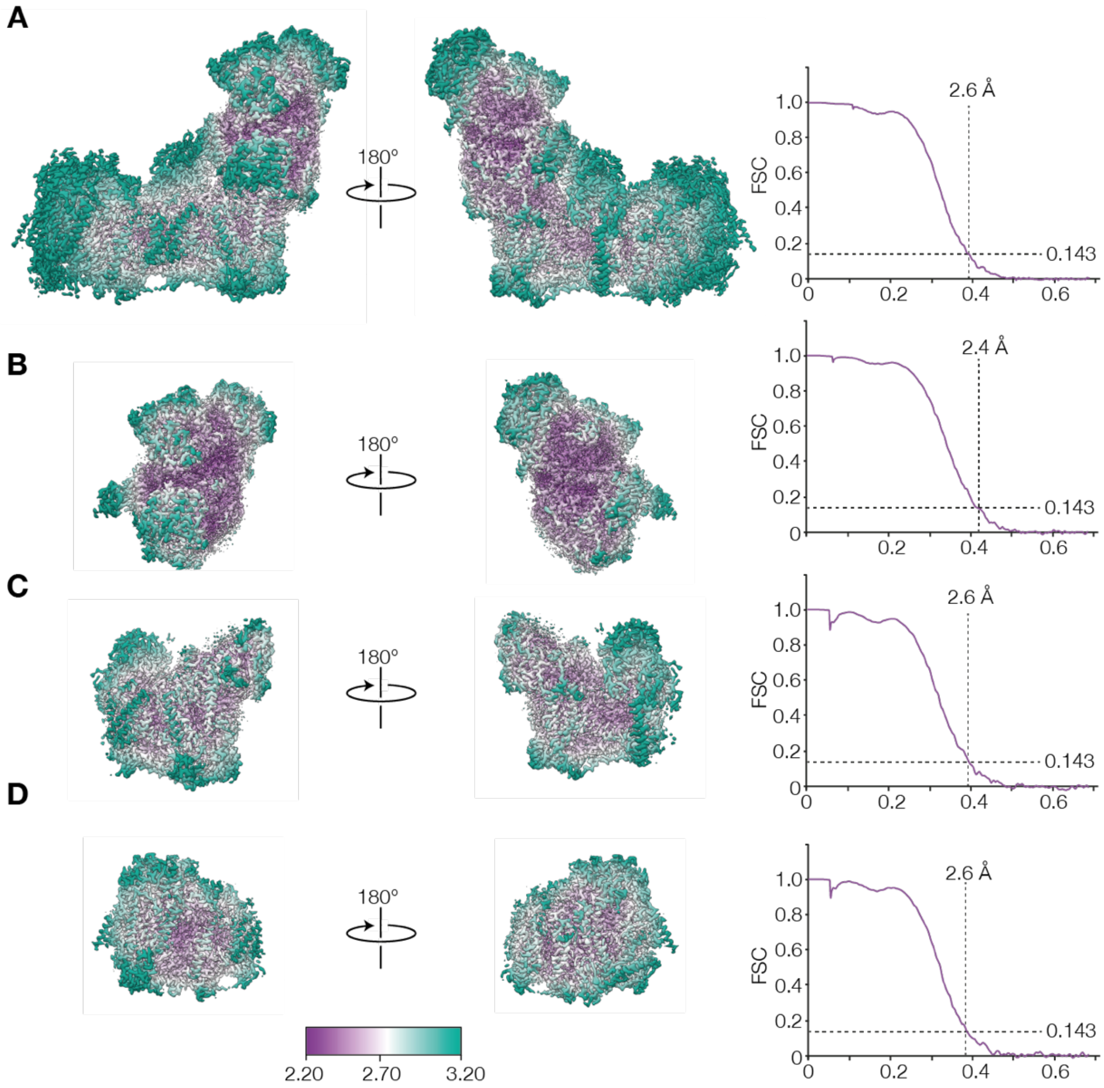


**Fig. S10. Local resolution maps for Deactive-1092-ii and FSC curves for half maps.** A) Consensus, B) hydrophilic domain, C) proximal hydrophobic domain, D) distal hydrophobic domain.

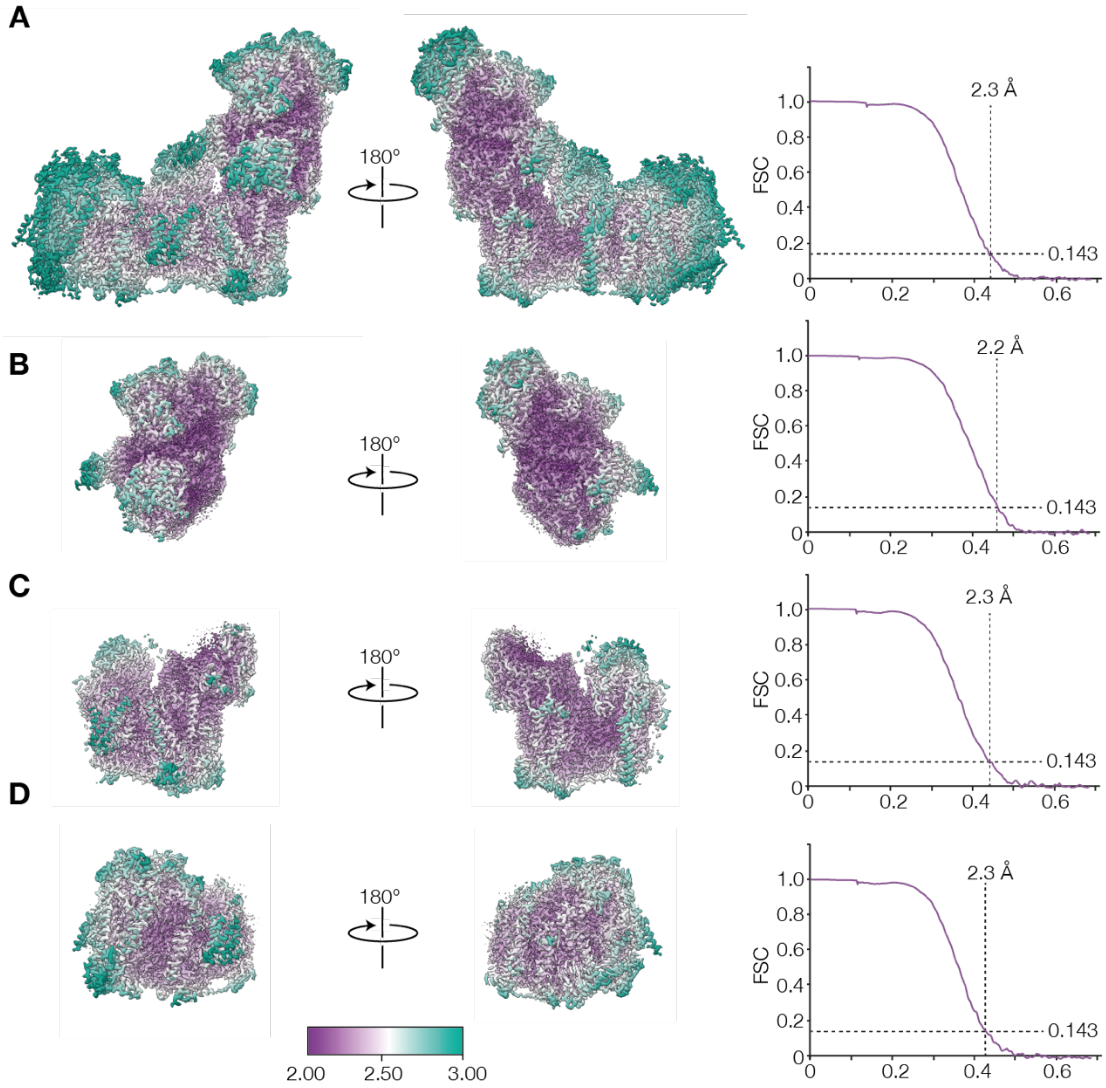


**Fig. S11. Local resolution maps for Deactive-1092-iii and FSC curves for half maps.** A) Consensus, B) hydrophilic domain, C) proximal hydrophobic domain, D) distal hydrophobic domain.

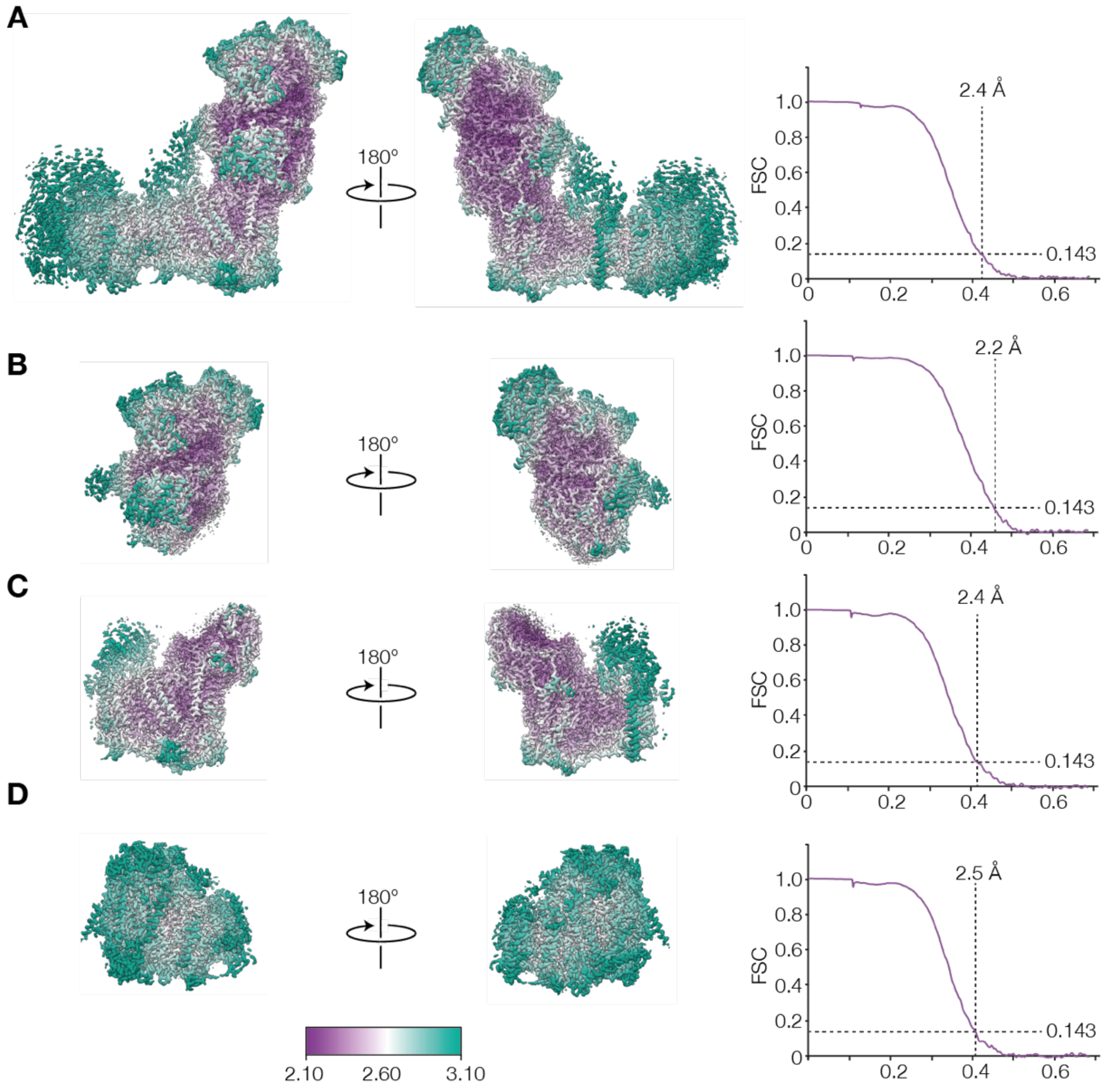


**Fig. S12. Local resolution maps for Slack-1092-i and FSC curves for half maps.** A) Consensus, B) hydrophilic domain, C) proximal hydrophobic domain, D) distal hydrophobic domain.

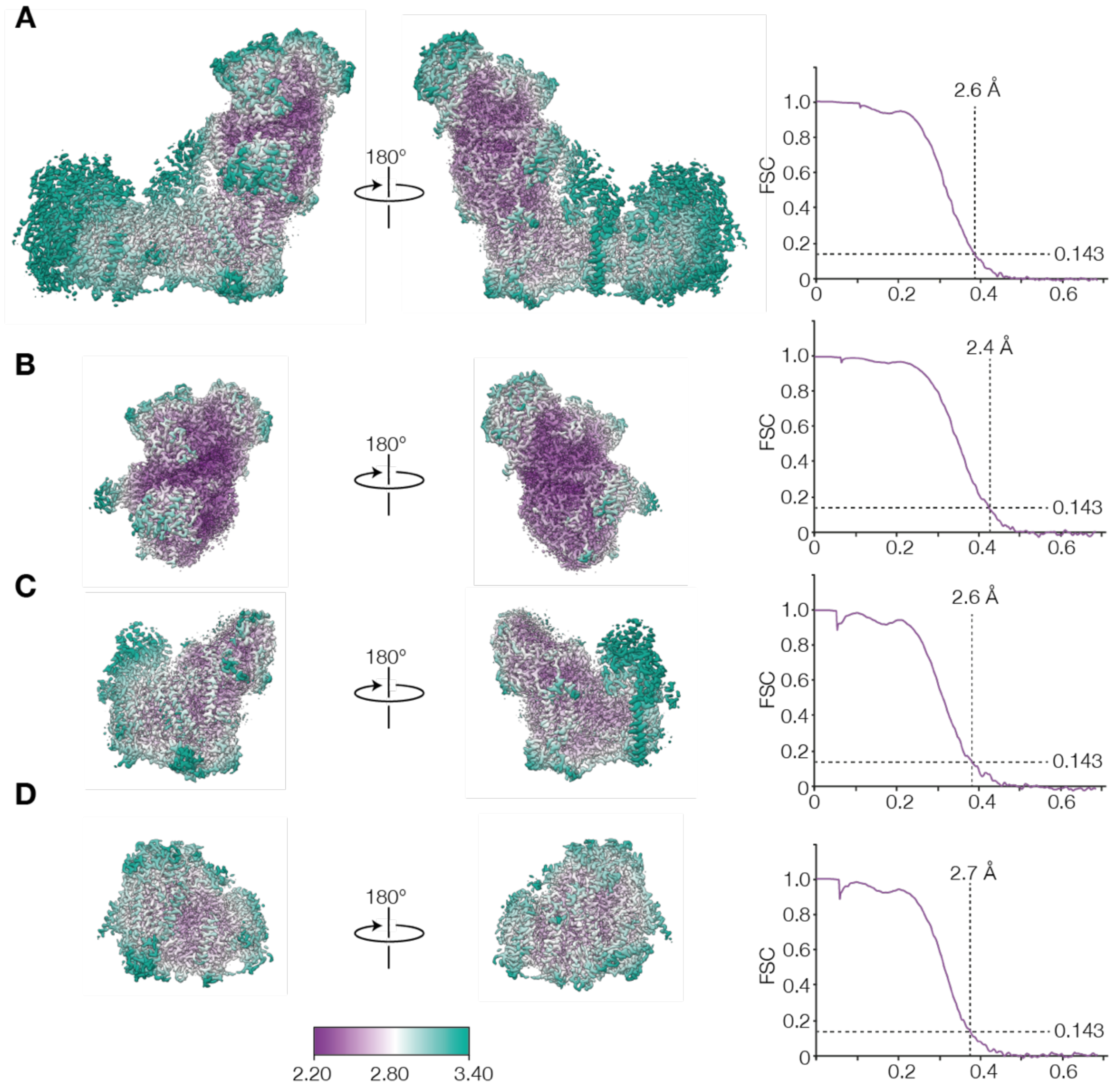


**Fig. S13. Local resolution maps for Slack-1092-ii and FSC curves for half maps.** A) Consensus, B) hydrophilic domain, C) proximal hydrophobic domain, D) distal hydrophobic domain.

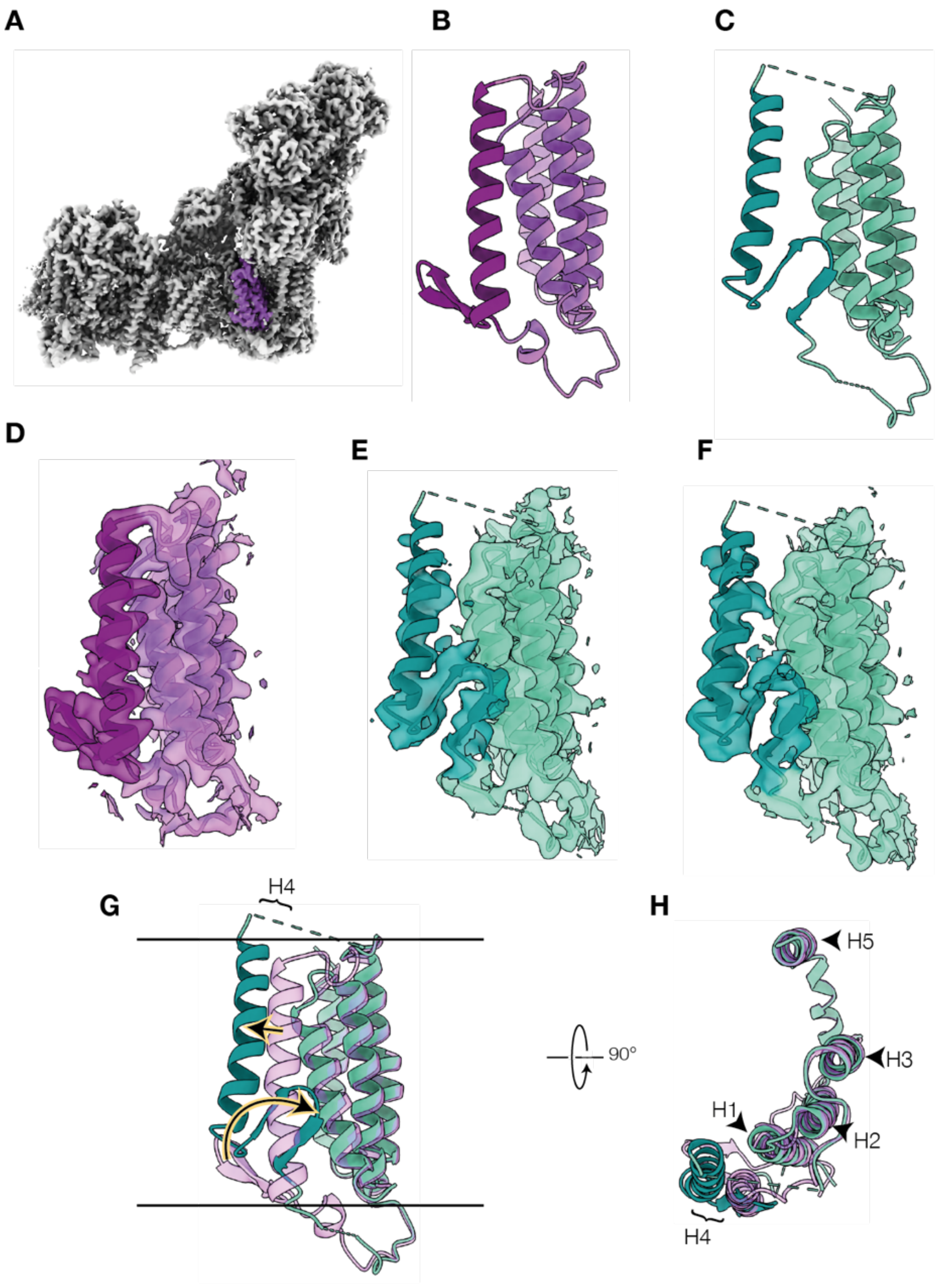


**Fig. S14. Changes to the structure of the ND6 subunit between the deactive and slack states**. A) ND6 from a representative class Deactive-1092-ii is indicated in purple. B) The model for ND6 from Deactive-1092-ii. C) An approximate model of ND6 in the biguanide-bound slack state (for illustrative purpose only; see supplementary text). D-F) Gaussian-filtered density maps for the ND6 subunit in D) Deactive-1092-ii; E) Slack-1092-i; and F) Slack-1092-ii each with density zoned and colored according to 3 Å proximity to the appropriate PDB models using ChimeraX (*24*). ND6 THM4 and the TMH4-5 β-hairpin are colored dark purple (deactive) or dark teal (slack). G-H) Overlays of the models showing the movement of TMH4 and the β-hairpin. Horizontal lines in G) depict the approximate position of the phospholipid bilayer. Purple, deactive-1092-ii; teal, slack classes.

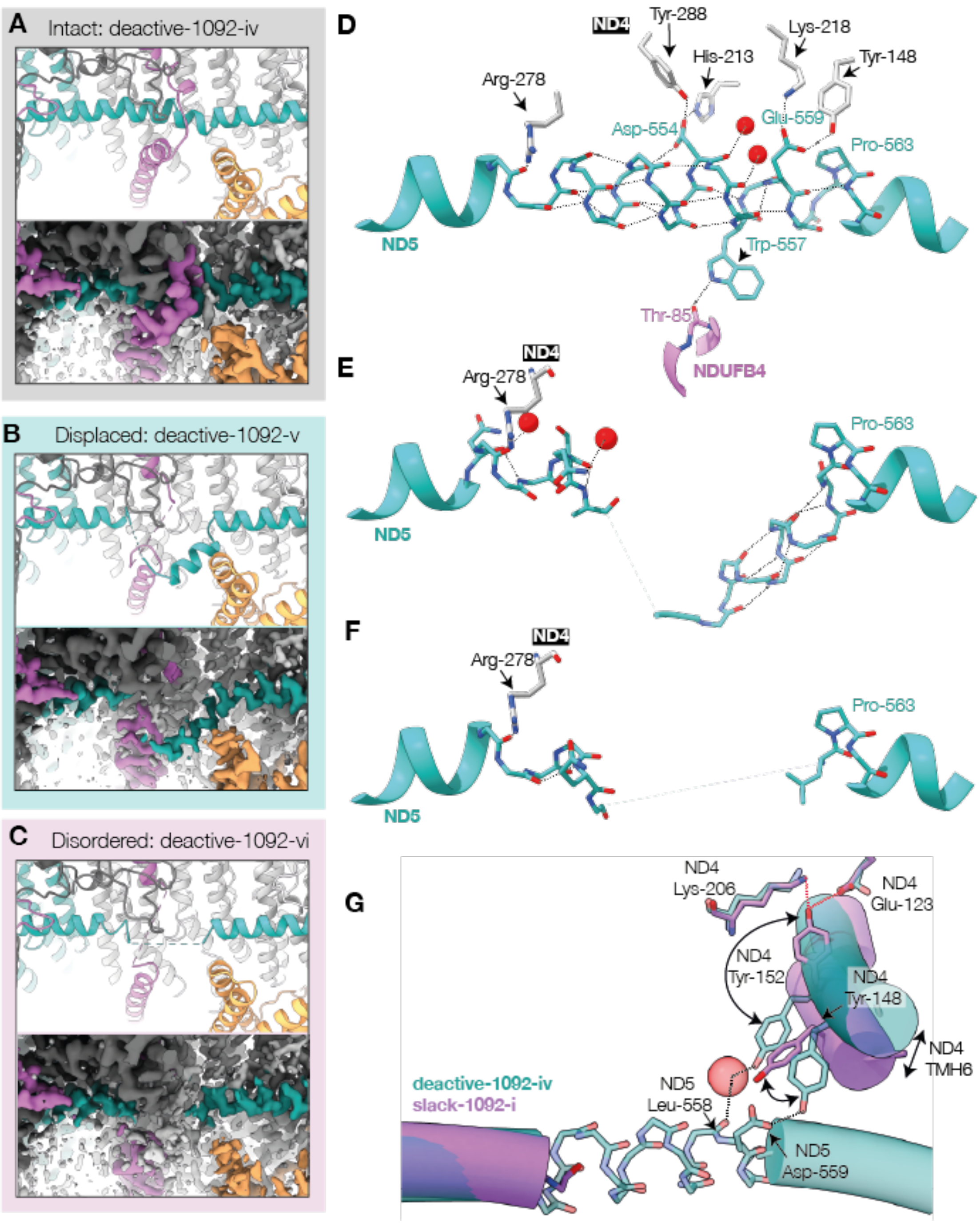


**Fig. S15. Biguanide-induced distortion and disordering of the ND5 lateral helix and NDUFB4 loop in deactive states, and lateral-helix associated conformational changes in the slack state.** A-C) PDB models and densities for the ND5 lateral helix in A) Deactive-1092-iv, B) Deactive-1092-v, C) Deactive-1092-vi. Density maps are colored using ChimeraX (*24*) by 3 Å proximity to the modelled structure: teal, ND5; orchid, NDUFB4; orange, NDUFA11; dark grey, NDUFB8. D-F) Hydrogen-bonding stabilization of π-bulges in the ND5 lateral helix for Deactive-1092-iv, Deactive-1092-v and Deactive-1092-vi. Water molecules are represented by red spheres, black dotted lines show hydrogen bonds: teal, ND5; orchid, NDUFB4; white, ND4. G) Overlay of Deactive-1092-iv (teal) and slack-1092-i (orchid); rotation of ND4 TMH6 in the slack state causes loss of two stabilizing interactions (black dotted lines) with the ND5 lateral helix, but creates additional hydrogen bonds (red dotted lines) with ND4-Lys206 and Glu123.

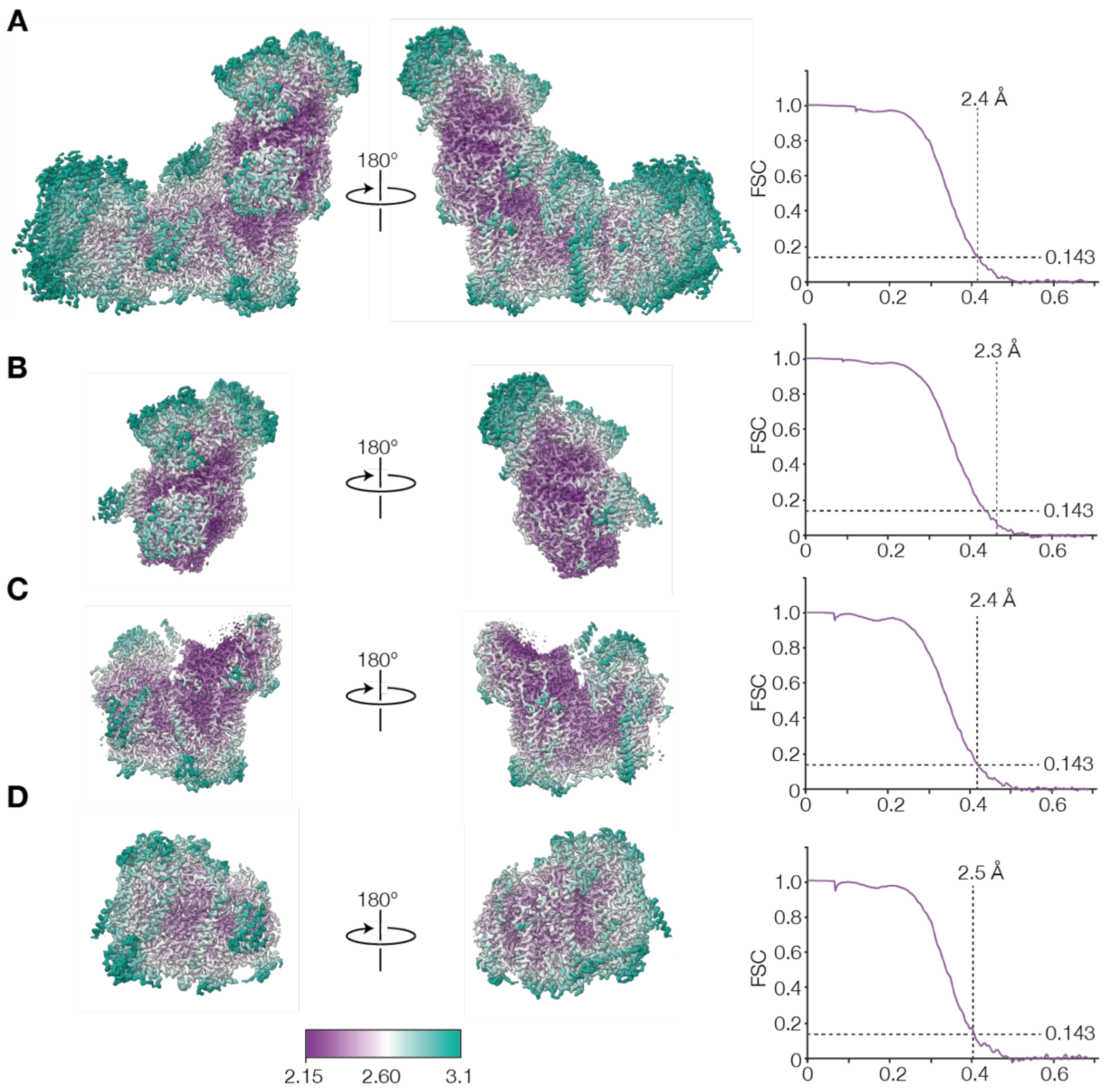


**Fig. S16. Local resolution maps for Active-1092-ii and FSC curves for half maps.** A) Consensus, B) hydrophilic domain, C) proximal hydrophobic domain, D) distal hydrophobic domain.

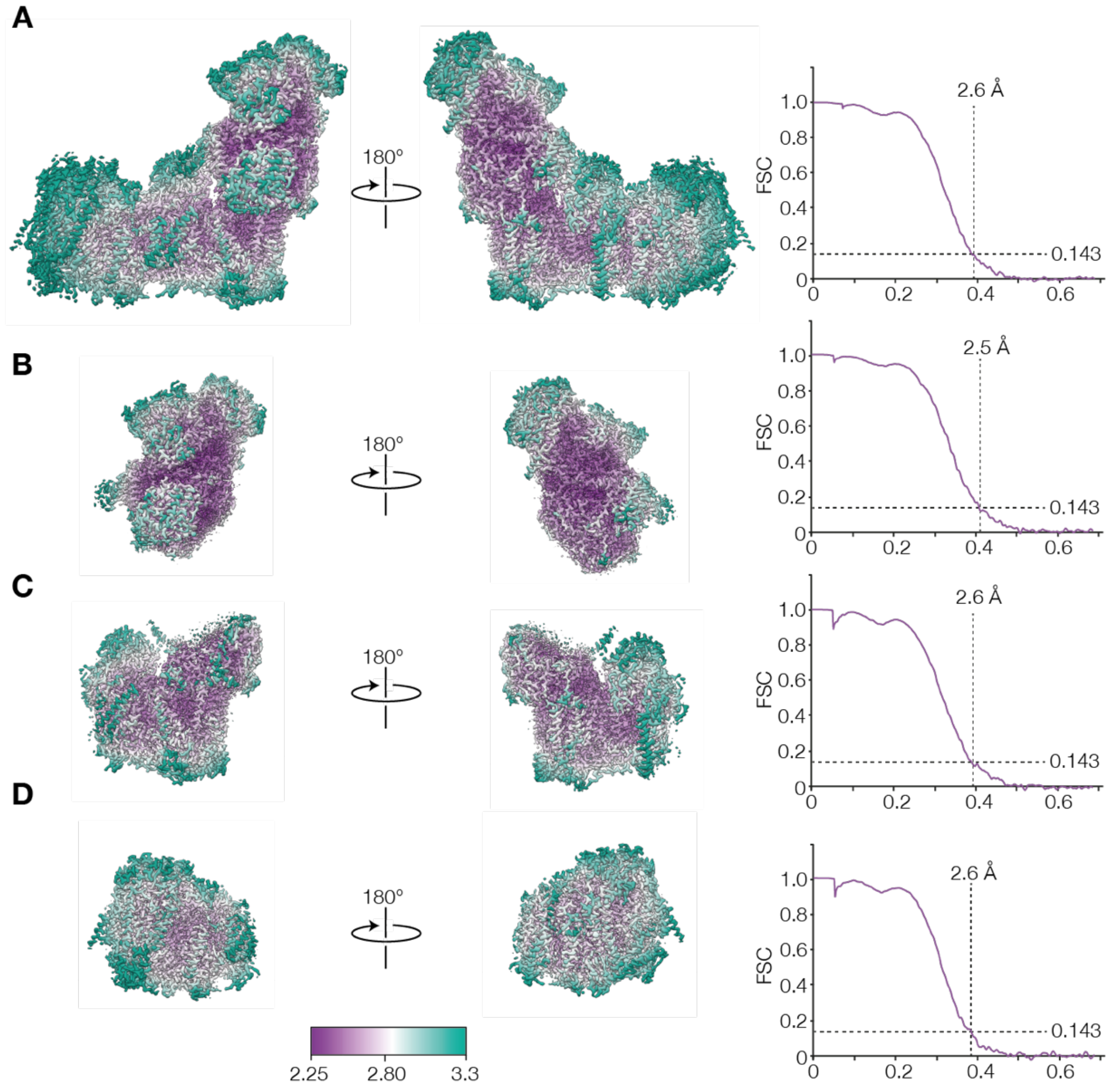


**Fig. S17. Local resolution maps for Active-1092-iii and FSC curves for half maps.** A) Consensus, B) hydrophilic domain, C) proximal hydrophobic domain, D) distal hydrophobic domain.

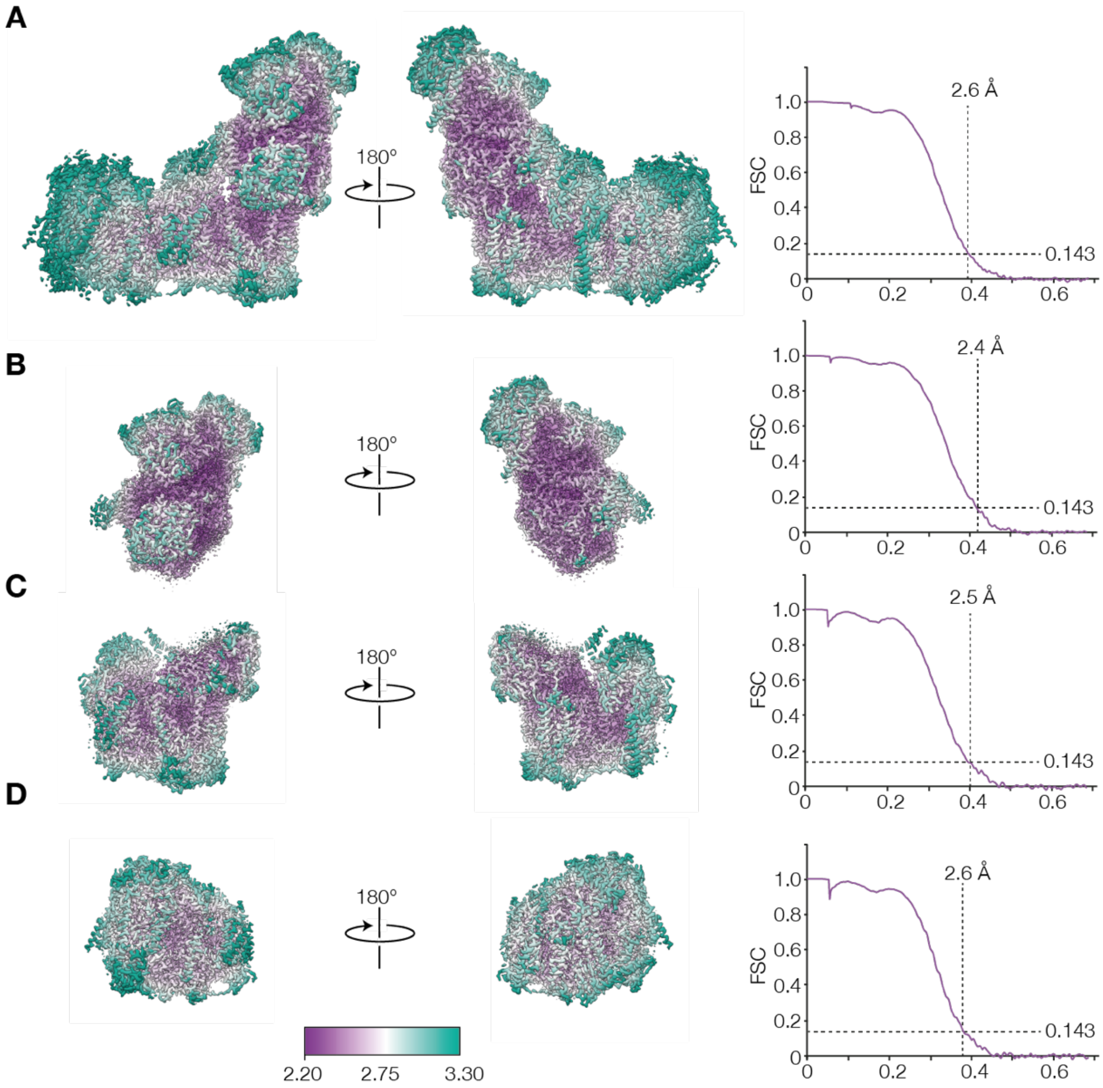


**Fig. S18. Local resolution maps for Active-1092-iv and FSC curves for half maps.** A) Consensus, B) hydrophilic domain, C) proximal hydrophobic domain, D) distal hydrophobic domain.

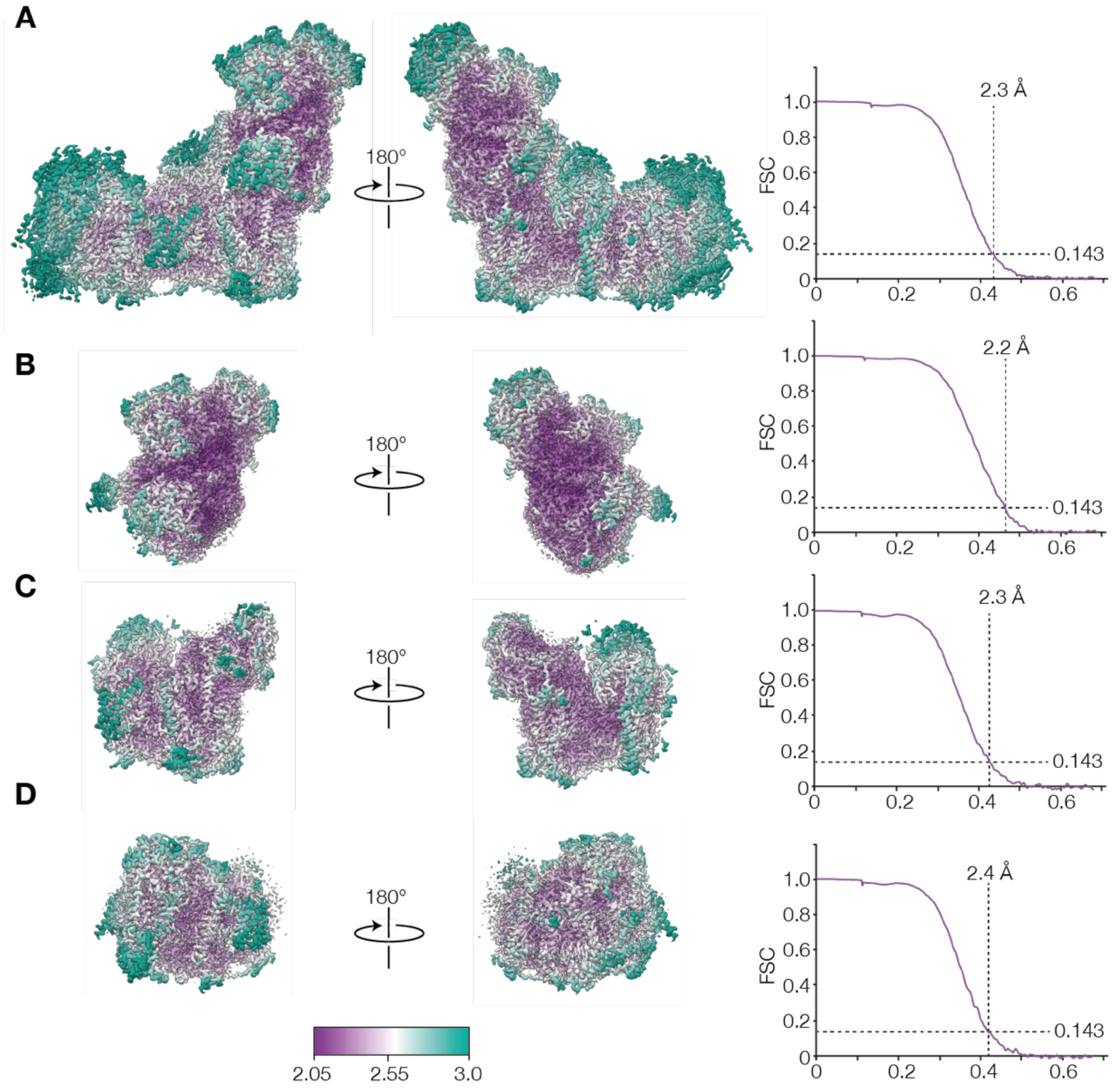


**Fig. S19. Local resolution maps for Deactive-1092-iv and FSC curves for half maps.** A) Consensus, B) hydrophilic domain, C) proximal hydrophobic domain, D) distal hydrophobic domain.

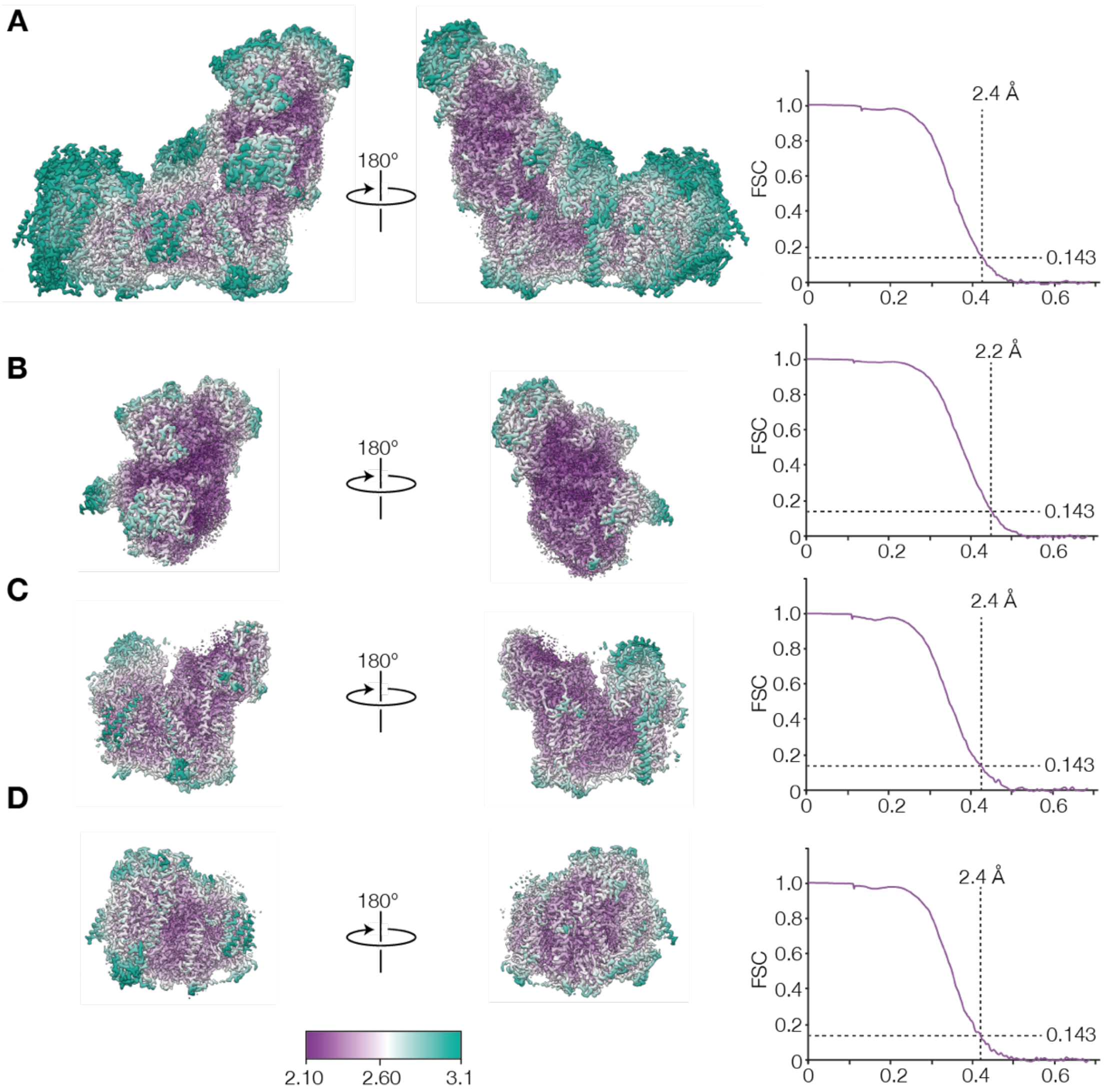


**Fig. S20. Local resolution maps for Deactive-1092-v and FSC curves for half maps.** A) Consensus, B) hydrophilic domain, C) proximal hydrophobic domain, D) distal hydrophobic domain.

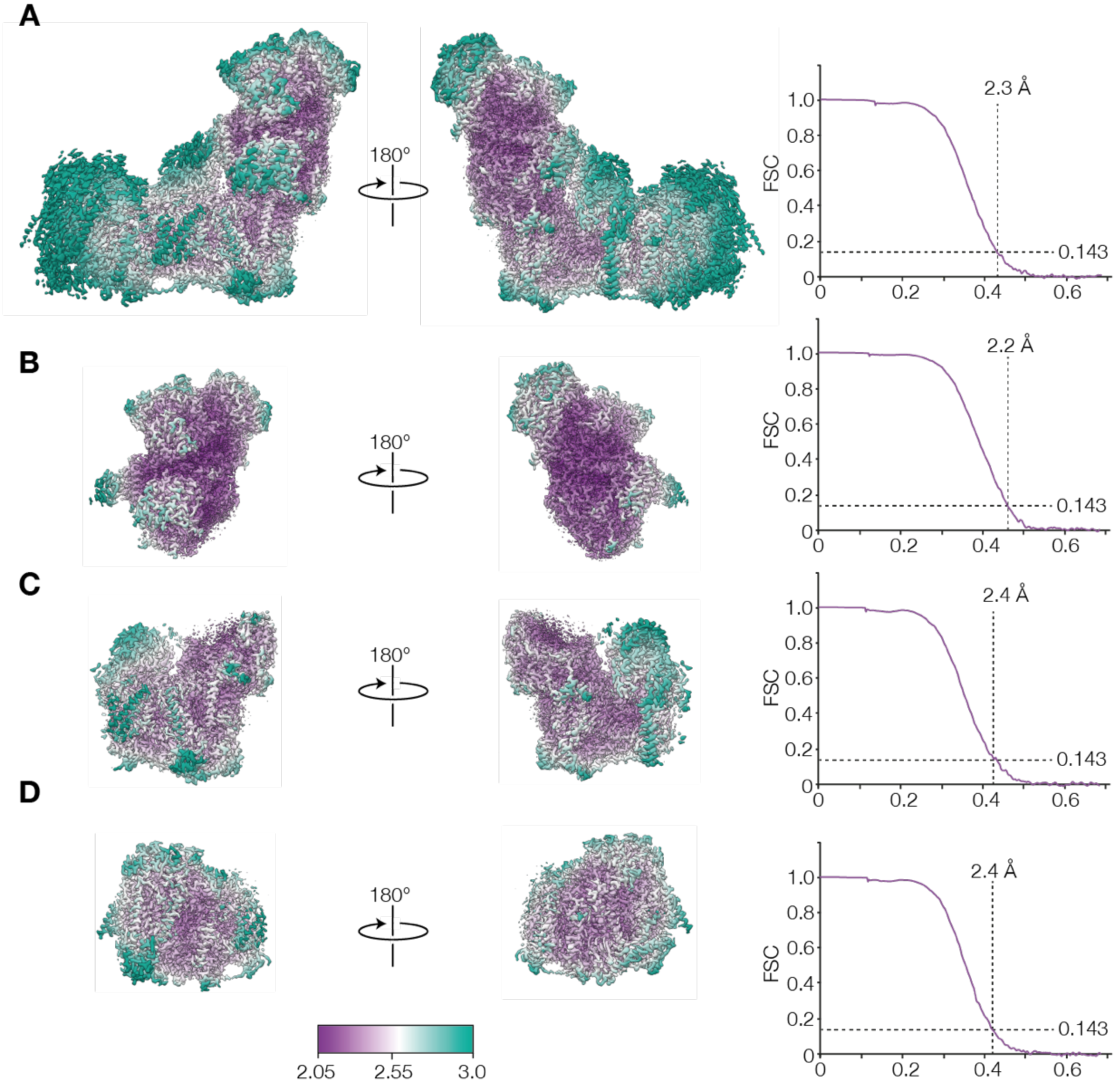


**Fig. S21. Local resolution maps for Deactive-1092-vi and FSC curves for half maps.** A) Consensus, B) hydrophilic domain, C) proximal hydrophobic domain, D) distal hydrophobic domain.

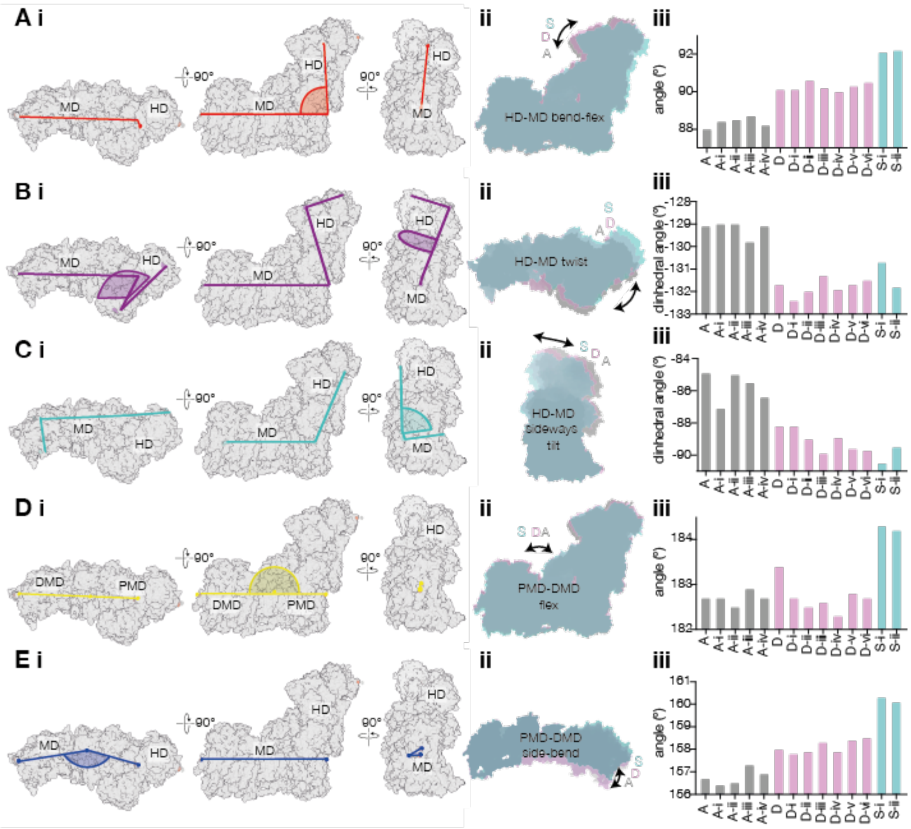

**Fig. S22. Definition of angles and dihedrals used to describe global motions between domains.** A-E i) Orthogonal views are shown for Active-1092-ii. A-D ii) Semi-transparent overlays of an active state (Active-1092-ii) (gray), a deactive state (Deactive-1092-iv) (pink) and a slack state (Slack-1092-ii) aligned to subunit ND1 (mint). E ii) Semi-transparent overlay of the hydrophobic domain core subunits of Active-1092-ii, Deactive-1092-iv and Slack-ii aligned to subunit ND4. Arrows indicate approximate relative domain positions. A) Hydrophilic domain-membrane domain (HD-MD) flex angle between Cα for NDUFB3-Ala75, NDUFA1-Val14 and NDUFS1-Glu165. B) Hydrophilic domain-membrane domain (HD-MD) twist dihedral between Cα for NDUFB3-Ala75, NDUFA1-Val14, NDUFS1-Leu651 and NDUFV1-Ser38. C) Hydrophilic domain-membrane domain (HD-MD) sideways tilt dihedral between Cα for NDUFB8-Phe111, ND5-Tyr42, NDUFA13-Leu44 and NDUFV1-Asp292. D) Proximal membrane domain-distal membrane domain (PMD-DMD) flex angle between the Cα atoms for NDUFB3-Ala75, ND2-Gln168 and NDUFA1-Val14. E) Proximal membrane domain-distal membrane domain (PMD-DMD) side bend angle between the Cα atoms for NDUFB3-Ala75, ND2-Met211 and NDUFA1-Val14. A-E iii) angles measured using PyMOL; A, active 7QSD; A-I, active-1092-I; A-ii, active-1092-ii; A-iii, active-1092-iii; A-iv, bactive-1092-iv; D, Deactive 5O3; D-ii, deactive-1092-I; D-ii, deactive-1092-ii; D-iii, deactive-1092-iii; D-iv, deactive-1092-iv; D-v, deactive-1092-v; D-vi, deactive-1092-vi; S-I, slack-1092-I; S-ii, Slack-1092-ii; For A-E iii) grey; active, pink; deactive, teal; slack.

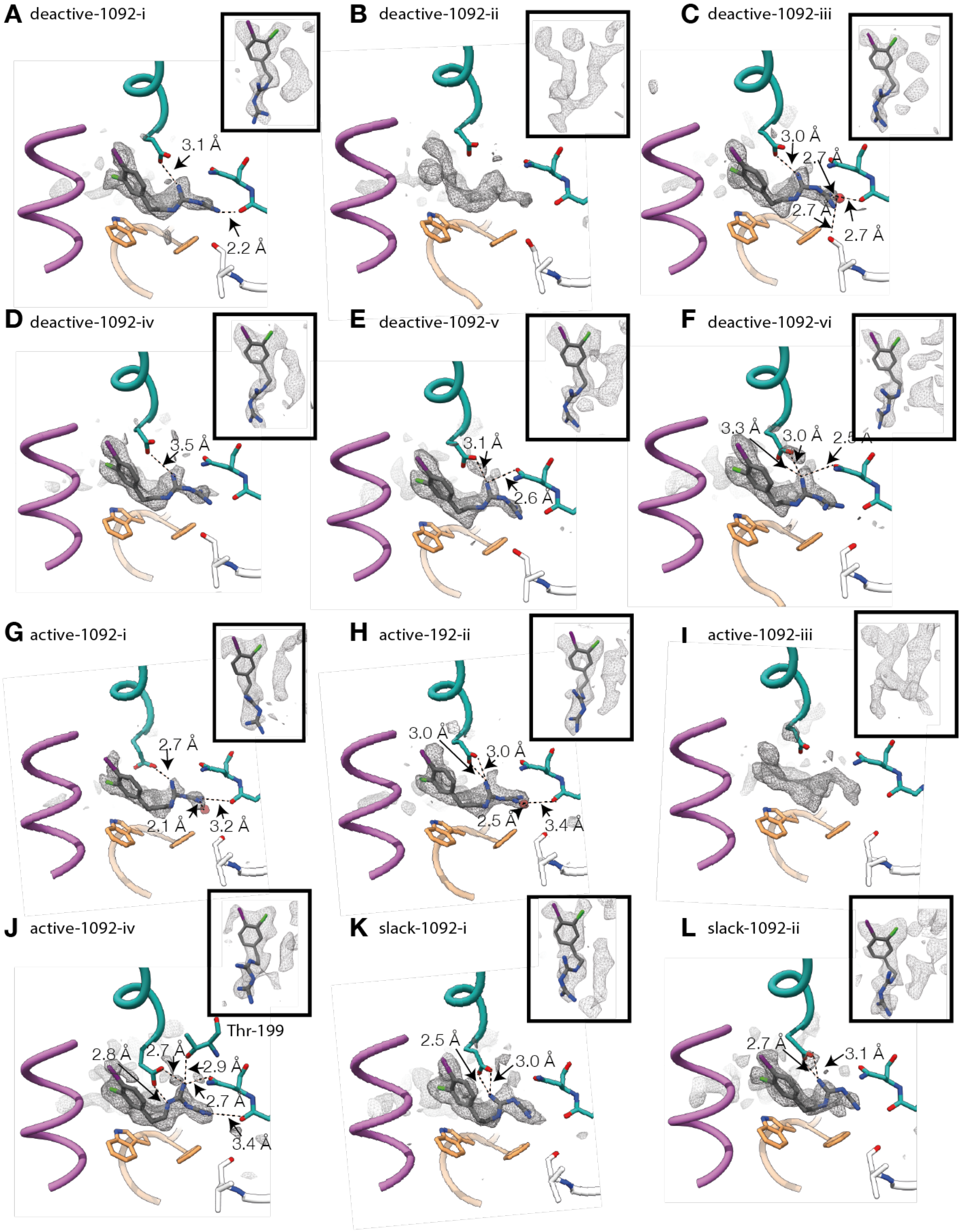


**Fig. S23. The biguanide binding site at the ND2/NDUFB5/NDUFA11 interface.** A) Deactive-1092-i, B) Deactive-1092-ii, C) Deactive-1092-iii, D) Deactive-1092-iv, E) Deactive-1092-v, F) Deactive-1092-vi, G) Active-1092-i, H) Active-1092-ii, I) Active-1092-iii, J) Active-1092-iv, K) Slack-1092-i and L) Slack-1092-ii. Teal, ND2; orchid, NDUFC2; orange, NDUFB5, white, NDUFA11. Model orientation is the same as Fig. 4B. Densities are from the difference maps between the biguanide-free model and experimental densities in the consensus globally sharpened maps. Orientation is the same as labelled in Fig. 4.

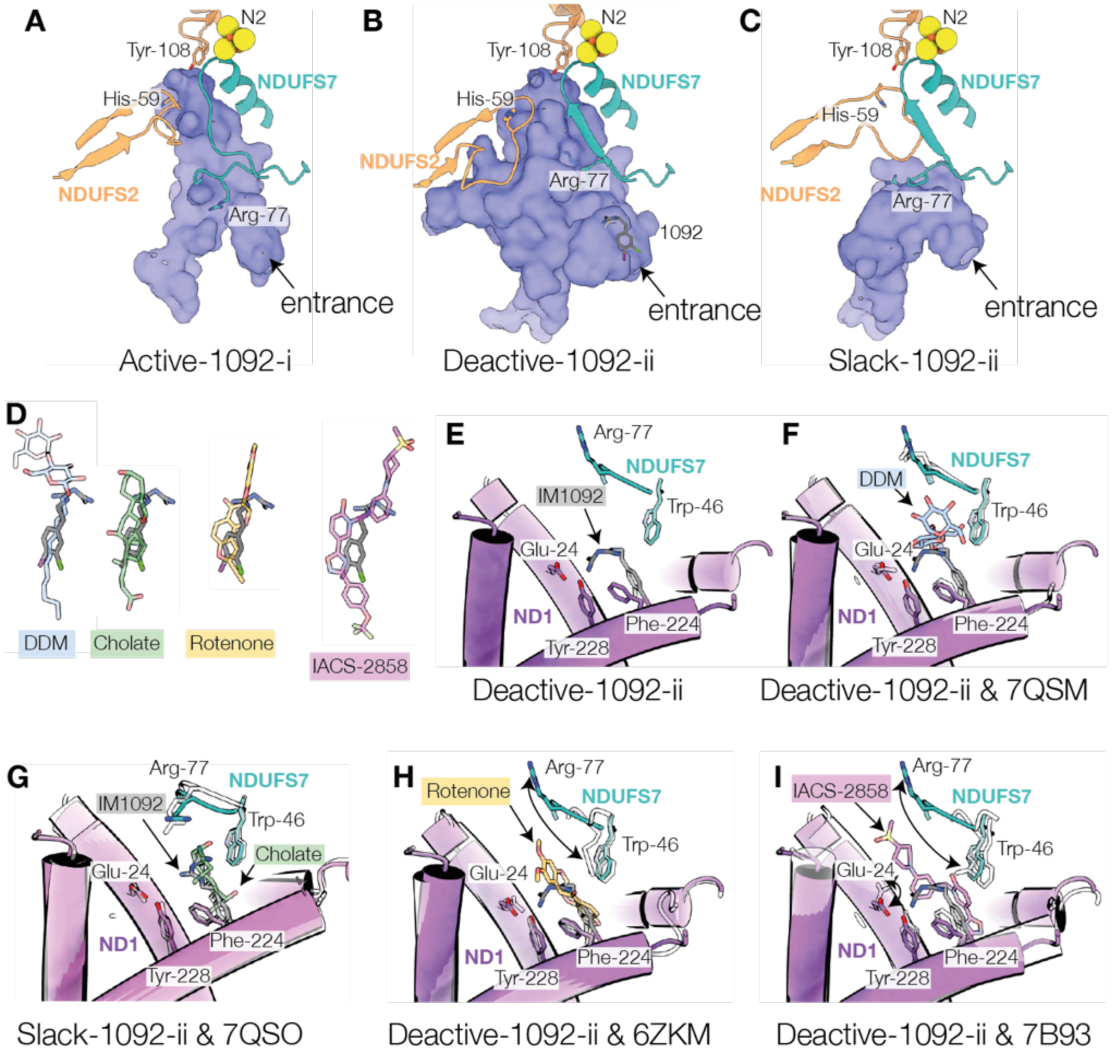


**Fig. S24. Variation in Q-channel cavity between the active, deactive and slack states with ordered β1–β2 loop, and overlap between bound IM1092 and other bound inhibitors.** A-C) Ordered structures for the NDUFS2 β1–β2 loop and a section of NDUFS7 in the A) Active-1092-i, B) Deactive-1092-i and C) Slack-1092-ii states. Blue, cavity predicted by CASTp (*25*) with probe radius of 1.4 Å; orange, NDUFS2; green, NDUFS7. D) Comparison of the binding position of DDM (pale blue), cholate (green), rotenone (yellow) and IACS-2858 (pink) with that of IM1092 (gray) near the Q-channel entrance. E-I) PDB models aligned to the ND1 subunit with Deactive-1092-ii or Slack-1092-ii models colored purple for ND1 and teal for NDUFS2, and overlayed models in white transparent with colored ligands. E) Local environment of IM1092 binding in the same orientation as shown in F-I. F) Comparison to DDM bound in deactive bovine complex I, PDB 7QSM. G) Comparison to cholate bound in slack bovine complex I, PDB 7QSO. H) Comparison to rotenone bound in the open2 state of ovine complex I, PDB 6ZKM. I) Comparison to IACS-2858 bound in the active state of mouse complex I, PDB 7B93.

| | | Active inhibitor-free | Deactive inhibitor-free | Slack inhibitor-free | ND6 TMH3 | ND4 TMH6 | NDUFA11 density | ND3 (31-46) density |
|---|---|---|---|---|---|---|---|---|
| | EMD | 14128 | 3731 | 14126 | | | | |
| Active-1092-i | 14252 | 99% | 84-89% | 80-85% | α | α | good | good |
| Active-1092-ii | 14257 | 99% | 82-89% | 80-84% | α | α | good | good |
| Active-1092-iii | 14262 | 99-100% | 84-88% | 80-84% | α | α | good | good |
| Active-1092-iv | 14267 | 99% | 84-89% | 80-84% | α | α | good | good |
| Deactive-1092-i | 14273 | 79-83% | 94-96% | 91% | π | α | good | poor |
| Deactive-1092-ii | 14278 | 78-82% | 95-96% | 80-94% | π | α | good | poor |
| Deactive-1092-iii | 14283 | 80-83% | 94-96% | 94-95% | π | α | good | poor |
| Deactive-1092-iv | 14288 | 79-83% | 94-96% | 93-94% | π | α | good | poor |
| Deactive-1092-v | 14293 | 79-83% | 94-96% | 95% | π | α | good | poor |
| Deactive-1092-vi | 14298 | 82-84% | 92-95% | 97% | π | α | good | poor |
| Slack-1092-i | 14303 | 78-83% | 86-90% | 97-98% | π | π | poor | poor |
| Slack-1092-ii | 14308 | 78-83% | 87-90% | 97% | π | π | poor | poor |

**Table S1. Map-map correlations for the maps determined here in comparison to maps of detergent solubilized bovine complex I in the absence of an inhibitor, as well as key map characteristics.** All correlations are for the consensus globally sharpened maps, low-pass filtered to 4.13 Å Values were calculated in Chimera (*15*) by testing both directions using the following density thresholds: Active 0.0193, Deactive 0.16, Slack 0.0172, Active-1092-i 0.0143, Active-1092-ii 0.0175, Active-1092-iii 0.0146, Active-1092-iv 0.0166, Deactive-1092-i 0.0148, Deactive-1092-ii 0.0145, Deactive-1092-iii 0.0146, Deactive-1092-iv 0.0146, Deactive-1092-v 0.0144, Deactive-1092-vi 0.0142, Slack-1092-i 0.0115, Slack-1092-ii 0.0112. Shaded boxes show fits of 95% and above. All fits were made with the hydrophilic domain aligned. Deactive-1092-iii, iv, v vi showed a similar level of correlation to both inhibitor-free deactive and slack states.

| | Deactive-1092-i | | | | |
|---|---|---|---|---|---|
| | Consensus | Hydrophilic domain | Proximal hydrophobic domain | Distal hydrophobic domain | Composite |
| PDB code | | | | | PDB 7R45 |
| EMDB code | EMD-14273 | EMD-14274 | EMD-14275 | EMD-14276 | EMD-14272 |
| **Data collection and processing** | | | | | |
| Magnification (nominal) | 81,000x | | | | |
| Voltage (kV) | 300 | | | | |
| Electron exposure ($e^{-}/Å^2$) | 40 | | | | |
| Defocus range (μm) | -1.0 to -2.4 | | | | |
| Calibrated Pixel size (Å) | 1.072 | | | | |
| Symmetry imposed | none | | | | |
| Initial particle images (no.) | 907,090 | | | | |
| Final particle images (no.) | 68,023 | | | | |
| Map sharpening B factor ($Å^2$) | -37 | -30 | -30 | -32 | |
| Map resolution (FSC=0.143) (Å) | 2.44 | 2.30 | 2.46 | 2.47 | |
| Map resolution D99 | | | | | 2.40 |
| Map angular accuracy (°) | 0.231 | 0.262 | 0.436 | 0.425 | |
| Map translational accuracy(Å) | 0.182 | 0.213 | 0.258 | 0.280 | |
| Map resolution range (Å) | 2.14-6.88 | 2.11-4.69 | 2.27-4.98 | 2.26-5.53 | |
| Final map sampling (Å/pix) | 0.731 | | | | |
| **Model Refinement** | | | | | |
| Initial model used | 7R43 | | | | |
| Model resolution (FSC=0.5) (Å) | | | | | 2.3 |
| Model composition<br>Nonhydrogen atoms<br>Protein residues<br>Waters<br>Ligands | | | | | <br>68195<br>8098<br>1334<br>52 |
| B factors mean ($Å^2$)<br>Protein<br>Ligand<br>Water | | | | | <br>45.57<br>48.50<br>39.45 |
| RMSD Bond lengths (Å)<br>RMSD Bond angles (°) | | | | | 0.006<br>1.032 |
| Validation<br>MolProbity score<br>Clashscore<br>Poor rotamers (%) | | | | | <br>1.19<br>2.72<br>1.20 |
| EMRinger score | 5.31 | | | | 5.82 |
| Ramachandran plot<br>Favored (%)<br>Allowed (%)<br>Disallowed (%)<br>Z-score (whole) | | | | | 97.69<br>2.29<br>0.01<br>0.76 |
| Map-model correlation $CC_{mask}$ | | | | | 0.90 |

**Supplementary Table 2. Cryo-EM data collection, refinement and validation statistics for deactive-1092-i.**

| | Deactive-1092-ii | | | | |
|---|---|---|---|---|---|
| | Consensus | Hydrophilic domain | Proximal hydrophobic domain | Distal hydrophobic domain | Composite |
| PDB code | | | | | PDB 7R46 |
| EMDB code | EMD-14278 | EMD-14279 | EMD-14280 | EMD-14281 | EMD-14277 |
| **Data collection and processing** | | | | | |
| Magnification (nominal) | 81,000 x | | | | |
| Voltage (kV) | 300 | | | | |
| Electron exposure ($e^{-}/Å^2$) | 40 | | | | |
| Defocus range (μm) | -1.0 to -2.4 | | | | |
| Calibrated Pixel size (Å) | 1.072 | | | | |
| Symmetry imposed | none | | | | |
| Initial particle images (no.) | 907,090 | | | | |
| Final particle images (no.) | 35,448 | | | | |
| Map sharpening B factor ($Å^2$) | -35 | -28 | -28 | -32 | |
| Map resolution (FSC=0.143) (Å) | 2.57 | 2.44 | 2.55 | 2.61 | |
| Map resolution D99 | | | | | 2.40 |
| Map angular accuracy (°) | 0.254 | 0.259 | 0.453 | 0.373 | |
| Map translational accuracy(Å) | 0.181 | 0.212 | 0.275 | 0.270 | |
| Map resolution range (Å) | 2.25-7.95 | 2.20-5.32 | 2.42-5.46 | 2.39-6.10 | |
| Final map sampling (Å/pix) | 0.731 | | | | |
| **Model Refinement** | | | | | |
| Initial model used | | | | | |
| Model resolution (FSC=0.5) (Å) | | | | | 2.5 |
| Model composition<br>Nonhydrogen atoms<br>Protein residues<br>Waters<br>Ligands | | | | | <br>67614<br>8100<br>859<br>51 |
| B factors mean ($Å^2$)<br>Protein<br>Ligand<br>Water | | | | | <br>46.28<br>49.63<br>38.67 |
| RMSD Bond lengths (Å)<br>RMSD Bond angles (°) | | | | | 0.006<br>1.017 |
| Validation<br>MolProbity score<br>Clashscore<br>Poor rotamers (%) | | | | | 1.29<br>3.34<br>0.00 |
| EMRinger score | 5.12 | | | | 5.72 |
| Ramachandran plot<br>Favored (%)<br>Allowed (%)<br>Disallowed (%)<br>Z-score (whole) | | | | | <br>97.07<br>2.91<br>0.03<br>0.30 |
| Map-model correlation $CC_{mask}$ | | | | | 0.89 |

**Supplementary Table 3. Cryo-EM data collection, refinement and validation statistics for deactive-1092-ii.**

| | Deactive-1092-iii | | | | |
|---|---|---|---|---|---|
| | Consensus | Hydrophilic domain | Proximal hydrophobic domain | Distal hydrophobic domain | Composite |
| PDB code | | | | | PDB 7R47 |
| EMDB code | EMD-14283 | EMD-14284 | EMD-14285 | EMD-14286 | EMD-14282 |
| **Data collection and processing** | | | | | |
| Magnification (nominal) | 81,000 x | | | | |
| Voltage (kV) | 300 | | | | |
| Electron exposure ($e^{-}/Å^{2}$) | 40 | | | | |
| Defocus range (µm) | -1.0 to -2.4 | | | | |
| Calibrated Pixel size (Å) | 1.072 | | | | |
| Symmetry imposed | none | | | | |
| Initial particle images (no.) | 907,090 | | | | |
| Final particle images (no.) | 123,012 | | | | |
| Map sharpening B factor ($Å^2$) | -42 | -33 | -35 | -38 | |
| Map resolution (FSC=0.143) (Å) | 2.31 | 2.17 | 2.29 | 2.34 | |
| Map resolution D99 | | | | | 2.30 |
| Map angular accuracy (°) | 0.218 | 0.243 | 0.406 | 0.319 | |
| Map translational accuracy(Å) | 0.171 | 0.202 | 0.251 | 0.239 | |
| Map resolution range (Å) | 2.05-6.16 | 2.02-4.23 | 2.14-4.43 | 2.16-4.89 | |
| Final map sampling (Å/pix) | 0.731 | | | | |
| **Model Refinement** | | | | | |
| Initial model used | | | | | |
| Model resolution (FSC=0.5) (Å) | | | | | 2.2 |
| Model composition<br>Nonhydrogen atoms<br>Protein residues<br>Waters<br>Ligands | | | | | <br>68578<br>8089<br>1836<br>51 |
| B factors mean ($Å^2$)<br>Protein<br>Ligand<br>Water | | | | | <br>40.74<br>43.28<br>36.02 |
| RMSD Bond lengths (Å)<br>RMSD Bond angles (°) | | | | | 0.006<br>1.024 |
| Validation<br>MolProbity score<br>Clashscore<br>Poor rotamers (%) | | | | | <br>1.12<br>2.53<br>1.18 |
| EMRinger score | 5.58 | | | | 6.13 |
| Ramachandran plot<br>Favored (%)<br>Allowed (%)<br>Disallowed (%)<br>Z-score (whole) | | | | | <br>97.88<br>2.10<br>0.03<br>0.85 |
| Map-model correlation $CC_{mask}$ | | | | | 0.90 |

**Supplementary Table 4. Cryo-EM data collection, refinement and validation statistics for deactive-1092-iii.**

| | Deactive-1092-iv | | | | |
|---|---|---|---|---|---|
| | Consensus | Hydrophilic domain | Proximal hydrophobic domain | Distal hydrophobic domain | Composite |
| PDB code | | | | | PDB 7R48 |
| EMDB code | EMD-14288 | EMD-14289 | EMD-14290 | EMD-14291 | EMD-14287 |
| **Data collection and processing** | | | | | |
| Magnification (nominal) | 81,000 x | | | | |
| Voltage (kV) | 300 | | | | |
| Electron exposure ($e^{-}$/Å$^2$) | 40 | | | | |
| Defocus range (μm) | -1.0 to -2.4 | | | | |
| Calibrated Pixel size (Å) | 1.072 | | | | |
| Symmetry imposed | none | | | | |
| Initial particle images (no.) | 907,090 | | | | |
| Final particle images (no.) | 111,066 | | | | |
| Map sharpening B factor (Å$^2$) | -40 | -33 | -33 | -29 | |
| Map resolution (FSC=0.143) (Å) | 2.34 | 2.19 | 2.34 | 2.38 | |
| Map resolution D99 | | | | | 2.30 |
| Map angular accuracy (°) | 0.241 | 0.238 | 0.402 | 0.345 | |
| Map translational accuracy(Å) | 0.171 | 0.212 | 0.253 | 0.259 | |
| Map resolution range (Å) | 2.10-6.33 | 2.05-4.35 | 2.21-4.63 | 2.20-5.18 | |
| Final map sampling (Å/pix) | 0.713 | | | | |
| **Model Refinement** | | | | | |
| Initial model used | | | | | |
| Model resolution (FSC=0.5) (Å) | | | | | 2.2 |
| Model composition<br>Nonhydrogen atoms<br>Protein residues<br>Waters<br>Ligands | | | | | <br>69247<br>8118<br>2124<br>55 |
| B factors mean (Å$^2$)<br>Protein<br>Ligand<br>Water | | | | | 41.74<br>45.38<br>37.65 |
| RMSD Bond lengths (Å)<br>RMSD Bond angles (°) | | | | | 0.007<br>1.036 |
| Validation<br>MolProbity score<br>Clashscore<br>Poor rotamers (%) | | | | | <br>1.14<br>2.50<br>1.26 |
| EMRinger score | 5.33 | | | | 6.11 |
| Ramachandran plot<br>Favored (%)<br>Allowed (%)<br>Disallowed (%)<br>Z-score (whole) | | | | | <br>97.86<br>2.11<br>0.03<br>0.80 |
| Map-model correlation $CC_{mask}$ | | | | | 0.91 |

**Supplementary Table 5. Cryo-EM data collection, refinement and validation statistics for deactive-1092-iv**

| | Deactive-1092-v | | | | |
|---|---|---|---|---|---|
| | Consensus | Hydrophilic domain | Proximal hydrophobic domain | Distal hydrophobic domain | Composite |
| PDB code | | | | | PDB 7R4C |
| EMDB code | EMD-14293 | EMD-14294 | EMD-14295 | EMD-14296 | EMD-14292 |
| **Data collection and processing** | | | | | |
| Magnification (nominal) | 81,000 x | | | | |
| Voltage (kV) | 300 | | | | |
| Electron exposure ($e^{-}/Å^2$) | 40 | | | | |
| Defocus range (μm) | -1.0 to -2.4 | | | | |
| Calibrated Pixel size (Å) | 1.072 | | | | |
| Symmetry imposed | none | | | | |
| Initial particle images (no.) | 907,090 | | | | |
| Final particle images (no.) | 91,673 | | | | |
| Map sharpening B factor ($Å^2$) | -39 | -31 | -33 | -36 | |
| Map resolution (FSC=0.143) (Å) | 2.38 | 2.22 | 2.38 | 2.40 | |
| Map resolution D99 | | | | | 2.30 |
| Map angular accuracy (°) | 0.232 | 0.248 | 0.393 | 0.363 | |
| Map translational accuracy(Å) | 0.176 | 0.212 | 0.265 | 0.258 | |
| Map resolution range (Å) | 2.11-6.88 | 2.07-4.45 | 2.22-4.84 | 2.22-5.39 | |
| Final map sampling (Å/pix) | 0.713 | | | | |
| **Model Refinement** | | | | | |
| Initial model used | | | | | |
| Model resolution (FSC=0.5) (Å) | | | | | 2.3 |
| Model composition<br>Nonhydrogen atoms<br>Protein residues<br>Waters<br>Ligands | | | | | <br>68045<br>8069<br>1525<br>51 |
| B factors mean ($Å^2$)<br>Protein<br>Ligand<br>Water | | | | | <br>40.17<br>43.63<br>34.29 |
| RMSD Bond lengths (Å)<br>RMSD Bond angles (°) | | | | | 0.006<br>1.021 |
| Validation<br>MolProbity score<br>Clashscore<br>Poor rotamers (%) | | | | | <br>1.15<br>3.06<br>0.01 |
| EMRinger score | 5.45 | | | | 6.21 |
| Ramachandran plot<br>Favored (%)<br>Allowed (%)<br>Disallowed (%)<br>Z-score (whole) | | | | | <br>97.75<br>2.23<br>0.03<br>0.69 |
| Map-model correlation $CC_{mask}$ | | | | | 0.90 |

**Supplementary Table 6. Cryo-EM data collection, refinement and validation statistics for dactive-1092-v**

| | Deactive-1092-vi | | | | |
|---|---|---|---|---|---|
| | Consensus | Hydrophilic domain | Proximal hydrophobic domain | Distal hydrophobic domain | Composite |
| PDB code | | | | | PDB 7R4D |
| EMDB code | EMD-14298 | EMD-14299 | EMD-14300 | ED-14301 | EMD-14297 |
| **Data collection and processing** | | | | | |
| Magnification (nominal) | 81,000 x | | | | |
| Voltage (kV) | 300 | | | | |
| Electron exposure ($e^{-}/Å^2$) | 40 | | | | |
| Defocus range (μm) | -1.0 to -2.4 | | | | |
| Calibrated Pixel size (Å) | 1.072 | | | | |
| Symmetry imposed | none | | | | |
| Initial particle images (no.) | 907,090 | | | | |
| Final particle images (no.) | 122,019 | | | | |
| Map sharpening B factor ($Å^2$) | -40 | -34 | -35 | -38 | |
| Map resolution (FSC=0.143) (Å) | 2.33 | 2.19 | 2.35 | 2.40 | |
| Map resolution D99 | | | | | 2.30 |
| Map angular accuracy (°) | 0.236 | 0.256 | 0.455 | 0.345 | |
| Map translational accuracy(Å) | 0.189 | 0.208 | 0.278 | 0.254 | |
| Map resolution range (Å) | 2.08-6.71 | 2.04-4.45 | 2.20-4.75 | 2.21-5.42 | |
| Final map sampling (Å/pix) | 0.731 | | | | |
| **Model Refinement** | | | | | |
| Initial model used | | | | | |
| Model resolution (FSC=0.5) (Å) | | | | | 2.3 |
| Model composition<br>Nonhydrogen atoms<br>Protein residues<br>Waters<br>Ligands | | | | | <br>68004<br>8056<br>1607<br>49 |
| B factors mean ($Å^2$)<br>Protein<br>Ligand<br>Water | | | | | <br>21.80<br>21.77<br>19.80 |
| RMSD Bond lengths (Å)<br>RMSD Bond angles (°) | | | | | 0.006<br>1.053 |
| Validation<br>MolProbity score<br>Clashscore<br>Poor rotamers (%) | | | | | 1.15<br>3.32<br>0.03 |
| EMRinger score | 5.15 | | | | 6.23 |
| Ramachandran plot<br>Favored (%)<br>Allowed (%)<br>Disallowed (%)<br>Z-score (whole) | | | | | <br>97.91<br>2.08<br>0.01<br>0.61 |
| Map-model correlation $CC_{mask}$ | | | | | 0.89 |

**Supplementary Table 7. Cryo-EM data collection, refinement and validation statistics for deactive-1092-vi.**

| | Active-1092-i | | | | |
|---|---|---|---|---|---|
| | Consensus | Hydrophilic domain | Proximal hydrophobic domain | Distal hydrophobic domain | Composite |
| PDB code | | | | | PDB 7R41 |
| EMDB code | EMD-14252 | EMD-14253 | EMD-14254 | EMD-14255 | EMD-14251 |
| **Data collection and processing** | | | | | |
| Magnification (nominal) | 81,000 x | | | | |
| Voltage (kV) | 300 | | | | |
| Electron exposure ($e^{-}/Å^2$) | 40 | | | | |
| Defocus range (μm) | -1.0 to -2.4 | | | | |
| Calibrated Pixel size (Å) | 1.072 | | | | |
| Symmetry imposed | none | | | | |
| Initial particle images (no.) | 907,090 | | | | |
| Final particle images (no.) | 48,367 | | | | |
| Map sharpening B factor ($Å^2$) | -23 | -21 | -21 | -24 | |
| Map resolution (FSC=0.143) (Å) | 2.44 | 2.32 | 2.41 | 2.51 | |
| Map resolution D99 | | | | | 2.30 |
| Map angular accuracy (°) | 0.224 | 0.246 | 0.389 | 0.374 | |
| Map translational accuracy(Å) | 0.166 | 0.219 | 0.246 | 0.267 | |
| Map resolution range (Å) | 2.16-7.93 | 2.14-5.79 | 2.25-5.40 | 2.35-5.51 | |
| Final map sampling (Å/pix) | 0.731 | | | | |
| **Model Refinement** | | | | | |
| Initial model used | | | | | |
| Model resolution (FSC=0.5) (Å) | | | | | 2.4 |
| Model composition<br>Nonhydrogen atoms<br>Protein residues<br>Waters<br>Ligands | | | | | <br>69138<br>8194<br>1307<br>57 |
| B factors mean ($Å^2$)<br>Protein<br>Ligand<br>Water | | | | | <br>45.95<br>49.51<br>38.88 |
| RMSD Bond lengths (Å)<br>RMSD Bond angles (°) | | | | | 0.007<br>1.025 |
| Validation<br>MolProbity score<br>Clashscore<br>Poor rotamers (%) | | | | | <br>1.22<br>3.27<br>1.01 |
| EMRinger score | 5.46 | | | | 5.72 |
| Ramachandran plot<br>Favored (%)<br>Allowed (%)<br>Disallowed (%)<br>Z-score (whole) | | | | | <br>97.54<br>2.44<br>0.02<br>0.57 |
| Map-model correlation $CC_{mask}$ | | | | | 0.88 |

**Supplementary Table 8. Cryo-EM data collection, refinement and validation statistics for active-1092-i.**

| | Active-1092-ii | | | | |
|---|---|---|---|---|---|
| | Consensus | Hydrophilic domain | Proximal hydrophobic domain | Distal hydrophobic domain | Composite |
| PDB code | | | | | PDB 7R42 |
| EMDB code | EMD-14257 | EMD-14258 | EMD-14259 | EMD-14260 | EMD-14256 |
| **Data collection and processing** | | | | | |
| Magnification (nominal) | 81,000 x | | | | |
| Voltage (kV) | 300 | | | | |
| Electron exposure ($e^{-}/Å^2$) | 40 | | | | |
| Defocus range (μm) | -1.0 to -2.4 | | | | |
| Calibrated Pixel size (Å) | 1.072 | | | | |
| Symmetry imposed | none | | | | |
| Initial particle images (no.) | 907,090 | | | | |
| Final particle images (no.) | 53,763 | | | | |
| Map sharpening B factor ($Å^2$) | -29 | -22 | -23 | -27 | |
| Map resolution (FSC=0.143) (Å) | 2.42 | 2.31 | 2.41 | 2.47 | |
| Map resolution D99 | | | | | 2.30 |
| Map angular accuracy (°) | 0.224 | 0.257 | 0.389 | 0.368 | |
| Map translational accuracy(Å) | 0.123 | 0.212 | 0.249 | 0.246 | |
| Map resolution range (Å) | 2.15-7.79 | 2.13-4.75 | 2.22-5.20 | 2.30-5.41 | |
| Final map sampling (Å/pix) | 0.731 | | | | |
| **Model Refinement** | | | | | |
| Initial model used | | | | | |
| Model resolution (FSC=0.5) (Å) | | | | | 2.3 |
| Model composition<br>Nonhydrogen atoms<br>Protein residues<br>Waters<br>Ligands | | | | | <br>69480<br>8204<br>1545<br>58 |
| B factors mean ($Å^2$)<br>Protein<br>Ligand<br>Water | | | | | <br>42.35<br>44.62<br>37.40 |
| RMSD Bond lengths (Å)<br>RMSD Bond angles (°) | | | | | 0.006<br>1.026 |
| Validation<br>MolProbity score<br>Clashscore<br>Poor rotamers (%) | | | | | <br>1.27<br>3.03<br>1.20 |
| EMRinger score | 5.59 | | | | 5.95 |
| Ramachandran plot<br>Favored (%)<br>Allowed (%)<br>Disallowed (%)<br>Z-score (whole) | | | | | <br>97.42<br>2.57<br>0.01<br>0.55 |
| Map-model correlation $CC_{mask}$ | | | | | 0.90 |

**Supplementary Table 9. Cryo-EM data collection, refinement and validation statistics for active-1092-ii.**

| | Active-1092-iii | | | | |
|---|---|---|---|---|---|
| | Consensus | Hydrophilic domain | Proximal hydrophobic domain | Distal hydrophobic domain | Composite |
| PDB code | | | | | PDB 7R43 |
| EMDB code | EMD-14262 | EMD-14263 | EMD-14264 | EMD-14265 | EMD-14261 |
| **Data collection and processing** | | | | | |
| Magnification (nominal) | 81,000 x | | | | |
| Voltage (kV) | 300 | | | | |
| Electron exposure ($e^{-}/Å^2$) | 40 | | | | |
| Defocus range (μm) | -1.0 to -2.4 | | | | |
| Calibrated Pixel size (Å) | 1.072 | | | | |
| Symmetry imposed | none | | | | |
| Initial particle images (no.) | 907,090 | | | | |
| Final particle images (no.) | 27,326 | | | | |
| Map sharpening B factor ($Å^2$) | -25 | -18 | -20 | -23 | |
| Map resolution (FSC=0.143) (Å) | 2.61 | 2.46 | 2.58 | 2.62 | |
| Map resolution D99 | | | | | 2.40 |
| Map angular accuracy (°) | 0.216 | 0.278 | 0.423 | 0.438 | |
| Map translational accuracy(Å) | 0.173 | 0.217 | 0.252 | 0.282 | |
| Map resolution range (Å) | 2.26-8.62 | 2.23-6.62 | 2.36-6.38 | 2.52-6.67 | |
| Final map sampling (Å/pix) | 0.731 | | | | |
| **Model Refinement** | | | | | |
| Initial model used | | | | | |
| Model resolution (FSC=0.5) (Å) | | | | | 2.4 |
| Model composition<br>Nonhydrogen atoms<br>Protein residues<br>Waters<br>Ligands | | | | | <br>68334<br>8188<br>885<br>47 |
| B factors mean ($Å^2$)<br>Protein<br>Ligand<br>Water | | | | | <br>45.58<br>47.91<br>39.06 |
| RMSD Bond lengths (Å)<br>RMSD Bond angles (°) | | | | | 0.006<br>1.013 |
| Validation<br>MolProbity score<br>Clashscore<br>Poor rotamers (%) | | | | | <br>1.24<br>3.33<br>0.01 |
| EMRinger score | 5.25 | | | | 5.73 |
| Ramachandran plot<br>Favored (%)<br>Allowed (%)<br>Disallowed (%)<br>Z-score (whole) | | | | | <br>97.44<br>2.54<br>0.02<br>0.39 |
| Map-model correlation $CC_{mask}$ | | | | | 0.89 |

**Supplementary Table 10. Cryo-EM data collection, refinement and validation statistics for active-1092-iii.**

| | Active-1092-iv | | | | |
|---|---|---|---|---|---|
| | Consensus | Hydrophilic domain | Proximal hydrophobic domain | Distal hydrophobic domain | Composite |
| PDB code | | | | | PDB 7R44 |
| EMDB code | EMD-14267 | EMD-14268 | EMD-14269 | EMD-14270 | EMD-14266 |
| **Data collection and processing** | | | | | |
| Magnification (nominal) | 81,000 x | | | | |
| Voltage (kV) | 300 | | | | |
| Electron exposure ($e^{-}/Å^2$) | 40 | | | | |
| Defocus range (μm) | -1.0 to -2.4 | | | | |
| Calibrated Pixel size (Å) | 1.072 | | | | |
| Symmetry imposed | none | | | | |
| Initial particle images (no.) | 907,090 | | | | |
| Final particle images (no.) | 31,957 | | | | |
| Map sharpening B factor ($Å^2$) | -26 | -19 | -20 | -24 | |
| Map resolution (FSC=0.143) (Å) | 2.55 | 2.41 | 2.54 | 2.64 | |
| Map resolution D99 | | | | | 2.40 |
| Map angular accuracy (°) | 0.227 | 0.260 | 0.410 | 0.39 | |
| Map translational accuracy(Å) | 0.171 | 0.223 | 0.263 | 0.278 | |
| Map resolution range (Å) | 2.24-8.62 | 2.20-6.62 | 2.36-6.38 | 2.52-6.67 | |
| Final map sampling (Å/pix) | 0.731 | | | | |
| **Model Refinement** | | | | | |
| Initial model used | | | | | |
| Model resolution (FSC=0.5) (Å) | | | | | 2.4 |
| Model composition<br>Nonhydrogen atoms<br>Protein residues<br>Waters<br>Ligands | | | | | <br>68213<br>8163<br>905<br>50 |
| B factors mean ($Å^2$)<br>Protein<br>Ligand<br>Water | | | | | <br>45.07<br>47.48<br>38.42 |
| RMSD Bond lengths (Å)<br>RMSD Bond angles (°) | | | | | 0.006<br>1.018 |
| Validation<br>MolProbity score<br>Clashscore<br>Poor rotamers (%) | | | | | <br>1.22<br>3.18<br>0.00 |
| EMRinger score | 5.50 | | | | 6.05 |
| Ramachandran plot<br>Favored (%)<br>Allowed (%)<br>Disallowed (%)<br>Z-score (whole) | | | | | <br>97.47<br>2.52<br>0.01<br>0.40 |
| Map-model correlation $CC_{mask}$ | | | | | 0.89 |

**Supplementary Table 11. Cryo-EM data collection, refinement and validation statistics for active-1092-iv.**

| | Slack-1092-i | | | | |
|---|---|---|---|---|---|
| | Consensus | Hydrophilic domain | Proximal hydrophobic domain | Distal hydrophobic domain | Composite |
| PDB code | | | | | PDB 7R4F |
| EMDB code | EMD-14303 | EMD-14304 | EMD-14305 | EMD-14306 | EMD-14302 |
| **Data collection and processing** | | | | | |
| Magnification (nominal) | 81,000 x | | | | |
| Voltage (kV) | 300 | | | | |
| Electron exposure ($e^{-}/Å^2$) | 40 | | | | |
| Defocus range (μm) | -1.0 to -2.4 | | | | |
| Calibrated Pixel size (Å) | 1.072 | | | | |
| Symmetry imposed | none | | | | |
| Initial particle images (no.) | 907,090 | | | | |
| Final particle images (no.) | 89,435 | | | | |
| Map sharpening B factor ($Å^2$) | -38 | -29 | -32.5 | -35 | |
| Map resolution (FSC=0.143) (Å) | 2.40 | 2.20 | 2.41 | 2.47 | |
| Map resolution D99 | | | | | 2.40 |
| Map angular accuracy (°) | 0.266 | 0.255 | 0.438 | 0.434 | |
| Map translational accuracy(Å) | 0.191 | 0.210 | 0.283 | 0.281 | |
| Map resolution range (Å) | 2.12-5.63 | 2.09-4.46 | 2.22-6.20 | 2.30-6.01 | |
| Final map sampling (Å/pix) | 0.731 | | | | |
| **Model Refinement** | | | | | |
| Initial model used | | | | | |
| Model resolution (FSC=0.5) (Å) | | | | | 2.4 |
| Model composition | | | | | |
| Nonhydrogen atoms | | | | | 65494 |
| Protein residues | | | | | 7788 |
| Waters | | | | | 1415 |
| Ligands | | | | | 44 |
| B factors mean ($Å^2$) | | | | | |
| Protein | | | | | 47.04 |
| Ligand | | | | | 47.72 |
| Water | | | | | 40.75 |
| RMSD Bond lengths (Å) | | | | | 0.006 |
| RMSD Bond angles (°) | | | | | 1.015 |
| Validation | | | | | |
| MolProbity score | | | | | 1.26 |
| Clashscore | | | | | 2.84 |
| Poor rotamers (%) | | | | | 1.29 |
| EMRinger score | 4.84 | | | | 5.70 |
| Ramachandran plot | | | | | |
| Favored (%) | | | | | 97.52 |
| Allowed (%) | | | | | 2.46 |
| Disallowed (%) | | | | | 0.01 |
| Z-score (whole) | | | | | 0.66 |
| Map-model correlation $CC_{mask}$ | | | | | 0.90 |

**Supplementary Table 12. Cryo-EM data collection, refinement and validation statistics for slack-1092-i**

| | Slack-1092-ii | | | | |
|---|---|---|---|---|---|
| | Consensus | Hydrophilic domain | Proximal hydrophobic domain | Distal hydrophobic domain | Composite |
| PDB code | | | | | PDB 7R4G |
| EMDB code | EMD-14308 | EMD-14309 | EMD-14310 | EMD-14311 | EMD-14307 |
| **Data collection and processing** | | | | | |
| Magnification (nominal) | 81,000 x | | | | |
| Voltage (kV) | 300 | | | | |
| Electron exposure ($e^{-}/Å^2$) | 40 | | | | |
| Defocus range (μm) | -1.0 to -2.4 | | | | |
| Calibrated Pixel size (Å) | 1.072 | | | | |
| Symmetry imposed | none | | | | |
| Initial particle images (no.) | 907,090 | | | | |
| Final particle images (no.) | 40,154 | | | | |
| Map sharpening B factor ($Å^2$) | -36 | -24 | -29 | -29 | |
| Map resolution (FSC=0.143) (Å) | 2.61 | 2.37 | 2.62 | 2.68 | |
| Map resolution D99 | | | | | 2.50 |
| Map angular accuracy (°) | 0.257 | 0.252 | 0.515 | 0.435 | |
| Map translational accuracy(Å) | 0.203 | 0.220 | 0.332 | 0.298 | |
| Map resolution range (Å) | 2.25-7.20 | 2.19-5.31 | 2.40-8.11 | 2.50-8.04 | |
| Final map sampling (Å/pix) | 0.731 | | | | |
| **Model Refinement** | | | | | |
| Initial model used | | | | | |
| Model resolution (FSC=0.5) (Å) | | | | | 2.5 |
| Model composition<br>Nonhydrogen atoms<br>Protein residues<br>Waters<br>Ligands | | | | | <br>65153<br>7802<br>851<br>46 |
| B factors mean ($Å^2$)<br>Protein<br>Ligand<br>Water | | | | | <br>50.13<br>52.43<br>41.12 |
| RMSD Bond lengths (Å)<br>RMSD Bond angles (°) | | | | | 0.007<br>1.026 |
| Validation<br>MolProbity score<br>Clashscore<br>Poor rotamers (%) | | | | | <br>1.39<br>4.13<br>0.03 |
| EMRinger score | 4.52 | | | | 5.55 |
| Ramachandran plot<br>Favored (%)<br>Allowed (%)<br>Disallowed (%)<br>Z-score (whole) | | | | | 96.80<br>3.19<br>0.04<br>0.01 |
| Map-model correlation $CC_{mask}$ | | | | | 0.89 |

**Supplementary Table 13. Cryo-EM data collection, refinement and validation statistics for slack-1092-ii**

| | Active inhibitor-free | | | | |
|---|---|---|---|---|---|
| | Consensus | Hydrophilic domain | Proximal hydrophobic domain | Distal hydrophobic domain | Composite |
| PDB code | PDB-7QSD | | | | |
| EMDB code | EMD-14127 | EMD-14129 | EMD-14130 | EMD-14131 | EMD-14128 |
| **Data collection and processing** | | | | | |
| Magnification (nominal) | 130,000 x | | | | |
| Voltage (kV) | 300 | | | | |
| Electron exposure ($e^{-}/Å^2$) | 50 | | | | |
| Defocus range (μm) | -1.5 to -3.1 | | | | |
| Calibrated Pixel size (Å) | 1.056 | | | | |
| Symmetry imposed | none | | | | |
| Initial particle images (no.) | 56,908 | | | | |
| Final particle images (no.) | 18,231 | | | | |
| Map sharpening B factor ($Å^2$) | -33 | -33 | -26 | -35 | |
| Map resolution<br>(FSC=0.143) (Å) | 3.15 | 3.09 | 3.28 | 3.37 | |
| Map resolution D99 | | | | | 2.80 |
| Map angular accuracy (°) | 0.368 | 0.412 | 0.633 | 0.593 | |
| Map translational accuracy(Å) | 0.280 | 0.358 | 0.442 | 0.465 | |
| Map resolution range (Å) | 2.77-7.28 | 2.75-15.97 | 2.97-11.7 | 2.99-17.77 | |
| Final map sampling (Å/pix) | 1.056 | | | | |
| **Model Refinement** | | | | | |
| Initial model used | | | | | |
| Model resolution<br>(FSC=0.5) (Å) | 3.0 | | | | |
| Model composition<br>Nonhydrogen atoms<br>Protein residues<br>Waters<br>Ligands | 67258<br>8204<br>41 | | | | |
| B factors mean ($Å^2$)<br>Protein<br>Ligand<br>Water | 45.70<br>45.62 | | | | |
| RMSD Bond lengths (Å)<br>RMSD Bond angles (°) | 0.007<br>1.019 | | | | |
| Validation<br>MolProbity score<br>Clashscore<br>Poor rotamers (%) | 1.70<br>5.69<br>0 | | | | |
| EMRinger score | 4.09 (Local) | | | | 4.06 |
| Ramachandran plot<br>Favored (%)<br>Allowed (%)<br>Disallowed (%)<br>Z-score (whole) | 94.22<br>5.70<br>0.04<br>0.64 | | | | |
| Map-model correlation $CC_{mask}$ | 0.82 | | | | |

**Supplementary Table 14. Cryo-EM data collection, refinement and validation statistics for Active inhibitor-free**

| | Slack inhibitor-free |
|---|---|
| | Consensus |
| EMDB code | EMD-14126 |
| **Data collection and processing** | |
| Magnification | 130,000 x |
| Voltage (kV) | 300 |
| Electron exposure ($e^{-}/Å^2$) | 50 |
| Defocus range (μm) | -1.5 to -3.1 |
| Calibrated Pixel size (Å) | 1.056 |
| Symmetry imposed | none |
| Initial particle images (no.) | 56,908 |
| Final particle images (no.) | 14,437 |
| Map sharpening B factor ($Å^2$) | -42 |
| Map resolution (FSC = 0.143) (Å) | 3.49 |
| Map angular accuracy (°) | 0.468 |
| Map translational accuracy (Å) | 0.319 |
| Map resolution range (Å) | 3.05-9.8 |
| Final map sampling (Å/pix) | 1.056 |

**Supplementary Table 15. Cryo-EM data collection, refinement and validation statistics for Slack inhibitor-free.**

### References and Notes


1. J. N. Blaza, R. Serreli, A. J. Y. Jones, K. Mohammed, J. Hirst, Kinetic evidence against partitioning of the ubiquinone pool and the catalytic relevance of respiratory-chain supercomplexes. *Proc. Natl. Acad. Sci. U. S. A.* **111**, 15735–15740 (2014).
2. A. N. A. Agip, J. N. Blaza, H. R. Bridges, C. Viscomi, S. Rawson, S. P. Muench, J. Hirst, Cryo-em structures of complex i from mouse heart mitochondria in two biochemically defined states. *Nat. Struct. Mol. Biol.* **25**, 548–556 (2018).
3. M. S. Sharpley, R. J. Shannon, F. Draghi, J. Hirst, Interactions between phospholipids and NADH:ubiquinone oxidoreductase (complex I) from bovine mitochondrial. *Biochemistry*. **45**, 241–248 (2006).
4. A. J. Y. Jones, J. N. Blaza, H. R. Bridges, B. May, A. L. Moore, J. Hirst, A Self-Assembled Respiratory Chain that Catalyzes NADH Oxidation by Ubiquinone-10 Cycling between ComplexI and the Alternative Oxidase. *Angew. Chemie - Int. Ed.* **55**, 728–731 (2016).
5. A. J. Y. Jones, J. N. Blaza, H. R. Bridges, B. May, A. L. Moore, J. Hirst, A Self-Assembled Respiratory Chain that Catalyzes NADH Oxidation by Ubiquinone-10 Cycling between Complex I and the Alternative Oxidase. *Angew. Chemie*. **128**, 738–741 (2016).
6. T. B. Walpole, D. N. Palmer, H. Jiang, S. Ding, I. M. Fearnley, J. E. Walker, Conservation of Complete Trimethylation of Lysine-43 in the Rotor Ring of c-Subunits of Metazoan Adenosine Triphosphate (ATP) Synthases*. *Mol. Cell. Proteomics*. **14**, 828–840 (2015).
7. J. Carroll, I. M. Fearnley, R. J. Shannon, J. Hirst, J. E. Walker, Analysis of the subunit composition of complex I from bovine heart mitochondria. *Mol. Cell. Proteomics*. **2**, 117–126 (2003).
8. J. N. Blaza, K. R. Vinothkumar, J. Hirst, Structure of the Deactive State of Mammalian Respiratory Complex I. *Structure*. **26**, 312-319.e3 (2018).
9. J. Zivanov, T. Nakane, B. O. Forsberg, D. Kimanius, W. J. H. Hagen, E. Lindahl, S. H. W. Scheres, New tools for automated high-resolution cryo-EM structure determination in RELION-3. *Elife*. **7**, e42166 (2018).
10. K. Zhang, Gctf: Real-time CTF determination and correction. *J. Struct. Biol.* **193**, 1–12 (2016).
11. J. Zhu, K. R. Vinothkumar, J. Hirst, Structure of mammalian respiratory complex i. *Nature*. **536**, 354–358 (2016).
12. I. Chung, J. J. Wright, H. R. Bridges, B. S. Ivanov, O. Biner, C. S. Pereira, G. M. Arantes, J. Hirst, Cryo-EM structures define ubiquinone-10 binding to mitochondrial complex I and conformational transitions accompanying Q-site occupancy. *Nat. Commun.* **13**, 2758 (2022).
13. J. Zivanov, T. Nakane, S. H. W. Scheres, Estimation of high-order aberrations and anisotropic magnification from cryo-EM data sets in RELION-3.1. *IUCrJ*. **7**, 253–267 (2020).
14. A. Rohou, N. Grigorieff, CTFFIND4: Fast and accurate defocus estimation from electron micrographs. *J. Struct. Biol.* **192**, 216–221 (2015).
15. E. F. Pettersen, T. D. Goddard, C. C. Huang, G. S. Couch, D. M. Greenblatt, E. C. Meng, T. E. Ferrin, UCSF Chimera - A visualization system for exploratory research and analysis. *J. Comput. Chem.* **25**, 1605–1612 (2004).
16. P. Emsley, K. Cowtan, Coot: Model-building tools for molecular graphics. *Acta*

*Crystallogr. Sect. D Biol. Crystallogr.* **60**, 2126–2132 (2004).
17. P. V. Afonine, B. K. Poon, R. J. Read, O. V. Sobolev, T. C. Terwilliger, A. Urzhumtsev, P. D. Adams, Real-space refinement in PHENIX for cryo-EM and crystallography. *Acta Crystallogr. Sect. D Struct. Biol.* **74**, 531–544 (2018).
18. J. Gu, T. Liu, R. Guo, L. Zhang, M. Yang, The coupling mechanism of mammalian mitochondrial complex I. *Nat. Struct. Mol. Biol.* **29**, 172–182 (2022).
19. D. Kampjut, L. A. Sazanov, The coupling mechanism of mammalian respiratory complex I. *Science (80-. ).* **370**, eabc4209 (2020).
20. Z. Yin, N. Burger, D. Kula-Alwar, D. Aksentijević, H. R. Bridges, H. A. Prag, D. N. Grba, C. Viscomi, A. M. James, A. Mottahedin, T. Krieg, M. P. Murphy, J. Hirst, Structural basis for a complex I mutation that blocks pathological ROS production. *Nat. Commun.* **12**, 707 (2021).
21. D. C. Bas, D. M. Rogers, J. H. Jensen, Very fast prediction and rationalization of pK. *Proteins* (2008).
22. K. Parey, O. Haapanen, V. Sharma, H. Köfeler, T. Züllig, S. Prinz, K. Siegmund, I. Wittig, D. J. Mills, J. Vonck, W. Kühlbrandt, V. Zickermann, High-resolution cryo-EM structures of respiratory complex I: Mechanism, assembly, and disease. *Sci. Adv.* **5**, eaax9484 (2019).
23. J. M. Berrisford, L. A. Sazanov, Structure of Respiratory Complex I. *Encycl. Biol. Chem. Second Ed.* **494**, 341–346 (2013).
24. E. F. Pettersen, T. D. Goddard, C. C. Huang, E. C. Meng, G. S. Couch, T. I. Croll, J. H. Morris, T. E. Ferrin, UCSF ChimeraX: Structure visualization for researchers, educators, and developers. *Protein Sci.* **30**, 70–82 (2021).
25. W. Tian, C. Chen, X. Lei, J. Zhao, J. Liang, CASTp 3.0: Computed atlas of surface topography of proteins. *Nucleic Acids Res.* **46**, W363–W367 (2018).
